# Mapping the Phenotypic Landscape of Beta-lactam Resistance in Streptococcus pneumoniae

**DOI:** 10.1101/2025.09.12.675231

**Authors:** Andrew J. Balmer, Gemma G. R. Murray, Stephanie W. Lo, Olivier Restif, Lucy A. Weinert

## Abstract

Beta-lactam resistance in *Streptococcus pneumoniae* is typically analysed one antibiotic at a time, often using resistant/susceptible classifications. This obscures the fact that minimum inhibitory concentrations (MICs) in this species are continuous, vary across beta-lactam subclasses in correlated but non-identical ways, and reflect complex genetic variation in underlying penicillin-binding proteins (PBPs). To address this, we develop a framework for analysing multivariate resistance phenotypes, which we term ’Antimicrobial Resistance Cartography’, and apply it to 3,628 invasive pneumococcal isolates from the U.S. Active Bacterial Core surveillance programme with MICs measured for six beta-lactams. We demonstrate how this approach simplifies visualisation of resistance data across multiple drugs, resolves common issues such as missing or censored values, and identifies genetic mutations driving correlated resistance changes across multiple beta-lactam antibiotics. We characterise penicillin-binding protein (PBP) substitutions associated with increases in minimum inhibitory concentration to beta-lactam subclasses, revealing subtle differences in their correlated phenotypic effects, and identifying potential constraints on resistance evolution across genetic backgrounds. Moreover, we find pneumococcal lineages harbour distinct combinations of PBP substitutions which give rise to lineage-specific multivariate resistance profiles. Together, these results establish a multivariate view of pneumococcal beta-lactam resistance, revealing how combinations of PBP substitutions work together to generate distinct resistance phenotypes across related antibiotics.

**Significance Statement:** Beta-lactam resistance in *Streptococcus pneumoniae* is typically analysed one antibiotic at a time, despite susceptibility being a continuous phenotype that covaries across multiple drugs simultaneously. Current analytical methods for resistance datasets overwhelmingly focus on genotypes, while tools to study beta-lactam phenotypes, the direct targets of selection and clinical intervention, remain limited. To address this, we developed a dimensionality reduction framework that maps complex susceptibility profiles into simple visualisations, allowing resistance to be studied as a continuous, multivariate trait. Applying this approach to 3,628 pneumococcal isolates revealed lineage-specific resistance profiles and identified candidate PBP substitutions associated with shared, subclass-specific, and contrasting effects across beta-lactam antibiotics. By preserving quantitative variation and correlations between drugs, this approach shows how multivariate phenotypic methods can improve interpretation of pneumococcal beta-lactam resistance and prioritise candidate variants for experimental validation.

## Introduction

*Streptococcus pneumoniae* is a major human pathogen responsible for an estimated 1.2 million infections and 7000 deaths annually in the USA alone (1–4). Beta-lactams remain central to the treatment of pneumococcal disease, but the extent of their use has driven widespread resistance – in some cases repeatedly, over short timeframes and across diverse populations (3,5). Because susceptibility phenotypes are the direct targets of both clinical intervention and natural selection, understanding the phenotypic evolution of beta-lactam resistance, and its effects on pneumococcal population dynamics, has become a major focus of research (6–9).

Consequently, over the past two decades, large-scale surveillance programs have amassed collections of thousands of *S. pneumoniae* isolates, each tested for susceptibility to multiple beta-lactams using phenotypic assays such as minimum inhibitory concentration (MIC) (4,8,10). Two observations repeatedly emerge from these studies: I) susceptibility profiles can vary widely in *S. pneumoniae*–even within defined resistance categories, where MICs span multiple dilution steps (e.g. 4, 8, 16, or 32 μg/mL), and II) susceptibility phenotypes across beta-lactam drugs are typically correlated (11–14). This covariation can arise through several mechanisms: in some cases, one genotype confers resistance to multiple antibiotics; in others, epidemiological factors, such as sequential treatment with separate beta-lactams, generates co-resistance (13). Understanding how these correlations arise and persist is crucial, because they could accelerate drug resistance across multiple beta-lactams simultaneously, restricting the efficacy of the entire drug class (3,11). There is therefore a need for methods that can study these patterns of phenotypic variation and covariation and test hypotheses on their causes.

However, to develop these tools, three methodological challenges must be addressed. Firstly, beta-lactam MIC values in *S. pneumoniae* are often reduced to resistant/susceptible categories based on clinical or epidemiological cut-offs (6,15,16). While useful for guiding treatment and surveillance, these simplifications discard quantitative variation within and below resistance classes that is crucial in clinical, epidemiological and evolutionary contexts. For example, gradual increases in beta-lactam MIC can reduce treatment efficacy, prolong infections, and confer selective advantages during low-level antibiotic exposure–common in both carriage sites and environmental reservoirs (14,17–21). Secondly, MICs are typically analysed separately for each drug, despite the underlying resistance mechanisms being known to affect multiple beta-lactams simultaneously (6,22–24). Clinically, this is important for selecting effective therapies, but it is also important from an evolutionary perspective, because genetic changes that affect multiple phenotypes can shape subsequent trajectories (13). For instance, if a single mechanism confers resistance to multiple beta-lactams, selection for resistance to any of those drugs may propagate resistance to others (25,26). Finally, MIC datasets for *S. pneumoniae* frequently contain missing or heterogeneous measurements, such as when values fall outside the dilution range of the assay, or when multiple susceptibility-testing methods are combined within a single collection (27–29). These issues often necessitate separate analyses per drug, complicating clinical surveillance and reducing statistical power to identify subtle, but important evolutionary changes over time (30,31). Existing multivariate approaches (pairwise correlations, categorical classification rules, multi-trait GWAS) address some of these problems, but are limited in their ability to capture fine-scale variation in continuous MIC data, identify covarying effects, and provide an intuitive representation of high-dimensional resistance in relation to metadata (32–34). Addressing these limitations therefore requires novel methods that can visualise and simplify these complex phenotypic patterns and understand how they relate to the genetic and epidemiological context of *S. pneumoniae* (3,35).

These challenges are particularly relevant in *S. pneumoniae* because of the complex genetic mechanisms underlying beta-lactam resistance. In streptococci, beta-lactams bind to the active site of the penicillin-binding proteins (PBPs), inhibiting their role in peptidoglycan cell wall biosynthesis (22). Resistance primarily arises through the modification of three PBPs—PBP1A, PBP2B, and PBP2X—where changes in the transpeptidase domains reduce antibiotic binding affinity (8). The variants within these proteins that reduce antibiotic binding (i.e. causal variants) are contained within highly variable ‘mosaic’ blocks, formed via recombination with other streptococci species (6,36,37). However, within these blocks, many substitutions are merely hitchhikers (i.e. variants that do not directly affect resistance), and others only increase MIC in combination or alter the fitness effects of primary resistance mutations (38–40). While existing GWAS and machine learning methods reliably link changes in the PBPs with non-susceptible phenotypes, precisely identifying the individual substitutions associated with changes in MIC remains challenging, particularly when GWAS analyses each antibiotic separately or as binary resistant/susceptible traits, as this limits the detection of fine-scale, correlated effects across drugs (6,30,34). Consequently, there has been limited overlap between methods in which PBP substitutions are identified as associated with non-susceptible phenotypes (3). Applying novel methods which could better identify important substitutions would improve resistance predictions, inform diagnostic assays, and strengthen surveillance of emerging resistance.

However, to address these issues, we need tools which can account for the fact that beta-lactam phenotypes–and their underlying genetic mechanism–are both continuous and multivariate. While all beta-lactams target PBPs, different subclasses bind to each PBP with distinct binding affinities–for example, penicillins (e.g. penicillin, amoxicillin) primarily bind to PBP2X and PBP2B, whereas cephalosporins bind predominantly to PBP2X and 1A (24,41,42). Because of this, a single PBP mutation can affect susceptibility to multiple beta-lactams in different ways (23). Multiple PBP alterations are required for high MICs, each contributing modest effects to one or more drugs, but cumulatively leading to clinical resistance across several drugs (36–38). Although subclasses show positively correlated MICs (i.e. partial cross-resistance), these differences in binding affinity mean beta-lactam resistance is not a single phenotype, but rather a complex, interrelated set of traits. Together, these features make *S. pneumoniae* an ideal model system for developing methods to visualise multivariate drug resistance phenotypes, and to use them to interpret how mutations collectively confer high-level resistance. These tools would then allow us to investigate whether phenotypes differ across phylogenetic lineages (i.e. Multilocus Sequence Types - MLST) (35,43).

To develop this toolkit, we propose an approach inspired by ‘Antigenic Cartography’. Originally developed to visualise antigenic evolution in H3N2 influenza, antigenic cartography has enabled the precise tracking of antigenic changes in viral pathogens–guiding vaccine design and improving understanding of immune responses to evolving pathogens (44–47). The tool is based on a dimensionality reduction method called multidimensional scaling (MDS)–a technique that visualises high-dimensional data by placing isolates in a low-dimensional phenotypic ‘map’–where distances between points represent differences in measured assay data. This enables cartography to summarise complex antigenic data in a concise, visual format, reduce measurement error, and improve interpretation of phenotypic differences among viral strains. Since its assumptions do not restrict its application to antigenic phenotypes, we adapted it to map antimicrobial resistance phenotypes, using MIC profiles for multiple antibiotics.

As a proof of concept, we apply this method to a large collection of clinical isolates of *S. pneumoniae* for which joint phenotypic and genotypic data were available: focusing on MIC values against six beta-lactams and amino acid sequences of the transpeptidase regions of PBPs. We have three objectives: I) quantitatively visualise and interpret continuous covariation in susceptibility across six beta-lactams; II) examine how these phenotypes associate with specific phylogenetic lineages (MLSTs); and III) identify and rank PBP amino acid substitutions according to the strength and consistency of their association with increased beta-lactam MIC. In addition, we demonstrate the approach mitigates common experimental issues – measurement error, censored MIC values, missing data – and increases the resolution of genotype-phenotype associations, including the detection of epistatic effects for experimental follow-up. Together, we show the method allows for a more quantitative comparison of MIC values, which improves visualisation and experimental-error handling, allows comparison of multivariate distance metrics, and helps identify correlations and constraints on multivariate phenotypes.

## Methods

### Isolates and metadata

We used a published dataset of 4,309 invasive pneumococcal isolates, collected in the United States through the Active Bacterial Core surveillance (ABC) programme between 1998 and 2015 (8,48). The ABC project is an active, population-based surveillance system for invasive bacterial disease, run by the Centers for Disease Control and Prevention (CDC) Emerging Infectious Disease programme (4). As part of this initiative, microbiology laboratories in participating hospitals identify cases of invasive pneumococcal disease, conduct laboratory testing, and share data with the programme. Isolates were defined as pneumococcal strains isolated from a normally sterile site and were collected from clinical cases within ten U.S. states, each representing different geographic regions.

MICs were previously measured for six beta-lactam antibiotics using two-fold microbroth dilution methods by the ABC program (Supplementary Table S1). For our analysis, we focused on 3,628 isolates with MIC available for 6 beta-lactams, of three subclasses: penicillin, amoxicillin (penicillins), ceftriaxone, cefotaxime, cefuroxime (cephalosporins), and meropenem (a carbapenem). Whole genome sequencing for all isolates was performed using the Illumina HiSeq or MiSeq platforms (8,48), and sequencing data were used to determine capsular serotype, MLST, and transpeptidase sequences for penicillin binding proteins PBP1A, PBP2B and PBP2X. Additional metadata—including date, geographical location, and clinical symptoms—are publicly available, but were not used here. Our analysis therefore reflects invasive pneumococcal isolates from clinical disease in the United States between 1998 and 2015, and conclusions are accordingly limited to this surveillance context and to this drug class.

### Phenotype map construction

Phenotype maps visualise a Euclidean distance matrix derived from MIC data. To run iterative multidimensional scaling (SMACOF algorithm) using MIC data, MIC values were log₂ transformed to approximate a standard distribution (49,50). Titres for the six drugs were then transformed into a multivariate distance matrix by calculating pairwise Euclidean distances between isolates (*D_i_*_,*j*_), therefore representing differences across all drugs simultaneously. Given this set of target distances, MDS finds a low-dimensional representation of those distances with minimal error (49,50). Distances in the representation are determined by minimising the differences between map distances (*d_i_*_,*j*_) and table distances *(D_i_*_,*j*_) for each pair of isolates i and j, using an error function called stress (*S*):

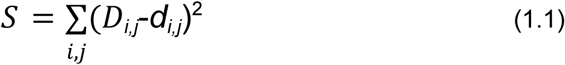

Here, ‘stress’ is the sum of the squared residuals between points on the map and their measured distances. The MDS algorithm searches for a minimum stress value by iteratively shifting the positions of isolates on the map. As MDS can be applied to either metric or rank data, both numeric and censored MIC values can be represented (Supplementary Section S1.6). We also developed “genetic maps”, which visualise genetic differences among isolates using a Hamming distance matrix based on an amino acid sequence alignment (Supplementary Figure S8) (44). We evaluated the goodness-of-fit of these maps using cross-validation, confidence intervals and dimensionality testing (Supplementary Section S1).

### Genotype-phenotype association methods

To elucidate the molecular basis of multivariate phenotypic variation, we focused on amino acid substitutions in the transpeptidase regions of PBPs, given their known role in beta-lactam resistance. We used four methods to identify substitutions associated with changes in position on the phenotype map (Supplementary Section S2.5-9), each of which were originally developed to identify substitutions underlying antigenic change in H3N2 influenza (44,51). First, we identified pairs of isolates that varied by a single amino acid and calculated the phenotypic distances between them. Second, we used clustering algorithms to partition the phenotype map into distinct regions and identify substitutions associated with these regions. Third, we used multivariate linear mixed models (mvLMM) to test associations between amino acid substitutions and isolate position on the phenotype map, and estimate the phenotypic effect sizes of each substitution (52,53). Lastly, we extended the mvLMM to test for interaction effects between PBP substitutions, while including the individual markers as covariates (see Supplementary Section S2.9). To enhance statistical power relative to conventional genome-wide association approaches, we used several methods to reduce the correction for multiple testing burden, such as using amino acid changes instead of nucleotide changes, and restricting associations to PBP transpeptidase sequences (Supplementary Section S2.10) (51).

## Results

### Antimicrobial Resistance Cartography of Streptococcus pneumoniae

We analysed 3,628 invasive pneumococcal isolates from the Active Bacterial Core (ABC) surveillance program, with MICs measured for six beta-lactam antibiotics using two-fold microbroth dilution. Across these isolates, most exhibited low MIC values for all six antibiotics, reflecting a high level of susceptibility to the drugs. Despite this, MIC values varied considerably across isolates, with penicillin MICs ranging from ≤0.03 to 8μg/mL. MIC values were positively correlated across all beta-lactams, although the strength of these correlations varied, particularly among drugs with a smaller range of MIC values e.g. cefuroxime (Pearson correlation coefficients = 0.77-0.96; Figure 1A).

**Figure 1.**
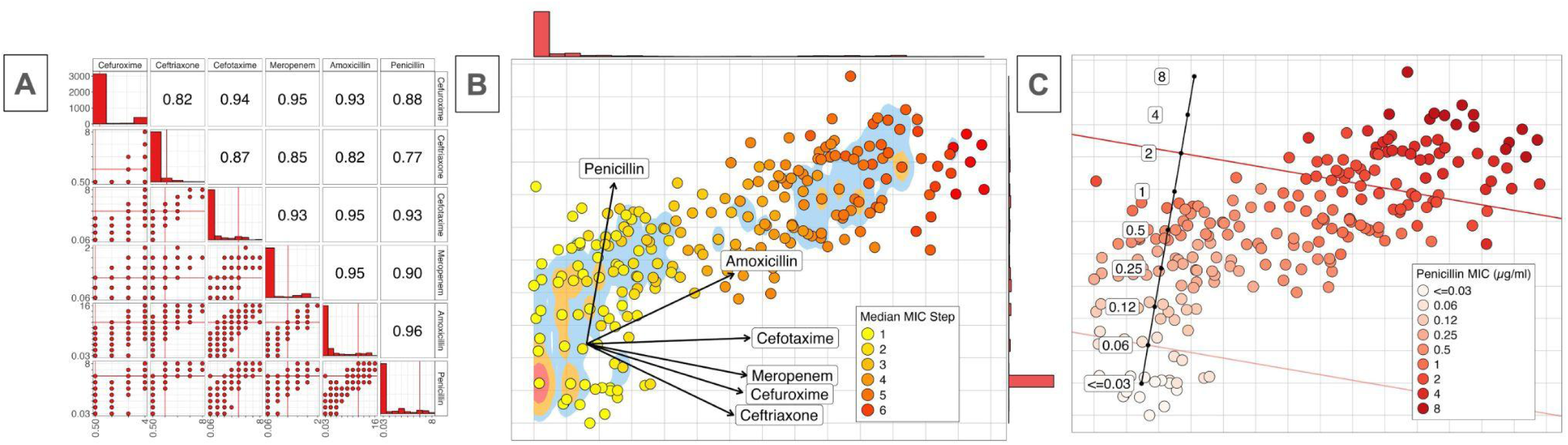
A) MIC measurements for 3628 isolates of *S. pneumoniae* across six beta-lactam drugs, visualised using a pairwise correlation plot (Pearson). Axes represent the dilution step in a MIC series for the 6 beta-lactam drugs, where red lines indicate clinical breakpoints for each drug. The upper triangle shows the pairwise correlation coefficients for each pair of drugs. Although beta-lactam MIC data is often visualised using pairwise correlation plots such as these, it is difficult to interpret susceptibility profiles across multiple drugs for any given isolate. For example, isolates might have varying resistance profiles e. g. S-S-S-R-R-R, or S-R-R-R-R-R, but it is not possible to interpret these nor their relative proportions. B) Antimicrobial Cartography map for the same 3628 isolates, coloured by median MIC dilution step (i.e. 1 = the lowest concentration measured for a given drug). Points represent isolates, while the two primary axes of the map represent phenotypic distance in MIC units. One grid space corresponds to one average two-fold dilution step across the six beta-lactam assays. As multiple isolates can occupy the same point in the map, density distributions (red area – 50%, orange - 80% and blue, 95% of isolates) and marginal histograms are used to highlight the distribution of isolates. Biplot axes for each axis indicate the direction of increasing MIC for the drugs. These axes indicate penicillin MIC strongly correlates with the vertical axis, amoxicillin correlates with both the vertical and horizontal axes, and the remaining drugs primarily align horizontally. C) Biplot axes calibrated to show the MIC distribution of isolates on the map, where isolates in line with a value on the scale have that MIC value. Isolates are coloured by Penicillin MIC. Clinically relevant cut-offs can be included to aid visualisation (red and pink lines indicate resistance cut-off for Penicillin in non-meningitis and meningitis infections respectively - 2μg/mL & 0.06μg/mL).

We visualised multivariate resistance profiles among the isolates using multidimensional scaling on a Euclidean distance matrix derived from log₂ transformed MIC data (Figure 1A). On the resulting phenotype ‘map’, each isolate is represented as a point, with distances between them reflecting differences in MIC values across the six antibiotics (Figure 1B). Instead of simplifying data into binary resistant/susceptible categories, this approach provides an intuitive representation of the full distribution of multivariate phenotypes, capturing both fine-scale MIC variation between isolates and the underlying correlations in resistance profiles across drugs. Most isolates displayed low MIC values for all antibiotics, and these are represented by the cluster in the lower-left region of the figure. The map also shows substantial variability in MIC profiles among the remaining isolates. For example, isolates with intermediate penicillin MIC (0.12–1μg/mL) varied widely in their MIC values for other drugs, sometimes differing by up to six two-fold dilution steps, yet remaining below the clinical breakpoint for penicillin (2μg/mL) (Figure 1C). Additionally, a subset of isolates in the top-right of the map had uniformly high penicillin MIC but varied in MIC to cephalosporins and carbapenems (Figure 1C, Supplementary Figure S2). The overall diagonal distribution of isolates on the map reflects the strong positive correlations in MIC values across beta-lactams, but the two dimensions required to represent the phenotypes demonstrate that correlations between different drugs are not perfectly linear.

To evaluate the accuracy of the map, we calculated the pairwise Euclidean distances between isolates in the original six-dimensional MIC space and compared them to their relative distances on the 2D map. This allowed us to quantify how well the 2D projection preserves the original multivariate relationships. Fewer than 0.3% of pairwise distances had a discrepancy greater than one log₂ dilution between their true multivariate MIC distance and their map-based distance (Table 1, Supplementary Figure S2). This indicates that a two-dimensional map is an accurate representation of this set of MIC values. The map also displayed high stability across a range of resampling, dimensionality, and cross-validation tests (Supplementary Sections S1.7-1.10).

**Table 1.** Goodness-of-fit metrics across MDS dimensionalities for the phenotype map. Projection error refers to the mean change in each unique phenotype’s position when adding an additional dimension to the map. Values therefore reflect the relative mean decrease in error when moving from 1D to 2D, 2D to 3D, etc. A higher value indicates a larger gain in information from adding an additional dimension. Note the sharp reduction in overall stress when moving from a 1D to 2D map (55.5%).

| Dimension | Decrease in stress relative to a 1D map (%) | Pairwise error (MIC units) |  |  |  | Projection Error of unique phenotypes (MIC units) |
| --- | --- | --- | --- | --- | --- | --- |
|  |  | Between all isolates |  |  |  |  |
|  |  | Mean | SD | Error 1-2 (%) | Error >2 (%) |  |
| 1D | NA | 0.105 | 0.234 | 1.323 | 0.186 | NA |
| 2D | 55.534 | 0.035 | 0.108 | 0.209 | 0.01 | 1.28 |
| 3D | 80.961 | 0.013 | 0.047 | 0.012 | 0 | 0.406 |
| 4D | 88.025 | 0.006 | 0.03 | 0.004 | 0 | 0.09 |
| 5D | 93.534 | 0.003 | 0.016 | 0 | 0 | 0.05 |

Although isolates could, in theory, be positioned anywhere throughout the multidimensional space, they predominantly aligned along a near-linear straight line. Nevertheless, goodness-of-fit comparisons confirmed that a two-dimensional map provided a significantly better representation of fine-scale differences in MIC profiles among isolates than a one-dimensional projection, as indicated by the large decrease in error when moving from a one- to two-dimensional map (Table 1, Supplementary Sections S1.5 and S1.12). Notably, although mean pairwise error values appear low across all dimensions, this is due to the fact that many isolates exhibited uniformly low, censored MIC values (i.e. below the dilution range of the assay), and were consequently well fit in a single dimension. However, phenotypes with elevated MIC contributed proportionally much higher stress when represented in a single dimension, as indicated by the higher pairwise error and decrease in stress values. This was further supported by a projection error analysis, which quantified the average change in position of each unique phenotype when adding additional dimensions, showing a large decrease in error when moving from one- to two-dimensions (1.28 MIC units).

While higher-dimensional projections further reduced error (Table 1), improvements beyond two dimensions were more modest. Moving from two to three dimensions changed the projected positions of unique phenotypes by a mean of 0.406 MIC units—less than the expected error of MIC testing—suggesting that two-dimensional maps adequately capture the key phenotypic patterns in this dataset (54). Moreover, as 2D maps performed better in cross-validation analyses, and isolate distributions on the 2D and 3D maps remained directly comparable, the 2D maps are presented throughout this manuscript.

### Robustness of AMR Cartography in addressing common assay issues

Theoretically, by positioning isolates using data from multiple drugs, the map can use the associations between them to more accurately infer MIC values, even when underlying data is missing or inaccurate. In principle, this means the map can offer greater accuracy than using MIC values for any single drug individually. We tested whether this was the case by assessing whether the map could alleviate common measurement issues in resistance surveillance datasets, such as missing values, experimental error, and combining multiple assay methods.

To test robustness in handling missing data, we randomly removed 10% of MIC values, reconstructed the map with the remaining data, and predicted the missing MICs using the relative distances between points (Supplementary Figure S4). Across 100 iterations, 86.8% of predictions for missing MIC values were within a single MIC dilution of the true value (mean error = 0.467)—the range of error expected if isolates were remeasured experimentally (15,16,55)—demonstrating the robustness of the map in imputing missing data (Table 2, Supplementary Figure S4). Prediction error was low across the distribution of the map, though typically increased with the number of missing MIC values for a given isolate. This was because isolates with additional missing values had less information on where to position them, although error still typically remained below 2 MIC units (Supplementary Figure S4).

**Table 2.**
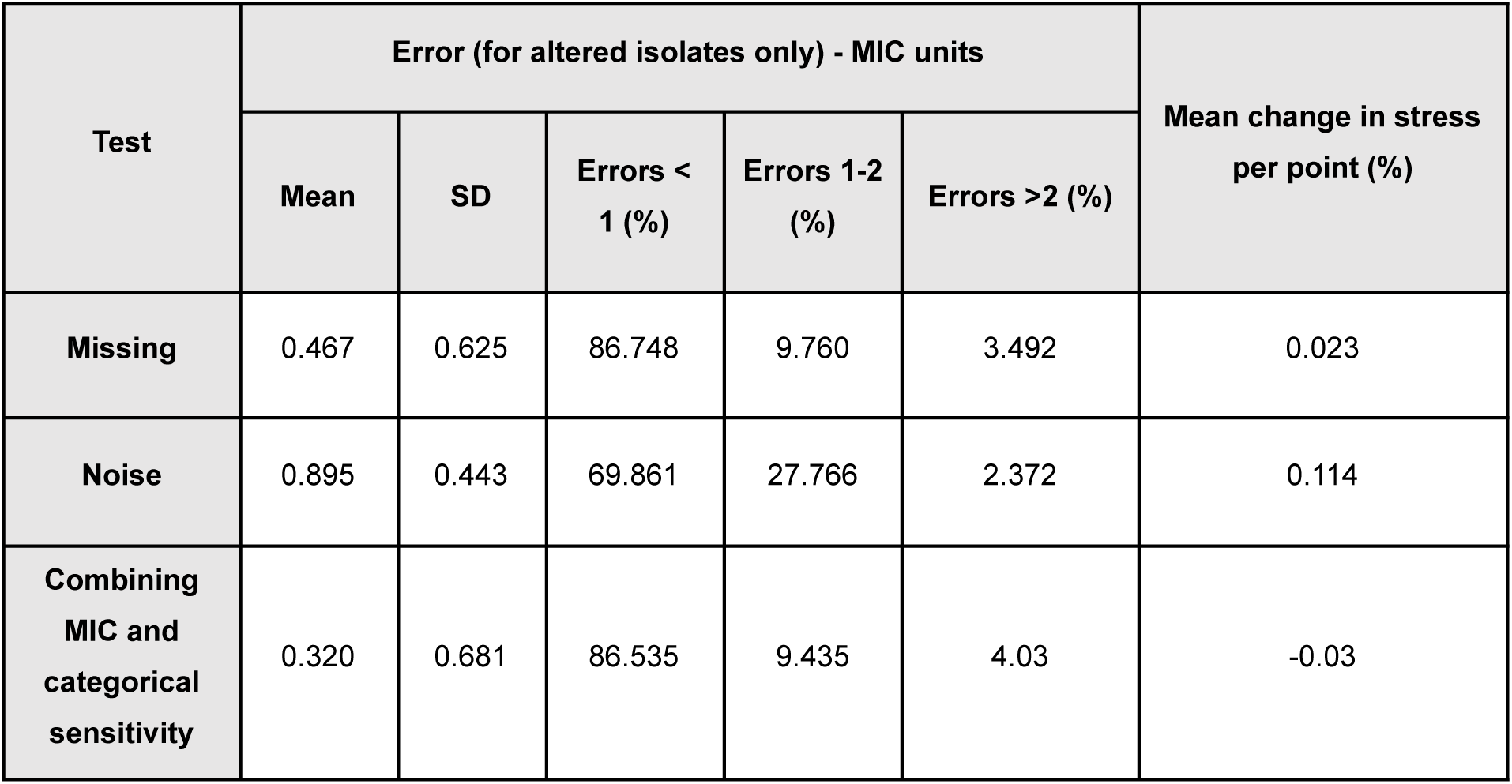
Results of error-prediction analyses for the phenotype map. Each row summarises results from 100 datasets, each adjusted by either removing values (Missing), adding random measurement error (Noise), or integrating categorical susceptibility data with MIC data (Combining MIC and categorical sensitivity). For isolates with altered MIC values, prediction error was measured as the mean Euclidean distance between predicted and observed positions of the isolates on the map (in MIC units) (Supplementary Figures S4-6). The proportion of errors within specified thresholds (<1, 1–2, and >2 MIC units) as well as the mean relative change in each isolate’s stress contribution (as a % of total stress) were also calculated. Negative values indicate a reduction in stress contribution of the isolates compared to their contribution when values were not altered.

| Test | Error (for altered isolates only) - MIC units |  |  |  |  | Mean change in stress per point (%) |
| --- | --- | --- | --- | --- | --- | --- |
|  | Mean | SD | Errors < 1 (%) | Errors 1-2 (%) | Errors >2 (%) |  |
| Missing | 0.467 | 0.625 | 86.748 | 9.760 | 3.492 | 0.023 |
| Noise | 0.895 | 0.443 | 69.861 | 27.766 | 2.372 | 0.114 |
| Combining MIC and categorical sensitivity | 0.320 | 0.681 | 86.535 | 9.435 | 4.03 | -0.03 |

The same imputation method also allowed multiple antimicrobial susceptibility assay results to be combined within a single analysis. For example, isolates characterised by categorical susceptibility only (e.g., susceptible/resistant profiles–as derived from disc-diffusion tests) could be accurately positioned on the phenotype map, enabling imputation of numeric MIC values within a single dilution (mean error = 0.320) (Supplementary Figure S5). Moreover, the framework allowed us to quantify and mitigate error (Supplementary Figure S1 and Section S1.10), delineate confidence intervals for isolate phenotypes (Supplementary Figure S3), and identify samples that have potential measurement issues based on their relative stress contribution (Supplementary Section S1.10). The method also improved interpretation of censored MIC values i.e. those beyond the dilution range of the assay, by positioning points using the information available for other drugs (Supplementary Section S1.11).

### Associating phenotypes with MLST lineages

Overlaying the 12 most common MLST types (of those with non-susceptible isolates) onto the phenotypic map revealed that lineages occupy distinct regions of phenotypic space, highlighting between-lineage variation in beta-lactam MIC (Figure 2A-B). Isolates within the same MLST mostly cluster on the map, indicating similar resistance profiles. For example, isolates of MLST 695 and 1840 showed mean pairwise phenotypic distances of 1.68 and 1.02 MIC units (Supplementary Table S11). However, some MLSTs (63, 156, and 1092) exhibited broader variation, with mean pairwise distances over 2 MIC units.

**Figure 2.**
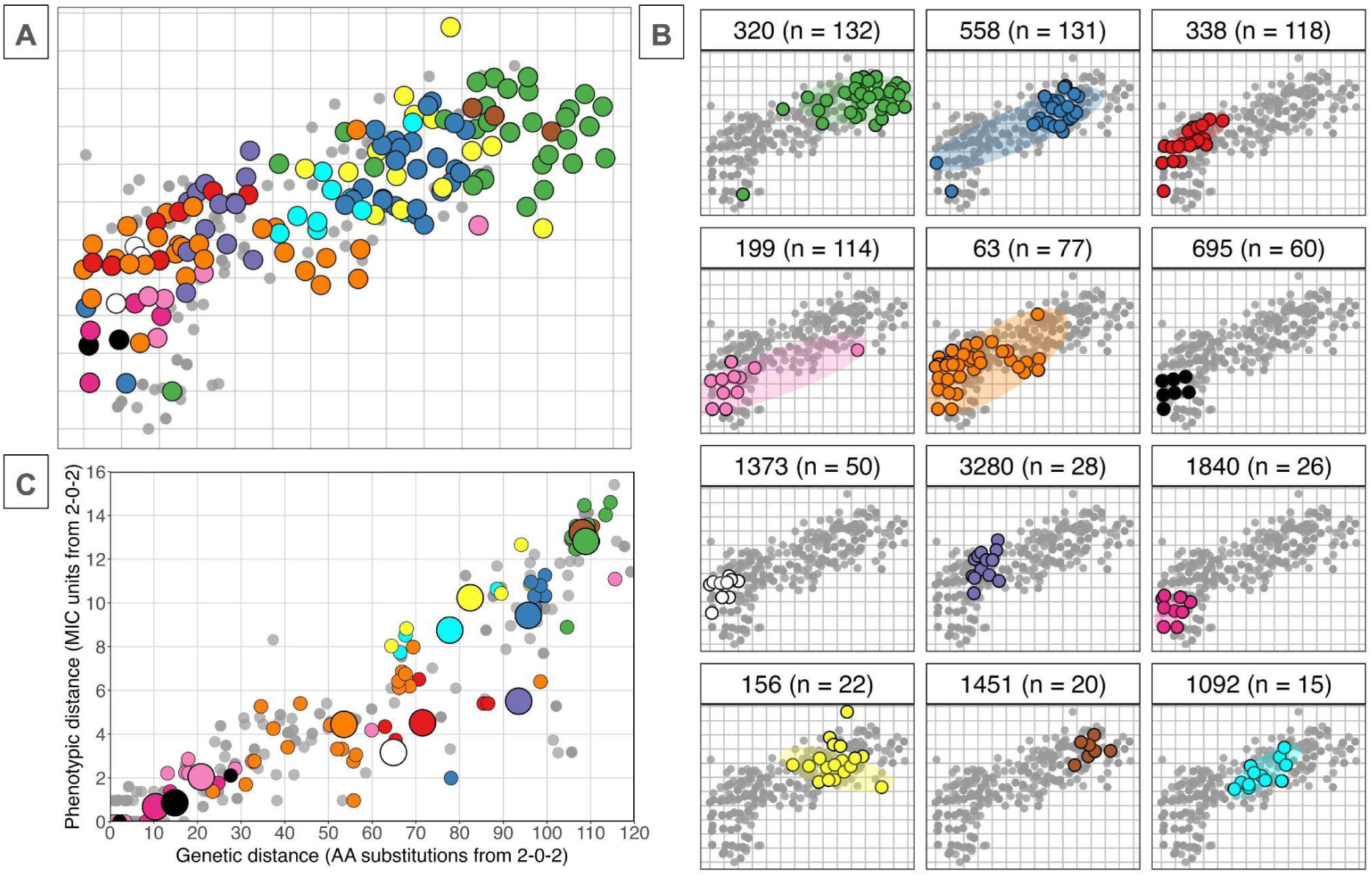
Phenotype map coloured by the 12 most common MLSTs among non-sensitive isolates. A) Coloured points highlight isolates from the 12 MLSTs; smaller grey points indicate isolates of all other MLST types. B) Isolates of each MLST type are shown separately. Shaded regions indicate the range of locations for each unique PBP sequence centroid within that MLST. Most MLSTs cluster tightly, though MLST 63, 199, and 558 show greater phenotypic variability, spanning both susceptible and intermediate MIC values. Three MLST types had extensive modification of their PBP-types but showed low-to-intermediate MIC values (338, 1373 and 3280). C) Relationship between genetic divergence in PBP transpeptidase regions and phenotypic distance on the map (MIC units), relative to the most common susceptible PBP-type (2-0-2). Small points represent the median distances for each unique PBP-type (coloured by MLST); larger points show the median distances across all PBP-types within each MLST.

Previous findings have shown PBP types containing divergent transpeptidase sequences typically display elevated MIC (8). While this was also observed here, isolates of MLSTs 338, 1373, and 3280 had extensive modification of their PBPs relative to the most common susceptible PBP-type (mean 71.53, 64.85 and 93.52 amino acid changes respectively), but exhibited only low-to-intermediate MICs (4.52, 3.16 and 5.50 MIC units difference) (Figure 2C). This highlights that while overall divergence in PBP sequence is strongly correlated with phenotypic distance, the strength and direction of this relationship varied across lineages. However, substantial shifts on the phenotype map did not occur without a corresponding change in PBP sequence, highlighting genetic variation in PBPs as a prerequisite for change in MIC. Together, these observations suggest genetic changes in the PBPs correspond strongly with differences in MICs; yet, the precise phenotypic effects of these changes depend on the specific underlying molecular variants (see below), and these can differ across evolutionary lineages.

### Identifying and visualising PBP substitutions associated with variation in beta-lactam MIC

Multivariate methods increase statistical power to identify substitutions with effects on multiple drugs, potentially revealing trade-offs between beta-lactam subclasses (30,33,34,51). We therefore used these methods to identify specific candidate PBP substitutions underlying variation in MIC. Variation in PBP transpeptidase sequences accounted for almost all variation on the phenotypic map (broad-sense heritability, *H2* = 97.6% and 95.8%, narrow-sense heritability, *h2* = 71.4% and 57.8% for axes 1 and 2 respectively), with PBP2X accounting for the majority (76% and 61.3%, respectively). To identify specific amino acid substitutions associated with changes in beta-lactam MIC, we applied four complementary approaches (Supplementary Section S2.5-9). First, we identified all pairs of isolates differing by a single amino acid substitution and quantified the corresponding shift in their positions on the phenotype map. Second, we applied clustering algorithms to define distinct groups of PBP-types and tested which substitutions segregated between regions of the phenotype map. Third, we used a multivariate linear mixed model (mvLMM) on the map axes, testing for an association between each amino acid substitution and changes in multivariate resistance phenotype. Fourth, we extended the mvLMM to test for epistatic (interaction) effects between substitutions while controlling for individual effects. Substitutions were considered to have the strongest evidence if they were supported by multiple methods.

In total, we identified 215 amino acid changes, at 179 unique PBP locations, as associated with phenotypic effects on beta-lactam MIC, representing 62.8% of all 285 varying positions within the transpeptidase regions (Table 3 and Supplementary Section S2.11). Of these, 117 changes at 108 sites were supported by two or more methods (25 in PBP1A, 42 in PBP2B, 41 in PBP2X), with PBP2X I371T identified in all four analyses (Figure 3). The effect sizes of these substitutions were within the range of 0.374 to 3.187 MIC units. Most substitutions showed evidence consistent with both additive and epistatic effects (108/117, 92.3%), with interactions both within and across proteins (Supplementary Section S2.11). Notably, several high-ranking substitutions mapped to or near conserved PBP active-site motifs (Supplementary Table S9). For example, in PBP2X, I371T, A369V, L364F, and M343T formed a cluster near the active site between the S337TMK and S395SN motifs, while V523L and S531Y lay closer to the K547SG motif (22,41). In PBP1A, T371S was within the S370TMK motif, and several further candidates lay between S370TMK and S466SN or downstream of K557TG. Moreover, PBP2B candidates with subclass-specific effects were located between S443SN and K615TG, including L609S near K615TG (22,41).

**Figure 3.**
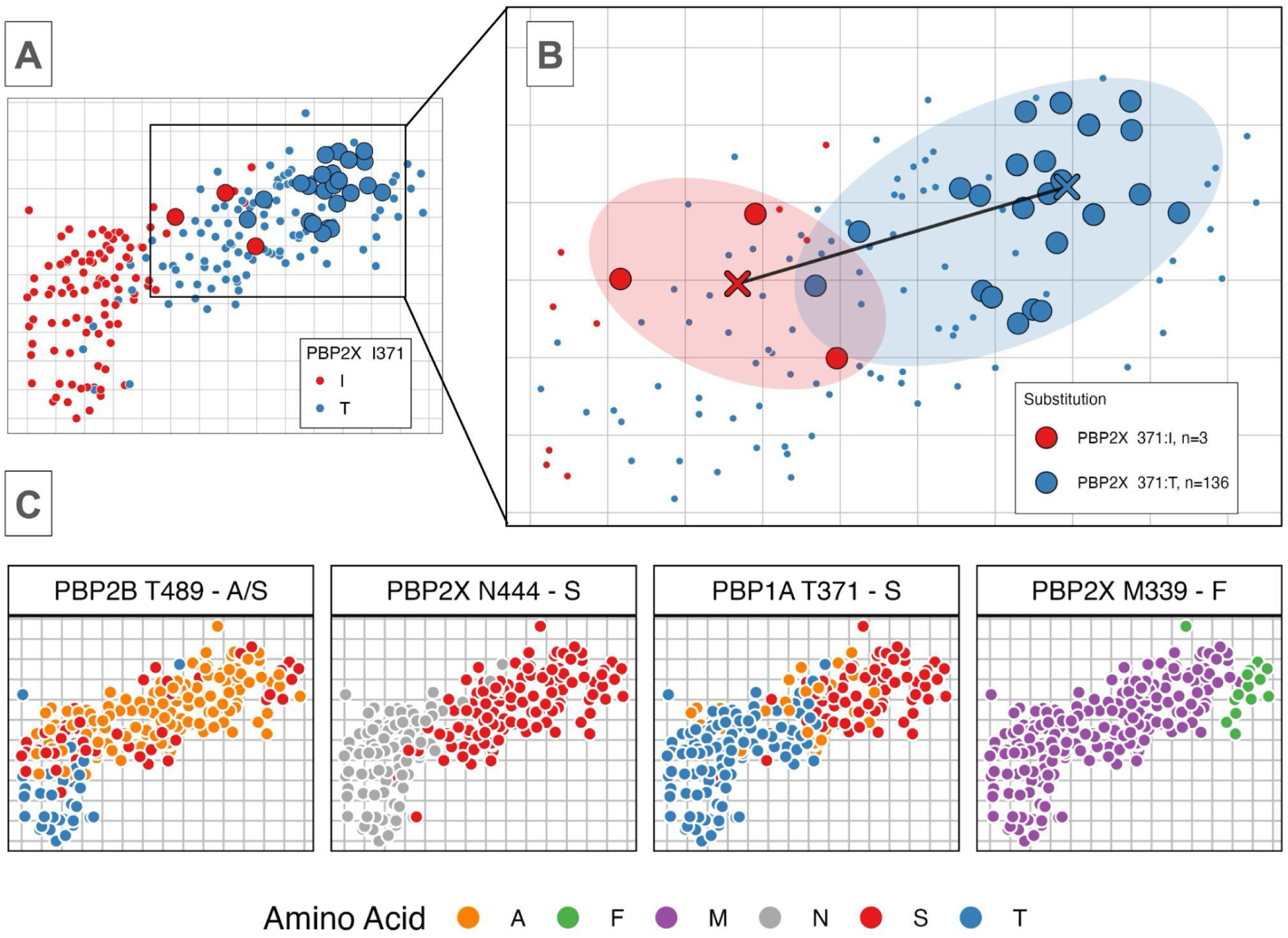
Visualising phenotypic effects of candidate PBP substitutions on beta-lactam MICs A) A single substitution comparison, where isolates on the phenotype map are coloured by the amino acid at location PBP2X-371. B) A subset of isolates (larger points) have the same PBP genetic background except for a single substitution at location PBP2X-371. The shaded areas highlight the distribution of the groups that differ in their amino acid at this location, while the coloured crosses and black line represent the median centroid positions and the phenotypic distance between them respectively. C) Amino acid changes can be visualised on the map by colouring isolates by the amino acid at a given position. The hierarchical structuring of substitutions highlights that some only appear on specific genetic backgrounds (left to right) (Supplementary Figure S11).

**Table 3.**
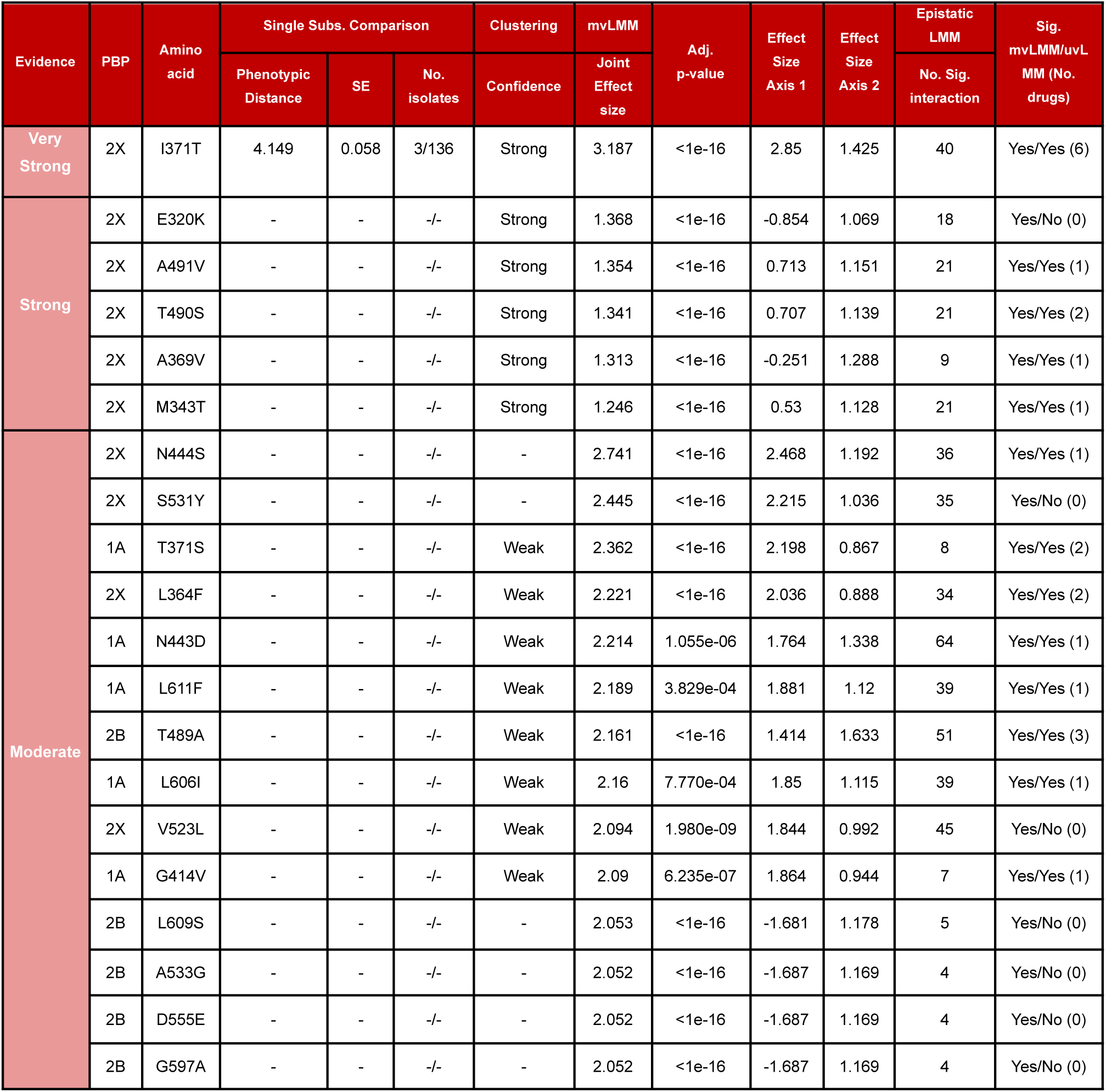
PBP substitutions associated with changes in beta-lactam MIC (only the 20 substitutions with the largest effect sizes are shown). The strength of evidence for each substitution is categorised by the number of methods which identified a phenotypic effect: 4 methods - ‘Very Strong’ evidence, 3 – ‘Strong’, 2 - ‘Moderate’, 1 – ‘Weak’, and 0 – ‘No Evidence’. ’-’ indicates an absence of evidence for a given method, such as positions without single substitution comparisons (full table in Supplementary File 1). The column ‘Sig. mvLMM/uvLMM’ compares the results of the multivariate and univariate LMMs. ‘Yes’ and ‘No’ indicates whether a substitution was significant in the mvLMM on map axes, and the uvLMM tests on individual MIC values analysis respectively. The numbers in brackets show the number of drugs which were significantly associated in the uvLMM.

Typically, substitutions were associated with specific effects on individual drug subclasses (53.85%), rather than common effects across multiple subclasses (26.49%) (Supplementary Section S3.1). However, several substitutions in PBP2X were associated with large correlated effects across multiple drugs, including I371T, N444S and S531Y. In contrast, several PBP2B variants showed evidence of contrasting phenotypic effects, increasing penicillin MIC but reducing cephalosporin and meropenem MIC, e.g. P568S, L609S and G467L/S469N/G483A/A490S. Notably, isolates from MLSTs 338 and 1373 had distinct combinations of these PBP2B mutations, even though these were uncommon in other MLST types. Correspondingly, these MLSTs showed intermediate penicillin MICs but minimal increases in cephalosporin MICs, reflecting the contrasting phenotypic effects of their PBP substitutions (Figure 2).

### Improved resolution of multivariate over univariate association methods

Using several combined multivariate methods (alongside techniques to reduce multiple testing burden) increased statistical power to detect associations between PBP substitutions and phenotypic effects. After assessing PBP positions previously identified by GWAS, random forest, and *in vitro* methods, we found that combining the cartography framework with several multivariate tests improved overlap with these prior evidence sources relative to any single method (Supplementary Figure S20). We also compared the results of the multivariate mvLMM to conventional univariate methods applied to individual MICs for each drug. Multivariate mvLMMs detected 123 candidate substitutions at 107 PBP positions which were not identified by univariate LMMs on MICs for each drug individually (Table 3 and Supplementary Section S3.3). These included several substitutions with contrasting subclass effects—such as those in PBP1A and PBP2B—11 of which were also supported by the clustering analyses.

## Discussion

In this study, we developed Antimicrobial Resistance Cartography, a multivariate framework for analysing complex, continuous resistance phenotypes and applied it to 3,628 *Streptococcus pneumoniae* isolates with MICs measured for six beta-lactam antibiotics. Our analysis revealed three main findings. First, a two-dimensional phenotype map accurately captured correlated changes in MIC values across six beta-lactam subclasses, providing simple visualisations of complex datasets with minimal information loss. Although susceptibility profiles were highly correlated across drugs, phenotypes deviated from a strictly linear path, as shown by the improved fit of a two-dimensional over a one-dimensional map. We hypothesise these divergences arise from intrinsic variation in drug-protein binding and the relative use of specific beta-lactam subclasses across populations (44,56). Second, genetic background was strongly associated with resistance phenotype, with many substitutions only appearing on the background of others, a result consistent with previous findings, but demonstrated here across multiple beta-lactam subclasses (8,38,57). While some PBP substitutions were common across MLST types, others were lineage-specific, emphasising the role of genetic context in resistance evolution. Third, multivariate methods enabled us to identify 117 candidate PBP substitutions–primarily in PBP2X and PBP2B–associated with increased beta-lactam MIC, including several with contrasting effects across subclasses, highlighting the multidimensional nature of beta-lactam resistance in this species. Together, these findings highlight the epidemiological relevance of sub-clinical MIC changes, the importance of multivariate genotype-phenotype comparisons, and the value of novel tools for phenotypic surveillance.

Beyond these core findings, the application of cartography methods addressed several technical challenges common in resistance monitoring of *S. pneumoniae*. By leveraging correlations in MIC values across antibiotics, the framework was able to impute missing or censored measurements while mitigating experimental error. We also showed it was able to integrate categorical susceptibility data with MIC data in the same analysis, enabling imputation of numeric MICs from disc-diffusion assays. These features improve the curation and quality of beta-lactam MIC data for *S. pneumoniae*, and in future, may allow better integration with genotype-phenotype prediction methods (8,58). Moreover, we also introduced several methodological refinements to genotype-phenotype association testing in bacteria, such as fine-mapping within specific genomic regions and using amino acid changes rather than nucleotides (33,51,59). Combined with the incorporation of sub-clinical variation in MIC and multivariate methods, these refinements increased statistical power (by reducing multiple-testing burden) and improved overlap with computational and *in vitro* results.

Beta-lactam resistance in *S. pneumoniae* provided an ideal case study to develop these methods, due to its complex genetic basis and phenotypic variation and covariation (6,12,13). Applying the framework to this system allowed us to examine beta-lactam resistance as a joint susceptibility phenotype rather than as six independent MIC measurements. Subsequently, these tools allowed us to pinpoint specific PBP substitutions associated with phenotypic effects and quantify how they collectively drive covariation in resistance. The resulting list of candidate PBP substitutions—including effect sizes, specific vs. common effects, and evidence strength—will aid in tracking resistance among *S. pneumoniae* populations, particularly where phenotypic data are unavailable, or novel combinations of substitutions emerge (3,8,9,48). Notably, the analysis recovered known PBP resistance regions highlighted by previous pneumococcal GWAS and PBP-signature studies, while also identifying additional substitutions associated with continuous and subclass-specific MIC variation. While *in vitro* experimental testing will be required to demonstrate the phenotypic effects of these substitutions, this list of markers may help in identifying emerging resistance during clinical interventions, such as serotype displacement after vaccination (60).

Our findings reaffirm that multiple PBP substitutions underlie increased beta-lactam MIC, each exerting small to moderate effects (0.374-3.187 MIC units). Remarkably, 179 of 285 variable positions (62.8%) within the PBP transpeptidase regions showed some evidence of phenotypic effects, many with both additive and epistatic effects. Of these, 108 positions had evidence from multiple methods, a proportion substantially higher than previous studies. For example, Chewapreecha *et al.* and Li *et al.* highlighted 31 and 27 positions as associated with high-level MIC to beta-lactams respectively (3,6,8). While this increase may be driven in part by differences across datasets or phenotypic definitions, our results highlight the greater sensitivity of multivariate approaches in detecting subtle and correlated genotype-phenotype associations (6,33,34).

Many substitutions were associated with changes in MIC below clinical breakpoints, demonstrating the value of detecting variants which generate sub-breakpoint changes in MIC, in addition to those which cause transitions between clinical thresholds. The presence of such variants among bacterial populations suggests small, incremental MIC changes are epidemiologically relevant–either by directly contributing to treatment failure rates or by facilitating stepwise progression toward clinically significant resistance (14,31,61). Even if transitory, these changes may confer competitive advantages under variable drug exposure, such as in carriage sites or biofilm formations (14,20,62). Early detection of such changes is therefore essential for informing treatment guidelines and pre-empting the emergence of high-level resistance (31,61). Further *in vitro* validation of these substitutions will be required, including in additional isolate collections, as well as exploration of other genomic regions known to affect beta-lactam MIC (e.g. *murM*) (57).

While some PBP2X substitutions (e.g. I371T, N444S and S531Y) were associated with large, correlated effects across multiple beta-lactams, most substitutions showed subclass-specific effects (53.85%). Indeed, some showed evidence of contrasting effects–for example, several changes in PBP2B increased penicillin MIC but reduced cephalosporin MIC (e.g. P568S and G467L/S469N/G483A/A490S). This pattern is consistent with subclass-specific PBP target preferences, with these substitutions potentially decreasing PBP2B binding affinity for penicillins, while simultaneously increasing it for cephalosporins (23,24,41). While *in vitro* testing will be required to validate such effects, these patterns are also highlighted by the two-dimensional phenotype map, which shows that although PBP changes can exhibit correlated effects, they often diverge, resulting in multidimensional phenotypes. This suggests that selection from one drug subclass may not necessarily confer cross-resistance to others; instead, independent changes are possible. The precise multivariate phenotypes under selection likely depend on local drug usage, fitness dynamics, linkage disequilibrium, and the target-binding specificity of drug structures (7,14,23,24,40). In this case, the strong correlations in MIC between subclasses may have likely emerged through co-occurring selective pressures, rather than direct selection for cross-resistance. This is an important distinction, as substitutions with effects on multiple drugs have a greater capacity to evade multiple antibiotics, and how this affects their relative selection pressure may determine how resistance to the entire class of drugs might evolve. In future, these insights could inform therapeutic protocols that exploit collateral sensitivity or calibrate models of co-selection in response to interventions (25,63,64).

Mapping MLSTs revealed substantial phenotypic differences between lineages, reflecting variation in selective pressures, genetic backgrounds, or recombination rate (5,7,65,66). Some MLSTs, including 320 and 156, shifted in prevalence following the introduction of pneumococcal conjugate vaccines–which coincided with increases in resistance frequencies due to serotype replacement (67,68). These MLSTs (along with 558 and 1451), exhibited uniformly high MICs, consistent with highly divergent PBPs. In contrast, other MLSTs displayed narrower MIC ranges, suggesting weaker selective pressure for increased MIC. Notably, MLSTs 338 and 1373 displayed extensive PBP divergence yet only intermediate penicillin MICs, potentially due to contrasting phenotypic effects among their PBP2B substitutions. Together, these findings suggest *S. pneumoniae* can achieve elevated MICs through multiple distinct mutational combinations, that these combinations can be specific to certain lineages, and that they are likely shaped by the extensive epistatic effects among PBPs (5,65,66). This has practical implications for clinical surveillance, as tailoring treatment to genetic backgrounds may help slow the emergence of high-level resistance (7,65). More broadly, this highlights the importance of genetic background in shaping evolutionary trajectories in this species, as it demonstrates the fitness and phenotypic outcomes of individual mutations are likely context-dependent (7,38,40,57).

More broadly, we provided a quantitative framework for analysing multivariate beta-lactam resistance in *S. pneumoniae*. By visualising several continuous phenotypic traits simultaneously, these ‘maps’ could enable further comparisons among pneumococcal populations (13,69), facilitate modelling of full susceptibility profiles (32), and help track phenotypic changes over evolutionary time (46,70,71). We hypothesise the dimensionality of the *S. pneumoniae* beta-lactam map is likely shaped by intrinsic aspects of species biology–including molecular mechanisms, fitness dynamics, and population-level antibiotic usage–and studying the map may therefore offer further insights into these factors. In future, the method could support comparative studies of *S. pneumoniae* lineages using Global Pneumococcal Sequencing data (10,35), or could be applied at smaller scales to experimental evolution studies and within-host treatment monitoring (72). Moreover, because the toolkit relies solely on phenotypic data, it could, in principle, be adapted to other species and drugs. This could also include multi-class drug resistance, as phenotypic correlations have been demonstrated across drug classes for several important bacterial pathogens, yet the causes of this are poorly understood (13). With adequate testing, the tool could be applied to MIC data from other large-scale surveillance programs, and used to help identify lineage- or species-specific genetic predispositions in acquiring resistance (73,74). AMR Cartography is therefore both a valuable clinical surveillance tool, and a framework for understanding the evolutionary dynamics which govern multivariate beta-lactam resistance in *S. pneumoniae*.

## Contributions

Conceptualisation - A.J.B., O.R. & L.A.W., Methodology - A.J.B., O.R. & L.A.W., Formal Analysis - A.J.B., Visualisations - A.J.B., Investigation - A.J.B., Writing code - A.J.B., Writing Initial Draft - A.J.B., Writing, Reviewing and Editing Subsequent Drafts - A.J.B., O.R. & L.A.W., G.G.M, S. L., Supervision - O.R. & L.A.W., Funding Acquisition - O.R. & L.A.W.. All authors provided input to the manuscript and reviewed the final version.

## Acknowledgements

We would like to thank both the CDC and ABC programme for making the data used in this manuscript publicly available. We would also like to acknowledge several current and former members of the Centre for Pathogen Evolution (Department of Zoology, University of Cambridge) for comments and feedback during development of the project. In particular, this includes David Pattinson for discussion of his PhD thesis–which inspired many of the genotype-phenotype methods used here, as well as Sam Wilks, Sarah Jones, Leah Katzelnick, Daniel Fabian, Longzhu Shen, and Derek Smith. We would also like to thank Yuan Li, Benjamin Metcalf, Lesley McGee of the CDC, the members of Stephen Bentley’s group (Wellcome Trust Sanger Institute) and Henrik Salje’s Group (Department of Genetics, University of Cambridge) for useful comments and feedback during the project. Lastly, we would like to thank Nicole Wheeler and Julian Parkhill, for examining the PhD thesis on which this work was based.

## Data availability

All MIC and PBP sequence data analysed in this study were generated by the CDC Active Bacterial Core surveillance programme and are publicly available. All analysis code and workflows, together with the source data underlying the figures and tables reported here, are available at https://github.com/AndrewBalmer/AMR-cartography. Large raw and intermediate data files are not redistributed in the repository; the scripts document how to obtain the primary data from the original sources and how to regenerate intermediate files.

## Declaration of generative AI and AI-assisted technologies in the manuscript preparation process

During the preparation of this work the author(s) used ChatGPT 5.5 in order to improve readability, grammar and syntax during writing of the manuscript, but did not use this tool for content generation. After using this tool/service, the author(s) reviewed and edited the content as needed and take(s) full responsibility for the content of the published article.

## References

1. Beall B, Chochua S, Gertz RE, Li Y, Li Z, McGee L, et al. A Population-Based Descriptive Atlas of Invasive Pneumococcal Strains Recovered Within the U.S. During 2015–2016. Front Microbiol. 2018 Nov 19;9. doi:10.3389/fmicb.2018.02670

2. McCormick AW, Whitney CG, Farley MM, Lynfield R, Harrison LH, Bennett NM, et al. Geographic diversity and temporal trends of antimicrobial resistance in Streptococcus pneumoniae in the United States. Nat Med. 2003 Apr;9(4):424–30. doi:10.1038/nm839

3. Dewé TCM, D’Aeth JC, Croucher NJ. Genomic epidemiology of penicillin-non-susceptible Streptococcus pneumoniae. Microb Genomics. 2019 Oct;5(10):e000305. doi:10.1099/mgen.0.000305 PubMed PMID: 31609685; PubMed Central PMCID: PMC6861860.

4. Schuchat A, Hilger T, Zell E, Farley MM, Reingold A, Harrison L, et al. Active bacterial core surveillance of the emerging infections program network. Emerg Infect Dis. 2001 Jan;7(1):92–9. doi:10.3201/eid0701.010114 PubMed PMID: 11266299; PubMed Central PMCID: PMC2631675.

5. Croucher NJ, Harris SR, Fraser C, Quail MA, Burton J, van der Linden M, et al. Rapid pneumococcal evolution in response to clinical interventions. Science. 2011 Jan 28;331(6016):430–4. doi:10.1126/science.1198545 PubMed PMID: 21273480; PubMed Central PMCID: PMC3648787.

6. Chewapreecha C, Marttinen P, Croucher NJ, Salter SJ, Harris SR, Mather AE, et al. Comprehensive Identification of Single Nucleotide Polymorphisms Associated with Beta-lactam Resistance within Pneumococcal Mosaic Genes. PLOS Genet. 2014 Aug 7;10(8):e1004547. doi:10.1371/journal.pgen.1004547

7. Lehtinen S, Chewapreecha C, Lees J, Hanage WP, Lipsitch M, Croucher NJ, et al. Horizontal gene transfer rate is not the primary determinant of observed antibiotic resistance frequencies in Streptococcus pneumoniae. Sci Adv. 2020 May 20;6(21):eaaz6137. doi:10.1126/sciadv.aaz6137 PubMed PMID: 32671212; PubMed Central PMCID: PMC7314567.

8. Li Y, Metcalf BJ, Chochua S, Li Z, Gertz RE, Walker H, et al. Penicillin-Binding Protein Transpeptidase Signatures for Tracking and Predicting β-Lactam Resistance Levels in Streptococcus pneumoniae. mBio. 2016 Jun 14;7(3):10.1128/mbio.00756-16. doi:10.1128/mbio.00756-16

9. D’Aeth JC, van der Linden MP, McGee L, de Lencastre H, Turner P, Song JH, et al. The role of interspecies recombination in the evolution of antibiotic-resistant pneumococci. eLife. 2021 Jul 14;10:e67113. doi:10.7554/eLife.67113 PubMed PMID: 34259624; PubMed Central PMCID: PMC8321556.

10. Gladstone RA, Lo SW, Lees JA, Croucher NJ, van Tonder AJ, Corander J, et al. International genomic definition of pneumococcal lineages, to contextualise disease, antibiotic resistance and vaccine impact. EBioMedicine. 2019 Apr 16;43:338–46. doi:10.1016/j.ebiom.2019.04.021 PubMed PMID: 31003929; PubMed Central PMCID: PMC6557916.

11. Link-Gelles R, Thomas A, Lynfield R, Petit S, Schaffner W, Harrison L, et al. Geographic and temporal trends in antimicrobial nonsusceptibility in Streptococcus pneumoniae in the post-vaccine era in the United States. J Infect Dis. 2013 Oct 15;208(8):1266–73. doi:10.1093/infdis/jit315 PubMed PMID: 23852588; PubMed Central PMCID: PMC3778966.

12. Colijn C, Cohen T, Fraser C, Hanage W, Goldstein E, Givon-Lavi N, et al. What is the mechanism for persistent coexistence of drug-susceptible and drug-resistant strains of Streptococcus pneumoniae? J R Soc Interface. 2009 Nov 25;7(47):905–19. doi:10.1098/rsif.2009.0400

13. Chang HH, Cohen T, Grad YH, Hanage WP, O’Brien TF, Lipsitch M. Origin and proliferation of multiple-drug resistance in bacterial pathogens. Microbiol Mol Biol Rev. 2015 Mar;79(1):101–16. doi:10.1128/MMBR.00039-14 PubMed PMID: 25652543; PubMed Central PMCID: PMC4402963.

14. Opatowski L, Mandel J, Varon E, Boëlle PY, Temime L, Guillemot D. Antibiotic dose impact on resistance selection in the community: a mathematical model of beta-lactams and Streptococcus pneumoniae dynamics. Antimicrob Agents Chemother. 2010 Jun;54(6):2330–7. doi:10.1128/AAC.00331-09 PubMed PMID: 20231396; PubMed Central PMCID: PMC2876390.

15. Jorgensen JH, Ferraro MJ. Antimicrobial susceptibility testing: a review of general principles and contemporary practices. Clin Infect Dis. 2009 Dec 1;49(11):1749–55. doi:10.1086/647952 PubMed PMID: 19857164.

16. Michael A, Kelman T, Pitesky M. Overview of Quantitative Methodologies to Understand Antimicrobial Resistance via Minimum Inhibitory Concentration. Anim Basel. 2020 Aug 12;10(8):1405. doi:10.3390/ani10081405 PubMed PMID: 32806615; PubMed Central PMCID: PMC7459578.

17. Negri MC, Morosini MI, Baquero MR, Campo R del, Blázquez J, Baquero F. Very Low Cefotaxime Concentrations Select for Hypermutable Streptococcus pneumoniae Populations. Antimicrob Agents Chemother. 2002 Feb;46(2):528–30. doi:10.1128/AAC.46.2.528-530.2002 PubMed PMID: 11796370; PubMed Central PMCID: PMC127056.

18. Weinstein MP, Klugman KP, Jones RN. Rationale for revised penicillin susceptibility breakpoints versus Streptococcus pneumoniae: coping with antimicrobial susceptibility in an era of resistance. Clin Infect Dis Off Publ Infect Dis Soc Am. 2009 Jun 1;48(11):1596–600. doi:10.1086/598975 PubMed PMID: 19400744.

19. Gullberg E, Cao S, Berg OG, Ilbäck C, Sandegren L, Hughes D, et al. Selection of resistant bacteria at very low antibiotic concentrations. PLoS Pathog. 2011 Jul;7(7):e1002158. doi:10.1371/journal.ppat.1002158 PubMed PMID: 21811410; PubMed Central PMCID: PMC3141051.

20. Andersson DI, Hughes D. Microbiological effects of sublethal levels of antibiotics. Nat Rev Microbiol. 2014 Jul;12(7):465–78. doi:10.1038/nrmicro3270 PubMed PMID: 24861036.

21. Baquero F, Negri MC, Morosini MI, Blázquez J. Antibiotic-selective environments. Clin Infect Dis Off Publ Infect Dis Soc Am. 1998 Aug;27 Suppl 1:S5–11. doi:10.1086/514916 PubMed PMID: 9710666.

22. Hakenbeck R, Brückner R, Denapaite D, Maurer P. Molecular mechanisms of β-lactam resistance in Streptococcus pneumoniae. Future Microbiol. 2012 Mar;7(3):395–410. doi:10.2217/fmb.12.2 PubMed PMID: 22393892.

23. Coffey TJ, Daniels M, McDougal LK, Dowson CG, Tenover FC, Spratt BG. Genetic analysis of clinical isolates of Streptococcus pneumoniae with high-level resistance to expanded-spectrum cephalosporins. Antimicrob Agents Chemother. 1995 Jun;39(6):1306–13. doi:10.1128/AAC.39.6.1306 PubMed PMID: 7574521; PubMed Central PMCID: PMC162732.

24. Kocaoglu O, Tsui HCT, Winkler ME, Carlson EE. Profiling of β-lactam selectivity for penicillin-binding proteins in Streptococcus pneumoniae D39. Antimicrob Agents Chemother. 2015;59(6):3548–55. doi:10.1128/AAC.05142-14 PubMed PMID: 25845878; PubMed Central PMCID: PMC4432181.

25. Pouwels KB, Freeman R, Muller-Pebody B, Rooney G, Henderson KL, Robotham JV, et al. Association between use of different antibiotics and trimethoprim resistance: going beyond the obvious crude association. J Antimicrob Chemother. 2018 Jun 1;73(6):1700–7. doi:10.1093/jac/dky031 PubMed PMID: 29394363.

26. Pouwels KB, Muller-Pebody B, Smieszek T, Hopkins S, Robotham JV. Selection and co-selection of antibiotic resistances among Escherichia coli by antibiotic use in primary care: An ecological analysis. PLoS One. 2019;14(6):e0218134. doi:10.1371/journal.pone.0218134 PubMed PMID: 31181106; PubMed Central PMCID: PMC6557515.

27. Seale AC, Hutchison C, Fernandes S, Stoesser N, Kelly H, Lowe B, et al. Supporting surveillance capacity for antimicrobial resistance: Laboratory capacity strengthening for drug resistant infections in low and middle income countries. Wellcome Open Res. 2017;2:91. doi:10.12688/wellcomeopenres.12523.1 PubMed PMID: 29181453; PubMed Central PMCID: PMC5686477.

28. Iskandar K, Molinier L, Hallit S, Sartelli M, Hardcastle TC, Haque M, et al. Surveillance of antimicrobial resistance in low- and middle-income countries: a scattered picture. Antimicrob Resist Infect Control. 2021 Mar 31;10(1):63. doi:10.1186/s13756-021-00931-w PubMed PMID: 33789754; PubMed Central PMCID: PMC8011122.

29. Argimón S, Masim MAL, Gayeta JM, Lagrada ML, Macaranas PKV, Cohen V, et al. Integrating whole-genome sequencing within the National Antimicrobial Resistance Surveillance Program in the Philippines. Nat Commun. 2020 Jun 1;11(1):2719. doi:10.1038/s41467-020-16322-5 PubMed PMID: 32483195; PubMed Central PMCID: PMC7264328.

30. Lees JA, Mai TT, Galardini M, Wheeler NE, Horsfield ST, Parkhill J, et al. Improved Prediction of Bacterial Genotype-Phenotype Associations Using Interpretable Pangenome-Spanning Regressions. mBio. 2020 Jul 7;11(4):e01344–20. doi:10.1128/mBio.01344-20 PubMed PMID: 32636251; PubMed Central PMCID: PMC7343994.

31. Dhand A, Sakoulas G. Reduced vancomycin susceptibility among clinical Staphylococcus aureus isolates (‘the MIC Creep’): implications for therapy. F1000 Med Rep. 2012 Feb 1;4:4. doi:10.3410/M4-4 PubMed PMID: 22312414; PubMed Central PMCID: PMC3270590.

32. Ryu S, Cowling BJ, Wu P, Olesen S, Fraser C, Sun DS, et al. Case-based surveillance of antimicrobial resistance with full susceptibility profiles. JAC Antimicrob Resist. 2019 Dec;1(3):dlz070. doi:10.1093/jacamr/dlz070 PubMed PMID: 32280945; PubMed Central PMCID: PMC7134534.

33. CRyPTIC Consortium. Quantitative measurement of antibiotic resistance in Mycobacterium tuberculosis reveals genetic determinants of resistance and susceptibility in a target gene approach. Nat Commun. 2024 Jan 12;15(1):488. doi:10.1038/s41467-023-44325-5 PubMed PMID: 38216576; PubMed Central PMCID: PMC10786857.

34. Mallawaarachchi S, Tonkin-Hill G, Croucher NJ, Turner P, Speed D, Corander J, et al. Genome-wide association, prediction and heritability in bacteria with application to Streptococcus pneumoniae. NAR Genomics Bioinforma. 2022 Mar;4(1):lqac011. doi:10.1093/nargab/lqac011 PubMed PMID: 35211669; PubMed Central PMCID: PMC8862724.

35. Gladstone RA, Lo SW, Goater R, Yeats C, Taylor B, Hadfield J, et al. Visualizing variation within global pneumococcal sequence clusters (GPSCS) and country population snapshots to contextualize pneumococcal isolates. Microb Genomics. 2020 Apr 30;6(5):1–13. doi:10.1099/MGEN.0.000357/ PubMed PMID: 32375991.

36. Hakenbeck R. Discovery of β-lactam-resistant variants in diverse pneumococcal populations. Genome Med. 2014 Sep 25;6(9):72. doi:10.1186/s13073-014-0072-8 PubMed PMID: 25473434.

37. Dowson CG, Hutchison A, Brannigan JA, George RC, Hansman D, Liñares J, et al. Horizontal transfer of penicillin-binding protein genes in penicillin-resistant clinical isolates of Streptococcus pneumoniae. Proc Natl Acad Sci U S A. 1989 Nov;86(22):8842–6. doi:10.1073/pnas.86.22.8842 PubMed PMID: 2813426; PubMed Central PMCID: PMC298386.

38. Skwark MJ, Croucher NJ, Puranen S, Chewapreecha C, Pesonen M, Xu YY, et al. Interacting networks of resistance, virulence and core machinery genes identified by genome-wide epistasis analysis. PLOS Genet. 2017 Feb 16;13(2):e1006508. doi:10.1371/journal.pgen.1006508

39. Pensar J, Puranen S, Arnold B, MacAlasdair N, Kuronen J, Tonkin-Hill G, et al. Genome-wide epistasis and co-selection study using mutual information. Nucleic Acids Res. 2019 Oct 10;47(18):e112. doi:10.1093/nar/gkz656

40. Orio AGA, Piñas GE, Cortes PR, Cian MB, Echenique J. Compensatory Evolution of pbp Mutations Restores the Fitness Cost Imposed by β-Lactam Resistance in Streptococcus pneumoniae. PLOS Pathog. 2011 Feb 17;7(2):e1002000. doi:10.1371/journal.ppat.1002000

41. Mouz N, Di Guilmi AM, Gordon E, Hakenbeck R, Dideberg O, Vernet T. Mutations in the active site of penicillin-binding protein PBP2x from Streptococcus pneumoniae. Role in the specificity for beta-lactam antibiotics. J Biol Chem. 1999 Jul 2;274(27):19175–80. doi:10.1074/jbc.274.27.19175 PubMed PMID: 10383423.

42. Zerfaß I, Hakenbeck R, Denapaite D. An Important Site in PBP2x of Penicillin-Resistant Clinical Isolates of Streptococcus pneumoniae: Mutational Analysis of Thr338. Antimicrob Agents Chemother. 2009 Mar;53(3):1107–15. doi:10.1128/AAC.01107-08 PubMed PMID: 19075056; PubMed Central PMCID: PMC2650557.

43. Enright MC, Spratt BG. A multilocus sequence typing scheme for Streptococcus pneumoniae: identification of clones associated with serious invasive disease. Microbiology. 1998 Nov;144 (Pt 11):3049–60. doi:10.1099/00221287-144-11-3049 PubMed PMID: 9846740.

44. Smith DJ, Lapedes AS, de Jong JC, Bestebroer TM, Rimmelzwaan GF, Osterhaus ADME, et al. Mapping the antigenic and genetic evolution of influenza virus. Science. 2004 Jul 16;305(5682):371–6. doi:10.1126/science.1097211 PubMed PMID: 15218094.

45. Wilks SH, Mühlemann B, Shen X, Türeli S, LeGresley EB, Netzl A, et al. Mapping SARS-CoV-2 antigenic relationships and serological responses. Science. 2023 Oct 6;382(6666):eadj0070. doi:10.1126/science.adj0070 PubMed PMID: 37797027; PubMed Central PMCID: PMC12145880.

46. Katzelnick LC, Coello Escoto A, Huang AT, Garcia-Carreras B, Chowdhury N, Maljkovic Berry I, et al. Antigenic evolution of dengue viruses over 20 years. Science. 2021 Nov 19;374(6570):999–1004. doi:10.1126/science.abk0058 PubMed PMID: 34793238; PubMed Central PMCID: PMC8693836.

47. Fonville JM, Wilks SH, James SL, Fox A, Ventresca M, Aban M, et al. Antibody landscapes after influenza virus infection or vaccination. Science. 2014 Nov 21;346(6212):996–1000. doi:10.1126/science.1256427 PubMed PMID: 25414313; PubMed Central PMCID: PMC4246172.

48. Li Y, Metcalf BJ, Chochua S, Li Z, Gertz RE, Walker H, et al. Validation of β-lactam minimum inhibitory concentration predictions for pneumococcal isolates with newly encountered penicillin binding protein (PBP) sequences. BMC Genomics. 2017 Aug 15;18(1):621. doi:10.1186/s12864-017-4017-7 PubMed PMID: 28810827.

49. Mair P, Groenen PJF, De Leeuw J. More on Multidimensional Scaling and Unfolding in R: smacof Version 2. J Stat Softw. 2022 May 13;102:1–47. doi:10.18637/jss.v102.i10

50. De Leeuw J, Mair P. Multidimensional Scaling Using Majorization: SMACOF in R. J Stat Softw. 2009 Aug 4;31:1–30. doi:10.18637/jss.v031.i03

51. Pattinson DJ. Predicting the Antigenic Evolution of Influenza Viruses with Application to Vaccination Strategy [Internet]. [Cambridge, UK]: University of Cambridge; 2020 [cited 2025 Aug 15]. Available from: https://www.repository.cam.ac.uk/items/0fd9bca6-c64b-47cd-92b4-34302f5e6bd6

52. Lippert C, Listgarten J, Liu Y, Kadie CM, Davidson RI, Heckerman D. FaST linear mixed models for genome-wide association studies. Nat Methods. 2011 Oct;8(10):833–5. doi:10.1038/nmeth.1681

53. Lippert C, Casale FP, Rakitsch B, Stegle O. LIMIX: genetic analysis of multiple traits [Internet]. bioRxiv; 2014 [cited 2025 Aug 15]. p. 003905. Available from: https://www.biorxiv.org/content/10.1101/003905v1doi:10.1101/003905

54. Brennan-Krohn T, Smith KP, Kirby JE. The Poisoned Well: Enhancing the Predictive Value of Antimicrobial Susceptibility Testing in the Era of Multidrug Resistance. J Clin Microbiol. 2017 Aug;55(8):2304–8. doi:10.1128/JCM.00511-17 PubMed PMID: 28468856; PubMed Central PMCID: PMC5527407.

55. Andrews JM. Determination of minimum inhibitory concentrations. J Antimicrob Chemother. 2001 Jul 1;48(Suppl 1):5–16. doi:10.1093/jac/48.suppl_1.5

56. Bedford T, Rambaut A, Pascual M. Canalization of the evolutionary trajectory of the human influenza virus. BMC Biol. 2012 Apr 30;10(1):38. doi:10.1186/1741-7007-10-38

57. Gibson PS, Bexkens E, Zuber S, Cowley LA, Veening JW. The acquisition of clinically relevant amoxicillin resistance in Streptococcus pneumoniae requires ordered horizontal gene transfer of four loci. PLOS Pathog. 2022 Jul 25;18(7):e1010727. doi:10.1371/journal.ppat.1010727

58. Batisti Biffignandi G, Chindelevitch L, Corbella M, Feil EJ, Sassera D, Lees JA. Optimising machine learning prediction of minimum inhibitory concentrations in Klebsiella pneumoniae. Microb Genomics. 2024 Mar;10(3):001222. doi:10.1099/mgen.0.001222 PubMed PMID: 38529944; PubMed Central PMCID: PMC10995625.

59. Schaid DJ, Chen W, Larson NB. From genome-wide associations to candidate causal variants by statistical fine-mapping. Nat Rev Genet. 2018 Aug;19(8):491–504. doi:10.1038/s41576-018-0016-z PubMed PMID: 29844615; PubMed Central PMCID: PMC6050137.

60. Lekhuleni C, Ndlangisa K, Gladstone RA, Chochua S, Metcalf BJ, Li Y, et al. Impact of pneumococcal conjugate vaccines on invasive pneumococcal disease-causing lineages among South African children. Nat Commun. 2024 Sep 27;15(1):8401. doi:10.1038/s41467-024-52459-3

61. Colangeli R, Jedrey H, Kim S, Connell R, Ma S, Chippada Venkata UD, et al. Bacterial Factors That Predict Relapse after Tuberculosis Therapy. N Engl J Med. 2018 Aug 30;379(9):823–33. doi:10.1056/NEJMoa1715849 PubMed PMID: 30157391; PubMed Central PMCID: PMC6317071.

62. Negri MC, Lipsitch M, Blázquez J, Levin BR, Baquero F. Concentration-dependent selection of small phenotypic differences in TEM beta-lactamase-mediated antibiotic resistance. Antimicrob Agents Chemother. 2000 Sep;44(9):2485–91. doi:10.1128/AAC.44.9.2485-2491.2000 PubMed PMID: 10952599; PubMed Central PMCID: PMC90089.

63. Liakopoulos A, Aulin LBS, Buffoni M, Fragkiskou E, van Hasselt JGC, Rozen DE. Allele-specific collateral and fitness effects determine the dynamics of fluoroquinolone resistance evolution. Proc Natl Acad Sci U S A. 2022 May 3;119(18):e2121768119. doi:10.1073/pnas.2121768119 PubMed PMID: 35476512; PubMed Central PMCID: PMC9170170.

64. Hernando-Amado S, Laborda P, Martínez JL. Tackling antibiotic resistance by inducing transient and robust collateral sensitivity. Nat Commun. 2023 Mar 30;14(1):1723. doi:10.1038/s41467-023-37357-4

65. Lehtinen S, Blanquart F, Croucher NJ, Turner P, Lipsitch M, Fraser C. Evolution of antibiotic resistance is linked to any genetic mechanism affecting bacterial duration of carriage. Proc Natl Acad Sci U S A. 2017 Jan 31;114(5):1075–80. doi:10.1073/pnas.1617849114 PubMed PMID: 28096340; PubMed Central PMCID: PMC5293062.

66. Hanage WP, Fraser C, Tang J, Connor TR, Corander J. Hyper-recombination, diversity, and antibiotic resistance in pneumococcus. Science. 2009 Jun 12;324(5933):1454–7. doi:10.1126/science.1171908 PubMed PMID: 19520963.

67. Dagan R, Klugman KP. Impact of conjugate pneumococcal vaccines on antibiotic resistance. Lancet Infect Dis. 2008 Dec;8(12):785–95. doi:10.1016/S1473-3099(08)70281-0 PubMed PMID: 19022193.

68. Kyaw MH, Lynfield R, Schaffner W, Craig AS, Hadler J, Reingold A, et al. Effect of introduction of the pneumococcal conjugate vaccine on drug-resistant Streptococcus pneumoniae. N Engl J Med. 2006 Apr 6;354(14):1455–63. doi:10.1056/NEJMoa051642 PubMed PMID: 16598044.

69. Wildfire J, Waterlow NR, Clements A, Fuller NM, Knight GM. MIC distribution analysis identifies differences in AMR between population sub-groups. Wellcome Open Res. 2024;9:244. doi:10.12688/wellcomeopenres.21269.1 PubMed PMID: 39119595; PubMed Central PMCID: PMC11306957.

70. Kenyon C, Laumen J, Van Den Bossche D, Van Dijck C. Where have all the susceptible gonococci gone? A historical review of changes in MIC distribution over the past 75 years. BMC Infect Dis. 2019 Dec 27;19(1):1085. doi:10.1186/s12879-019-4712-x

71. Bedford T, Suchard MA, Lemey P, Dudas G, Gregory V, Hay AJ, et al. Integrating influenza antigenic dynamics with molecular evolution. eLife. 2014 Feb 4;3:e01914. doi:10.7554/eLife.01914

72. Balmer A. Multivariate Methods for the Study of Beta-lactam Resistance in Streptococci [PhD Thesis] [Internet]. [Cambridge, UK]: University of Cambridge; 2025 [cited 2025 Aug 15]. Available from: https://www.repository.cam.ac.uk/items/a20b3ee7-5ae5-417f-a95f-15299a52caff

73. Global antimicrobial resistance and use surveillance system (GLASS) report: 2022 [Internet]. [cited 2024 Jun 22]. Available from: https://www.who.int/publications/i/item/9789240062702

74. Karp BE, Tate H, Plumblee JR, Dessai U, Whichard JM, Thacker EL, et al. National Antimicrobial Resistance Monitoring System: Two Decades of Advancing Public Health Through Integrated Surveillance of Antimicrobial Resistance. Foodborne Pathog Dis. 2017 Oct 1;14(10):545–57. doi:10.1089/fpd.2017.2283 PubMed PMID: 28792800; PubMed Central PMCID: PMC5650714.

