## Supplementary Information for "Mapping the Phenotypic Landscape of Beta-lactam Resistance in Streptococcus pneumoniae"

**Supplementary Section 1 - Isolate collection, map generation and testing** **3**
1. Isolate collection 3
2. Pairwise relationships between drugs 4
3. Phenotype (and genotype) map construction and testing 4
4. Error principle of MDS 4
5. Finding an optimal dimensionality 5
6. Metric vs non-metric MDS 6
7. Rotation, dilation, density distributions and calibrated biplots 7
8. Assessing map goodness-of-fit 9
9. Estimating coordination confidence 12
10. Cross-validation analyses 14
11. Inclusion of censored values on phenotype maps 21
12. Dimensionality tests 23

**Supplementary Section 2 - Genetic analyses in detail** **26**
1. PBP-types in S. pneumoniae 26
2. Genetic map and clustering 27
3. Heritability and per-PBP variance decomposition 30
    3.1. 'Broad-sense' heritability (H2) 30
    3.2. 'Narrow-sense' heritability (h2) 31
    3.3. Per-PBP variance decomposition 32
    3.4. Heritability results 34
4. Strong relationship between number of amino acid changes from susceptible PBP-type and beta-lactam MIC 36
    4.2 Phenotypic and genetic diversity within MLSTs 38
5. Identification of causal amino acid substitutions 41
6. Identifying pairs of isolates that differ by a single PBP substitution 41
7. Identifying 'cluster-difference' substitutions 45
8. Multivariate linear mixed models (LMMs) 48
9. Epistatic interactions between PBP loci 50
10. Increasing the statistical power of mvLMMs 51
    10.1. Focus on the transpeptidase regions of the PBP proteins only 51
    10.2. Applying mvLMMs to phenotype map axes 51
    10.3. Amino acid sequences rather than genetic sequences 51
    10.4. Dummy variables 52
    10.5. Per-test variance decomposition 52
    10.6. Correction of p-values for multiple comparisons 52

10.7. Setting conservative p-value thresholds 53
11. mvLMMs identify many PBP substitutions with additive and epistatic effects 55
12. Ranking PBP substitutions based on strength of evidence for phenotypic effect 57

**Supplementary Section 3 - Additional Analyses 59**
1. Many identified substitutions have contrasting effects on different subclasses of beta-lactams 59
2. PBP substitutions identified using multivariate methods show strong overlap with previously published work 61
3. Multivariate methods identify additional associations compared to univariate counterparts 62

### *Supplementary Section 1 - Isolate collection, map generation and testing*

1. *Isolate collection*

We used a dataset of 4,309 invasive pneumococcal isolates collected in the United States through the Active Bacterial Core surveillance (ABC) from 1998 to 2015 (1–3). The isolates were obtained from sterile infection sites from patients across 10 U.S. states. MIC values for six beta-lactam antibiotics were measured using 2-fold broth microdilution (Supplementary Table S1). Of the 4,309 isolates, 3,628 had MIC data for all six drugs, and only these were included in subsequent analyses. Genomic data were collected via Illumina HiSeq or MiSeq sequencing, determining capsular serotype, MLST, pilus type, and PBP amino acid sequences. Relevant metadata, including date, geographical location, and clinical symptoms, are available through the ABC programme. These datasets were originally published to develop machine learning models which can predict beta-lactam MIC values from PBP-type (2,3). Since the development of these models, surveillance of beta-lactam resistance has continued, although *in vitro* MIC testing is only conducted on a subset of new strains identified through the ABC programme.

**Supplementary Table S1.** Descriptions of drugs measured, dilution ranges and number of isolates measured for each. Of the 4309 isolates in the collection, 3628 had MIC available for all 6 drugs.

| Sub-class | Antibiotic | No isolates measured | Observed MIC range (µg/mL) | No. dilution steps performed |
|---|---|---|---|---|
| Penicillin | Penicillin | 4307 | 0.03-16 | 10 |
| | Amoxicillin | 4293 | 0.03-16 | 10 |
| Carbapenem | Meropenem | 4224 | 0.06-2 | 6 |
| Cephalosporin | Cefotaxime | 4244 | 0.06-16 | 9 |
| | Ceftriaxone | 3648 | 0.5-8 | 5 |
| | Cefuroxime | 4301 | 0.5-4 | 4 |

*2. Pairwise relationships between drugs*

Isolates showed extensive phenotypic variation and strong pairwise covariation across the six antibiotics (Supplementary Table S1 and Figure 1 - Main Text). One exception was cefuroxime, where most isolates had one of two censored MIC values (≤ 0.5 or ≥ 4μg/mL). MIC values for penicillin and amoxicillin displayed the widest range, varying by up to nine and ten log2 dilutions respectively (0.03-16μg/mL). Isolates also showed extensive variation in cefotaxime MIC (0.06-16μg/mL). Pairwise correlations between drugs were high (correlation coefficients = 0.77-0.96). For all antibiotics, most of the isolates had an MIC equal to (or lower than) the lowest concentration tested. These censored values may not accurately represent their true MIC value, as the true values could be lower than this (see Supplementary Section S1.11), and results in lower correlation coefficients between antibiotics for some drugs e.g. penicillin and ceftriaxone (correlation coefficient = 0.77).

*3. Phenotype (and genotype) map construction and testing*

Phenotype maps visualise a Euclidean distance matrix derived from MIC data, where each point represents an isolate and distances between them reflect differences in MIC values across all drugs. The axes serve to help visualise the distances between isolates, with grid squares representing $\log_2$ dilution steps in MIC values (Figure 1 - Main text). MIC values were $\log_2$-transformed to approximate a normal distribution. Pairwise Euclidean distances were then calculated between isolates to form an initial matrix (table distance - $d_{i,j}$). In generating this matrix, lower censored values (e. g. “<= X”) were approximated as X; and a value of “>X” as 2X, as described in: (2,3). As MDS can work using metric or rank data, this approximation did not significantly affect the final representation (discussed in detail in Supplementary Sections 1.3 and 1.11).

Genetic maps use a Hamming distance matrix based on an amino acid sequence alignment (Supplementary Figure S8) (4,5). Genetic distances are calculated using amino acid substitutions across the concatenated transpeptidase regions of the three PBP proteins (914 positions-see Supplementary Section 2.1). These maps represent genetic variation in a similar way to how phenotype maps represent MIC data.

*4. Error principle of MDS*

MDS aims to find a low-dimensional representation of target distances with minimal error (6,7). Distances in the representation are found by iteratively minimising the differences between map distances ($d_{i,j}$) and table distances ($D_{i,j}$) for every pair of isolates i and j, through an error function called stress (S):

$$S = \sum_{i,j}(D_{i,j} - d_{i,j})^2 \tag{2.1}$$

‘Stress’ refers to the sum of the squared residuals between the distances on the map and the original measured distances for each isolate pair (8). The lower the stress, the better the map fits the data, and can be used to compare solutions for the same dataset under different starting conditions and targets (8).

The MDS algorithm searches for a local minimum of the stress value by iteratively moving isolates on the map. As different starting positions can affect the final positions of the points, two strategies were used to search for a global minimum. One uses the output of a classical MDS (i. e. PCA) as the starting positions, while the other repeats the search multiple times using random start positions. As both methods were comparable, and the PCA method is easier to run and cross-validate, all maps shown in this manuscript were found using PCA as starting positions.

*5. Finding an optimal dimensionality*

To determine the optimal number of dimensions to capture the variation in the data, we generated maps in several different dimensionalities and compared their relative stress (Supplementary Table S2). For both phenotype and genotype maps, there was a large decrease in stress moving from 1D to 2D and from 2D to 3D, but little decrease when adding additional dimensions. The percentage of pairwise errors above 1 MIC unit dropped significantly when moving from 1D to 2D, from 1.509% to 0.219% (Table 1 - Main Text). Overall stress was low for the 2D maps, indicating a good fit to the data. Notably, the 3D phenotype map had lower total stress and fewer pairwise errors than the 2D map. However, isolate distributions on the 2D and 3D maps remained directly comparable and the 2D maps performed better in cross-validation analyses (see Supplementary Section 1.10). As a 2D map is easier to work with and visualise, the 2D maps are presented throughout this work. Importantly, results of genotype-phenotype comparisons were consistent regardless of whether 2D or 3D maps were used for phenotype or genetic data (Supplementary Section 2).

*6. Metric vs non-metric MDS*

The SMACOF algorithm was chosen due to its flexibility in distance transformations. Metric MDS directly preserves the measured distances in the table, while ordinal MDS preserves only rank values, offering more flexibility where there are many censored values. A third option is an interval transformation, which preserves the relative distance between points on each scale but can have separate transformations for each drug or subclass of drugs individually. All transformations yielded very similar map representations. Overall, ordinal MDS produced the lowest stress maps, which was expected given it is the least strict assumption (Supplementary Table S2). However, metric maps also fit the data very well, and as this transformation is more directly interpretable, metric MDS was used for all maps presented here.

**Supplementary Table S2.** Percentage decreases in stress relative to the stress for a one-dimensional map. Note bold and underlined indicates a degenerate solution for the one-dimensional genotype map.

| | **Percentage decrease in stress (%)** | | | | |
|---|---|---|---|---|---|
| **Map** | **Transformation** | **1D to 2D** | **2D to 3D** | **3D to 4D** | **4D to 5D** |
| **Phenotype** | Metric | 55.534 | 25.428 | 7.064 | 5.509 |
| | Ordinal | 44.306 | 31.221 | 8.039 | 11.285 |
| | Interval | 54.238 | 26.004 | 7.293 | 5.768 |
| **Genotype** | Metric | 62.547 | 17.356 | 6.25 | 2.741 |
| | **<u>Ordinal</u>** | **<u>NA</u>** | **<u>NA</u>** | **<u>NA</u>** | **<u>NA</u>** |
| | Interval | 63.288 | 16.872 | 5.819 | 2.674 |

*7. Rotation, dilation, density distributions and calibrated biplots*

As MDS is only concerned with finding the optimal pairwise distances between points, their rotation and dilation can be adjusted as needed. To dilate the maps to an interpretable scale, we used the slope of the relationship between the pairwise table distances and the pairwise distances on the map (Supplementary Table S3). The borders of the plots were chosen to correspond to the range of dilutions tested for the different drugs, so for any area of the map, isolates could have potentially fallen within that range of combinations. Density distributions and marginal histograms were included to aid visualisation in cases where isolates lie atop one another (Figure 1B - Main Text).

External variables can be mapped onto the plot using colour (Supplementary Figure S2) and biplot vectors (Figure 1C - Main Text), and calibrated by drawing tick marks which correspond to the isolate values on the plot ('Calibrate' package (9)). The distribution of MIC values were well fit across all drugs, although in general those with fewer numerical titres were less well fit (Supplementary Figure S1).

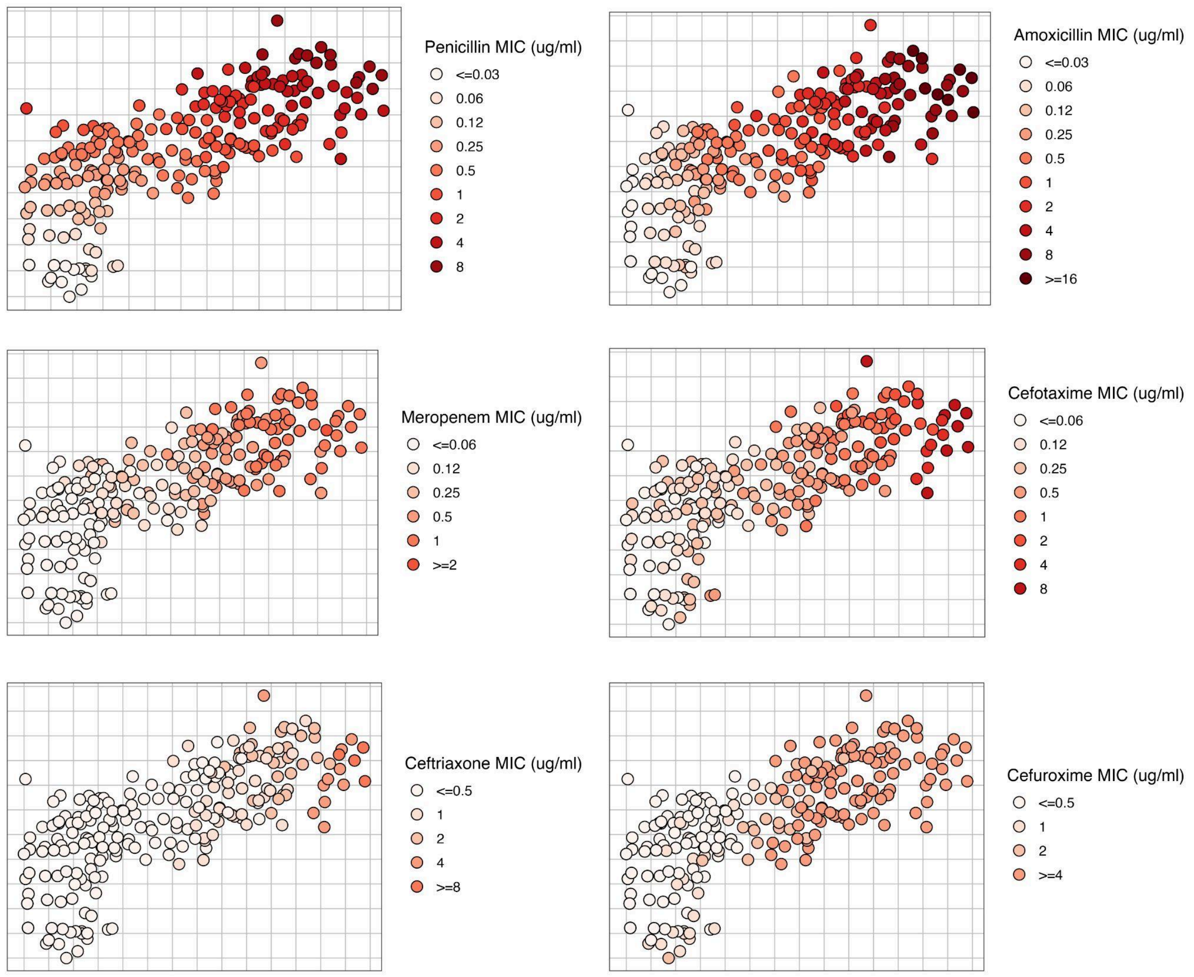


**Supplementary Figure S1.** Distribution of MIC values for each drug mapped using colour, where increasing MIC is shown in darker shades of red. Notably, all drugs were well fit across the map, though the fit of individual drugs varied depending on the proportion of censored MIC values present. For example, most isolates had upper or lower bound censored values for both ceftriaxone and cefuroxime. In cases where an isolate has censored values to one or more drugs, MDS uses the values across other drugs to position isolates on the map. This feature of the method allows interpretation of phenotypes beyond the measured dilution range for any individual drug.

*8. Assessing map goodness-of-fit*

The goodness-of-fit of the MDS plots were investigated by calculating the proportion of errors above certain thresholds and investigating the distribution of stress-per-point (as a % of total stress) (Supplementary Figure S2 and Table 1 - Main Text). A Shepard plot can be used to show the relationship between measured pairwise table distances against the pairwise map distances (6,8). This relationship can be assessed quantitatively using a linear model, where a map which perfectly fits the data would have an R-squared of 1 and an intercept of 0 (4,5). Overall, both phenotype and genotype maps fit the data well, with the Shepard plot regression slopes and $R^2$ values close to 1, and intercepts close to 0. For both maps, there were few examples of error over a single $\log_2$ dilution or 10 amino acids. Stress-per-point (%) was also low, especially among the most susceptible isolates, i.e. those with low or censored values to all drugs tested.

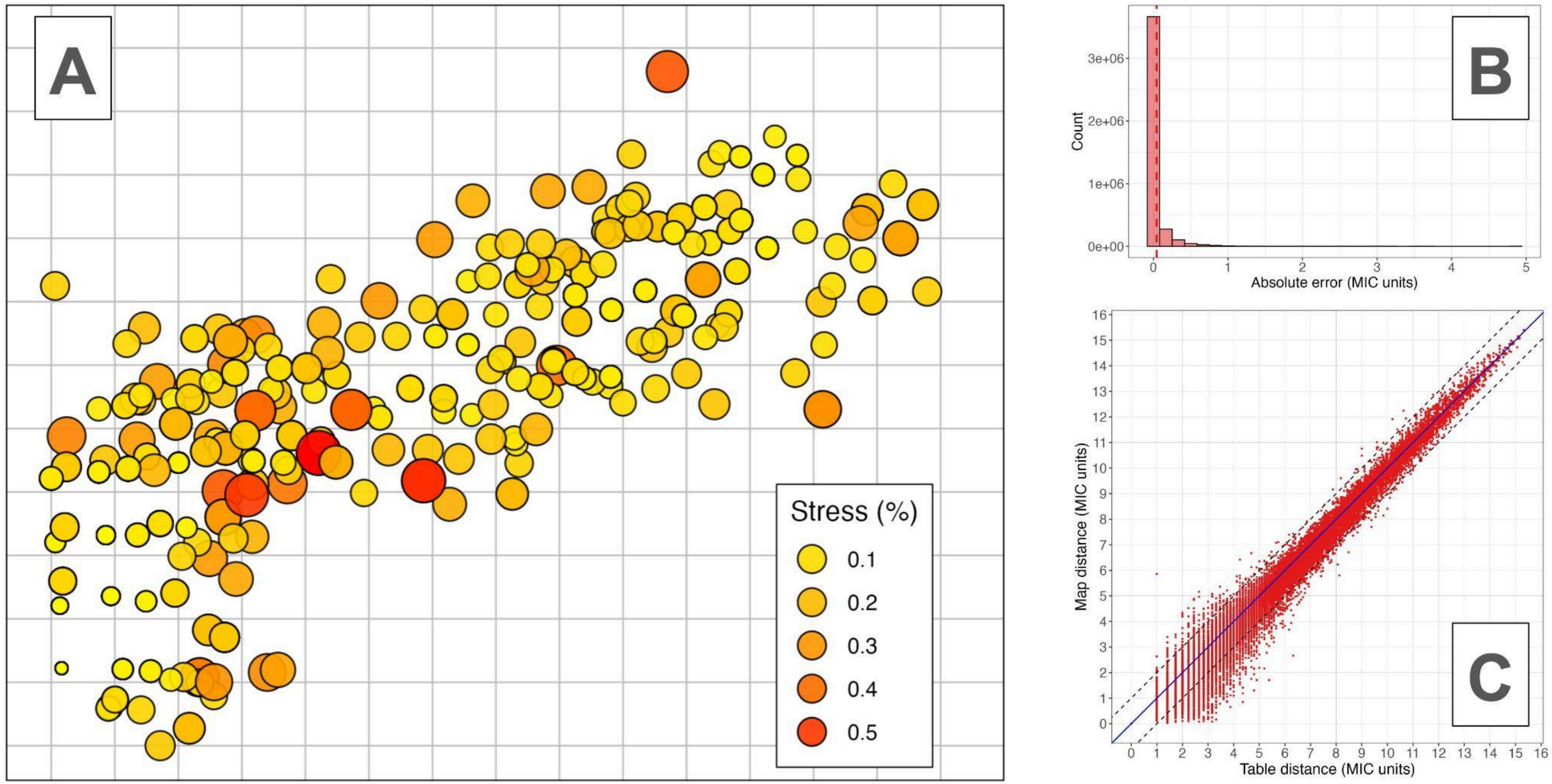


**Supplementary Figure S2.** Goodness-of-fit plots for *S. pneumoniae* map. Panel A shows the phenotype map, where the size and colour of the points represents their contribution to total stress on the map. Larger, red points indicate isolates with greater relative stress contributions. Panel B shows the pairwise residuals of all isolates as a histogram, where the dashed line indicates the mean of the distribution. Panel C shows a Shepard plot of the pairwise distances between points as measured in the assay and their pairwise distances on the map (after dilation). The solid black line shows a linear model of the relationship between the table and map distances. The dashed lines indicate one MIC unit deviation from a perfect fit, meaning points outside this line have an error above one MIC unit.

**Supplementary Table S3.** Goodness-of-fit statistics for phenotype and genotype maps.

| | Shepard plot | | | | | Pairwise errors (MIC units) | | | | Stress-per-point (%) | |
|---|---|---|---|---|---|---|---|---|---|---|---|
| **Map** | **Slope** | **Intercept** | **R.S.E.** | **Adj. $R^2$** | **Corr. Coef** | **>1-unit error (%)** | **>2-unit error (%)** | **Mean error** | **SD error** | **Mean stress per-point** | **SD stress per point** |
| **Phenotype** | 0.184 | -0.004 | 0.021 | 0.999 | 0.999 | 0.219 | 0.010 | 0.035 | 0.108 | 0.028 | 0.057 |

| | Shepard plot | | | | | Pairwise error (AA distance) | | | | Stress-per-point (%) | |
|---|---|---|---|---|---|---|---|---|---|---|---|
| **Map** | **Slope** | **Intercept** | **R.S.E.** | **Adj. $R^2$** | **Corr. Coef** | **>10 AA error (%)** | **>20 AA error (%)** | **Mean error** | **SD error** | **Mean stress per-point** | **SD stress per point** |
| **Genotype** | 0.018 | -0.019 | 0.082 | 0.988 | 0.994 | 3.648 | 1.189 | 1.973 | 4.182 | 0.028 | 0.054 |

*9. Estimating coordination confidence*

Depending on the dataset and method used, some MDS solutions can be ‘unstable’, i. e. the patterns observed on the map are substantively different when a subsample of the full dataset is used. We used three methods to ensure the stability of the MDS representations; jack-knife analysis, bootstrapping and pseudo-confidence intervals (6,8). Firstly, jack-knifing involves removing each isolate in turn from the dataset, while keeping all the others, remaking the map, and testing how similar the maps are. The map was highly stable to jack-knife resampling, with the process essentially having no effect on isolate map positions (Supplementary Table S4). Secondly, we bootstrap resampled the isolates of each dataset 100 times with replacement, generated a map for each and compared the solutions to the map made with the observed dataset. All bootstrapped solutions were close to a perfect fit, showing the map is robust to bootstrap resampling (Supplementary Table S4). Thirdly, we generated ‘pseudo-confidence’ intervals using the SMACOF algorithm. This is done by shifting the position of each isolate on the map while keeping the others stationary, until stress increases by a set amount (a proportional increase of 1/number of isolates). The confidence intervals produced using this method were essentially the same as the bootstrapping method (Supplementary Figure S3). In conclusion, the map was highly stable to several resampling methods, indicating high coordination confidence.

**Supplementary Table S4.** Coordination confidence analyses for *S. pneumoniae* phenotype maps. Stability coefficient represents the between/total variance ratio across the resampled maps.

| | **Jack-knife** | | | **Bootstrapping** | |
|---|---|---|---|---|---|
| **Map** | **Stability coefficient** | **Cross validity** | **Dispersion** | **Mean bootstrap stress** | **Stability coefficient** |
| **Phenotype** | 1 | 1 | 0 | 0.013 | 0.980 |

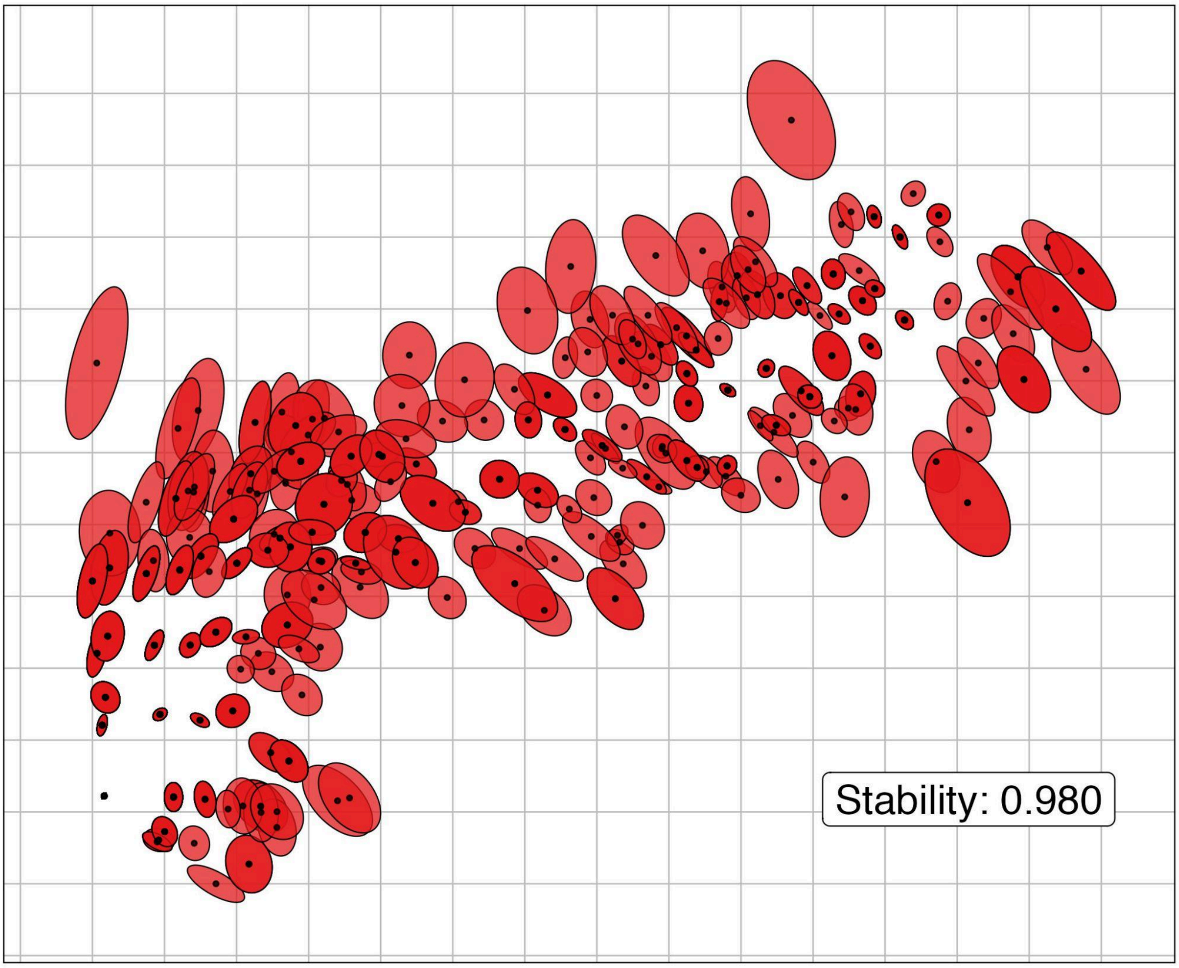


**Supplementary Figure S3.** Defining confidence intervals for phenotype maps based on bootstrap resampling with replacement for *S. pneumoniae*. Isolate mean centroid positions are marked in black after 100 bootstrap samples of the isolates with replacement. Margins indicate 95% confidence intervals of the mean centroid positions across bootstrap resamples, which were typically within one MIC unit in any direction.

*10. Cross-validation analyses*

In theory, because MDS maps position the isolates using several correlated variables, measurement error is averaged out, meaning the maps should be more accurate than any single MIC value taken individually (6,8). This was shown to be the case for haemagglutination titres in the original H3N2 influenza maps (4). This property of MDS is important, as when the MIC assay is conducted for many isolates, it is common for individual MIC values to be missing, or to have experimental errors in their measurement. In other cases, different assay methods have been used to measure phenotypes within a collection. These issues can occur for several reasons, such as limitations in time or resources, an inability to grow an isolate from culture on a given day, or where collaborators have contributed data using a different assay method. Experimental errors are also common, for example, as the MIC assay works on discrete 2-fold dilutions, a value of 4μg/mL can in fact be any value between 2μg/mL and 4μg/mL, meaning a small error in dilution can result in an MIC either side of its true value (10,11). Taken together, these issues pose a challenge for analysis, particularly when isolates have errors for several drugs. In some cases, separate statistical analyses need to be conducted, complicating interpretation, and resulting in a loss of statistical power in downstream analysis.

In contrast to univariate analysis, where these isolates are often simply excluded, multivariate methods can use the information present for other drugs to impute values. We tested whether this meant MDS could mitigate common issues such as measurement error, missing values, and combined AST methods. To do this, we conducted three separate analyses, each designed to test a different type of error. For each test, we generated 100 duplicate datasets, and added error to these datasets under one of three error schemes:

- *Missing values* – we randomly removed 10% of the MIC values from the dataset. To calculate a target distance matrix, for the isolates with missing values, pairwise distances were estimated by excluding the column with the missing value and using only the values present for each comparison.
- *Experimental noise* - we randomly selected 10% of the MIC values from the dataset. If the value was a numeric titre, +/- one $\log_2$ MIC dilution was added to this value, but if the titre was a censored value, then no error was added. Censored values were excluded because simply adding +1/-1 $\log_2$ dilution to a censored value could add a relatively large error if an isolates true value is in fact far below the dilution range, meaning the test would be invalid.

- *Combining AST methods* - we randomly selected 10% of isolates from the dataset and replaced their numeric MIC values with categorical susceptibility profiles (susceptible/resistant) based on clinically defined MIC breakpoints for each drug. For example, if a resistant breakpoint for a given drug was => 2μg/mL, an isolate which had an MIC value equal or higher than 2μg/mL for that drug was marked as 'R' – for resistant. We then calculated the most common MIC values for each combination of susceptible/resistant cut-offs among the remaining isolates with MIC, as each isolate has a combination of cut-off values for the 6 drugs:

  e. g. 'S S R S R S' - where S = susceptible and R = resistant.

  Since this is based on MIC values (e. g. 'R' for a given drug => 2μg/mL), it is possible to calculate the cut-off combination for isolates which did have their MIC values measured. For each combination of cut-offs, we calculated the most common MIC values for the six drugs among the remaining 90% of isolates,

  e. g. 'S S R S R S' = 1 - 2 - 8 - 1 - 16 - 1 (where digits refer to μg/mL)

  Then, for the isolates with cut-off data only, we imputed these MIC values, as they were the most common values among isolates with the same cut-off combination. A pairwise distance matrix for each dataset was then generated using these values and used as input for the MDS algorithm.

We generated maps for each of the 100 samples, for each error scheme, and aligned them with the original map using Procrustes (Supplementary Figure S4). Prediction error for MIC values was estimated using the Euclidean distance between the isolate position on the full map and the isolate position on the map made with altered values (12). The overall stability of maps with altered values was also assessed using the congruence coefficient, defined as the correlation of isolate positions on each axis around the origin (6). Generally in MDS, a coefficient above 0.95 is considered satisfactory. As an additional cross-validation, maps were made in different dimensionalities (1, 2, 3, 4, and 5), using the first 25 of the modified datasets and their respective error scores compared.

*Weighted multidimensional scaling*

For the missing values and combined susceptibility category analyses, we incorporated weighting structures to prevent isolates with missing or categorical susceptibility data from disproportionately influencing positions of isolates which had more information on where to position them. This was achieved by giving them less weight when generating the map. To do this, for each analysis, a weight matrix was constructed for each of the 100 samples. Here, the weight for each pairwise distance was calculated by multiplying the number of MIC values present for that pair of isolates. For the missing values analysis, this could be between 1 and 6, while for isolates with categorical susceptibility data only, this was 0. For example, if isolate 1 has 0 MIC values (+1), and isolate 2 has 6 (+1), the weight for the distance between them would be 7. The +1 was added to each number as the algorithm cannot operate on distances with 0 weight. These values were then squared to generate the relative weighting for each distance.

The map predicted MIC values (as estimated using pairwise distances between points) well under each separate error scheme (Supplementary Table S5). Mean congruence coefficient for the maps was very high for all three analyses (>0.99), meaning inclusion of missing values, additional error or combined AST methods did not strongly affect the overall distribution of isolates on the map (Supplementary Figure S4 and S5). Prediction error, as measured by distance on map, was typically lower than one $\log_2$ MIC unit, showing the maps were able to predict MIC values well across different error types, though this increased depending on the number of error added values. While prediction error was low across the distribution of the maps, error was lowest for isolates with censored values for several drugs (those positioned in the bottom left of the map). Prediction error (distance on map) was lowest when generating maps using a single dimension, but a two-dimensional map also generated strong predictions and had lower stress in the goodness of fit and dimensionality analyses, suggesting two dimensions provided a better representation.

Notably, isolates with additional added noise provided a higher stress contribution to the maps (Supplementary Table S6 and Supplementary Figure S6). We found this difference in relative stress levels could be used to identify isolates which may have been incorrectly measured, as points which contribute the most stress may suggest assay error for a given isolate. In future, researchers could re-measure isolates above a certain stress threshold to ensure their MIC values are correct, ensuring they represent a unique phenotype, or whether this was due to experimental error. Together, these methods provide a method of phenotypic data curation and validation.

**Supplementary Table S5.** Results of error prediction and cross-validation analyses for phenotype map. For the cross-validation analyses, the first 25 of the 100 simulated datasets were used to test predictions. Mean difference in stress per point refers to the mean difference in stress per point (as a %) between unaltered isolates and those with added error.

| Class | Error estimate (for altered isolates only) (n = 100) | | | | | | Mean error by number of altered values (MIC units) | | | | | | Cross-validation (MIC units) for altered isolates only - Mean prediction error (n = 25) | | | |
|---|---|---|---|---|---|---|---|---|---|---|---|---|---|---|---|---|
| | Mean Cong. Coef. | Mean Error (MIC units) | SD (MIC units) | Mean diff. in stress per point (%) | Errors < 1 MIC unit (%) | Errors > 1 MIC unit (%) | 0 | 1 | 2 | 3 | 4 | 5 | 1D | 2D | 3D | 4D |
| **Missing values** | 0.996 | 0.467 | 0.625 | 0.023 | 86.748 | 13.253 | 0.134 | 0.407 | 0.628 | 0.795 | 1.055 | 1.720 | 0.319 | 0.468 | 0.493 | 0.517 |
| **Added noise** | 0.999 | 0.895 | 0.443 | 0.114 | 69.861 | 30.139 | 0.020 | 0.832 | 1.163 | 1.491 | 1.730 | 0.536 | 0.488 | 0.899 | 0.948 | 1.001 |
| **Combining MIC and categorical sensitivity** | 0.999 | 0.320 | 0.681 | -0.03 | 86.535 | 13.465 | *NA* | *NA* | *NA* | *NA* | *NA* | *NA* | 0.232 | 0.314 | 0.35 | 0.355 |

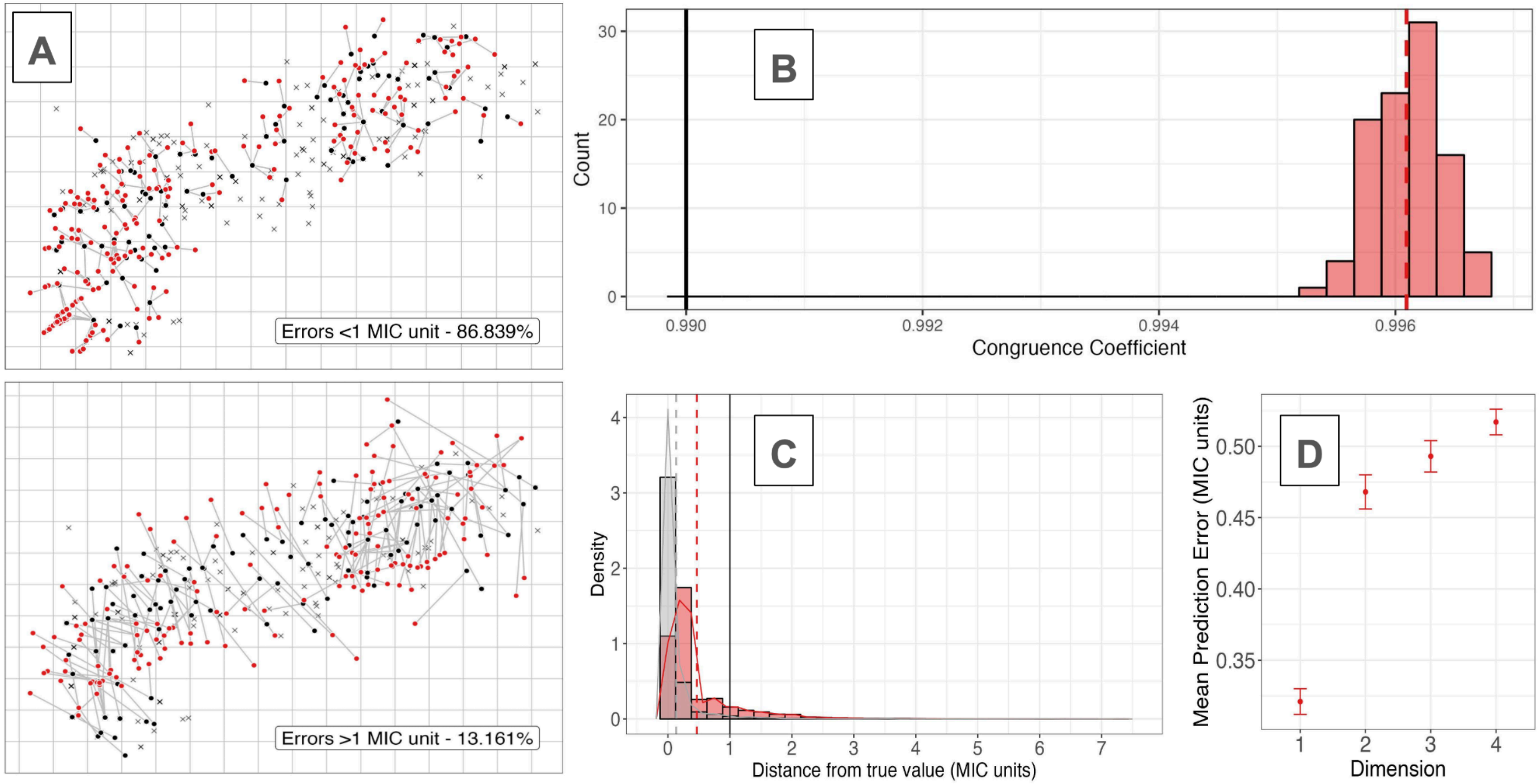


**Supplementary Figure S4.** Missing value cross-validation analysis. Panel A shows one sample dataset with 10% MIC values removed, where the red values represent the predicted locations of each isolate connected to their true location (black points) on the observed map. Labels show proportion of errors across all 100 datasets. Top panel shows prediction errors below 1 MIC unit (86.839%), while the bottom panel shows those above 1 MIC unit (13.161%). Panel B shows the congruence coefficients of the 100 sampled maps after Procrustes rotation and comparison to the true map. Panel C shows the pairwise distances between each isolates true position and their predicted position (across all samples). Red bars show the isolates with MIC values removed while grey shows the remaining isolates. Black, red and grey lines show a reference of 1 MIC unit, mean error of missing value samples, and mean error of all other samples respectively. Panel D shows the distances between an isolates true map position and their predicted positions for the first 25 samples, when maps are made across different dimensions.

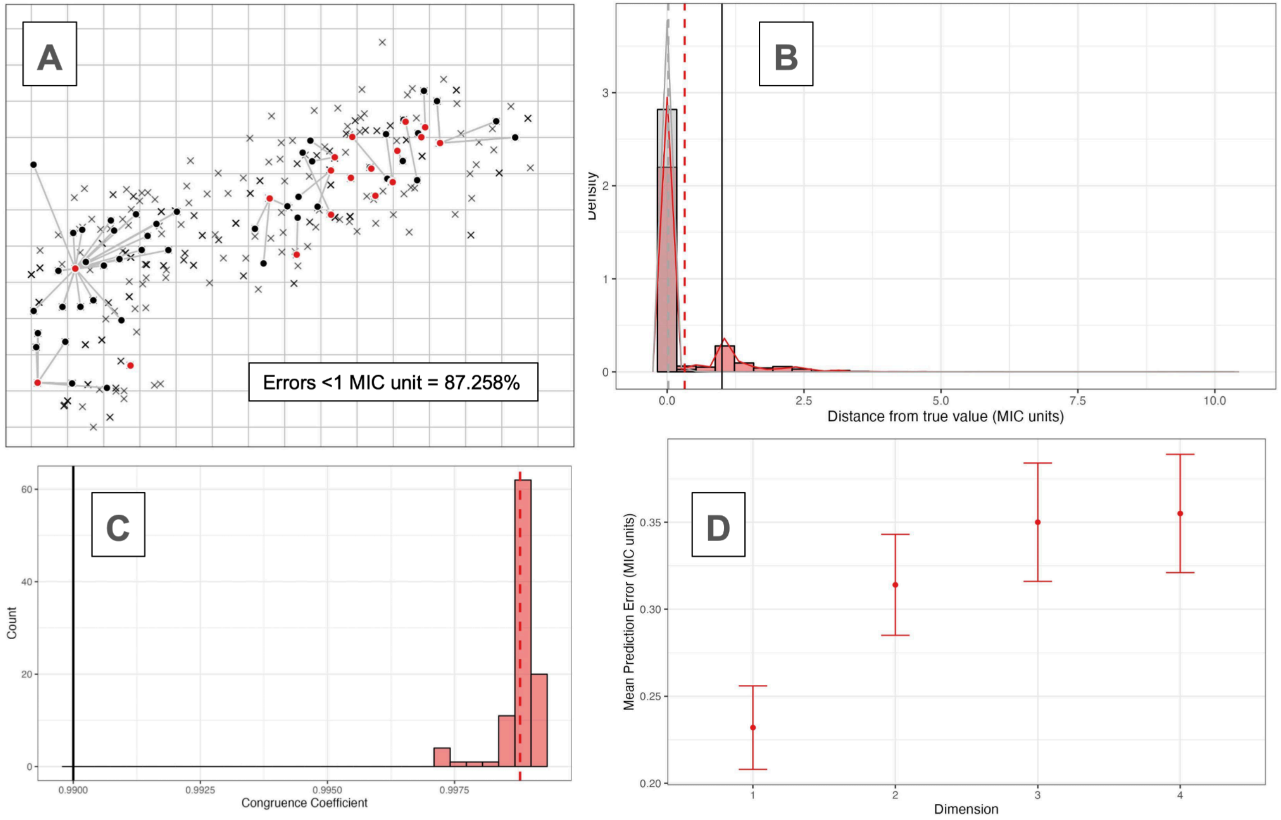


**Supplementary Figure S5.** Combining MIC and categorical sensitivity analysis. Panel A shows one sample dataset with 10% MIC values converted into categorical sensitivity classifications, where the red values represent the predicted locations of each isolate connected to their true location (black points) on the observed map. Labels show proportion of errors across all 100 datasets. Panel B shows the pairwise distances between each isolates true position and their predicted position (across all samples). Red bars show the isolates with MIC values removed while grey shows the remaining isolates. Black, red and grey lines show a reference of 1 MIC unit, mean error of missing value samples, and mean error of all other samples respectively. Panel C shows the congruence coefficients of the 100 sampled maps after Procrustes rotation and comparison to the true map. Panel D shows the distances between an isolates true map position and their predicted positions for the first 25 samples, when maps are made across different dimensions.

**Supplementary Table S6.** Relative increase in stress-per-point contribution for isolates with computationally added error. Note the contribution to overall map stress increased strongly among isolates which had error added to their MIC values, and this increased depending on the number of values which were adjusted. Across the 100 datasets, there were too few isolates with 5 MIC values missing to estimate SE, it is therefore marked as NA (also see Supplementary Figure S6).

| Number of error-adjusted MIC values for a given isolate | Mean change in stress-per-point (%) | Standard error change in stress-per-point (%) |
|---|---|---|
| 0 | -6.188 | 0.01 |
| 1 | 39.79 | 0.467 |
| 2 | 76.472 | 1.266 |
| 3 | 113.066 | 4.651 |
| 4 | 106.554 | 16.446 |
| 5 | 267.604 | *NA* |

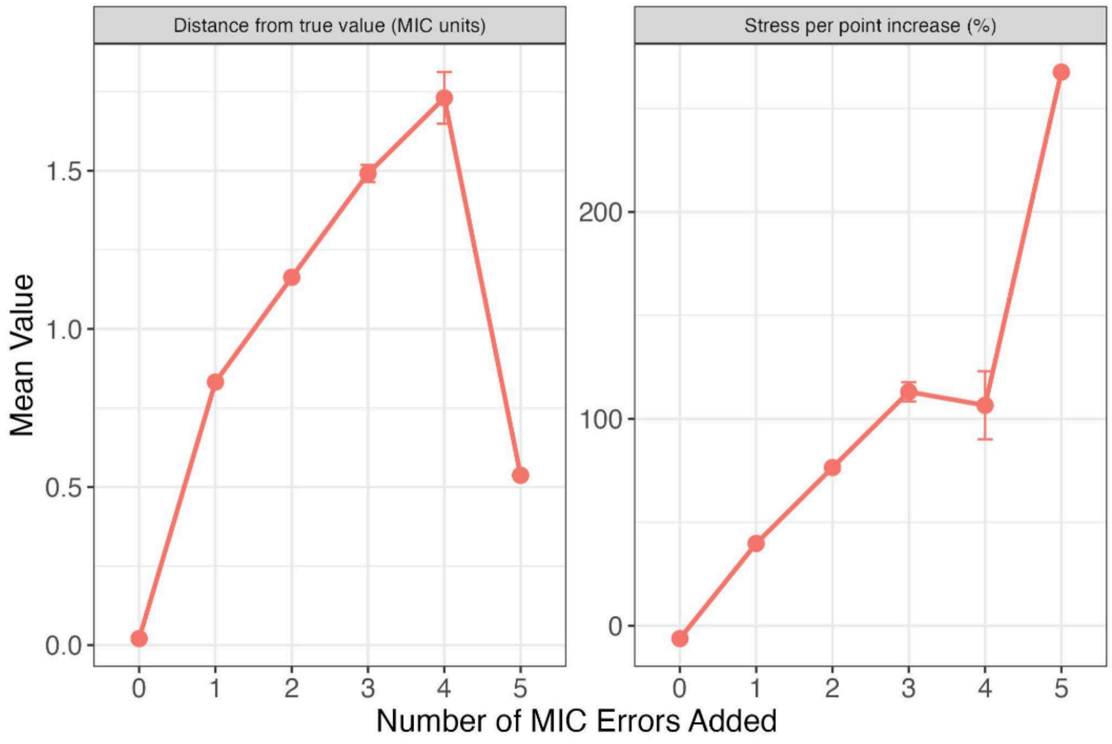


**Supplementary Figure S6.** Estimates of error (left) and stress-per-point (right) increased significantly when errors were added to multiple MIC values for a given isolate (error bars = standard error). The increase in relative stress contribution of isolates with additional error could allow further curation of phenotypes based on the results of the phenotypic map, as researchers could re-measure isolates above a threshold stress contribution to test if MIC results are accurate.

*11. Inclusion of censored values on phenotype maps*

Depending on the dilution range used, some MIC values can be beyond the sensitivity of the assay. For example, if drug concentrations are measured down to a certain dilution, isolates with MIC values lower than this are simply marked as equal to or below a lower censored value (e.g., <= 0.03 µg/mL) (10). The same can be true for an upper censored value (e.g., >8 µg/mL). This complicates downstream analysis, as isolates which vary in their phenotypes are marked as having the same MIC value. This is common in large surveillance datasets, such as the *S. pneumoniae* dataset. Theoretically, the maps can interpret the position of isolates beyond the range of the assay for a given drug, using the information available for the other drugs (Supplementary Figure S1). However, the presence of isolates with censored values for all drugs could still influence the positioning of other isolates on the map. We tested whether this was the case using three methods; firstly, we removed isolates with censored values for all drugs and remade the map. This did not affect the position of the remaining isolates (Congruence coefficient > 0.999) (Supplementary Table S7). Secondly, we removed isolates with fewer than two numerical titres and found that again maps were nearly identical (Congruence coefficient > 0.999). Lastly, we compared metric maps to those made with ordinal and weighted MDS, we then weighted isolates with censored values less strongly (as described in Supplementary Section 1.10). These maps were nearly identical to the original metric maps made without weighting (Congruence coefficient > 0.999). In conclusion, the maps were robust to the inclusion of isolates with several censored MIC values.

**Supplementary Table S7.** Summary statistics for censored MIC value analyses. Note, ordinal MDS only preserves the rank order of pairwise distances, giving the algorithm more flexibility to position isolates with censored MIC values, as long as it is equal to or greater than a threshold distance.

| Class | Error estimate | | |
|---|---|---|---|
| | Congruence Coef. | Mean pairwise dist. | SD pairwise dist. |
| **Ordinal compared to metric MDS** | 0.999 | 0.129 | 0.137 |
| **Removing all isolates with censored values** | 0.999 | 0.457 | 0.604 |

*12. Dimensionality tests*

It is only possible to make phenotype maps because of the strong patterns of covariation between drugs. However, one possibility is this covariation could also be observed under a random permutation of values. If so, the map might not be capturing a real biological signal. Another possibility is that the data could in fact be one-dimensional, and it is simply experimental noise which is being represented in the second dimension on the maps. To test these hypotheses, we generated 100 duplicate datasets and subjected them to different permutation schemes. We then generated a map for each permuted dataset and compared them to maps made with observed data.

- *Random permutation* – MIC values for each drug are independently permuted across isolates, disrupting the covariation structure while maintaining the relative frequency of MIC values. A map was then generated for each permuted dataset, and the resulting stress values are compared to that of the observed data. If the observed MIC values contain real biological structure, the maps constructed from permuted datasets should have significantly higher stress (6,8).
- *One-dimensional* – We generated a series of one-dimensional data with experimental error added. To do this, we took the observed values for a single drug from the dataset (selected at random for each permutation) and used this for each of the columns representing the 'different' drugs (6 in this case). We then added noise (+1/-1 one $\log_2$ dilution) to 10% of the values and tested how well the data could be represented in a single dimension, comparing their stress to a 1-dimensional map made with the observed data.

For the random permutation scheme, the stress values for the permuted column maps were higher than those for the observed data (Supplementary Figure S7), suggesting the observed covariances are unlikely to be due to chance. For the simulated one-dimensional maps in 1D, each map had much lower stress than the observed data represented in a single dimension, suggesting the second dimension is not merely capturing experimental noise. To compare the one- and two-dimensional maps more directly, we superimposed the single dimensional map onto the two-dimensional maps using Procrustes. For both maps, the isolates which had the highest stress per point were those which were furthest away from their position on the 1-dimensional map. The mean distance of each unique phenotype to its position on the 1-dimensional map was above one MIC unit, with many distances over two MIC units (Supplementary Table S8).

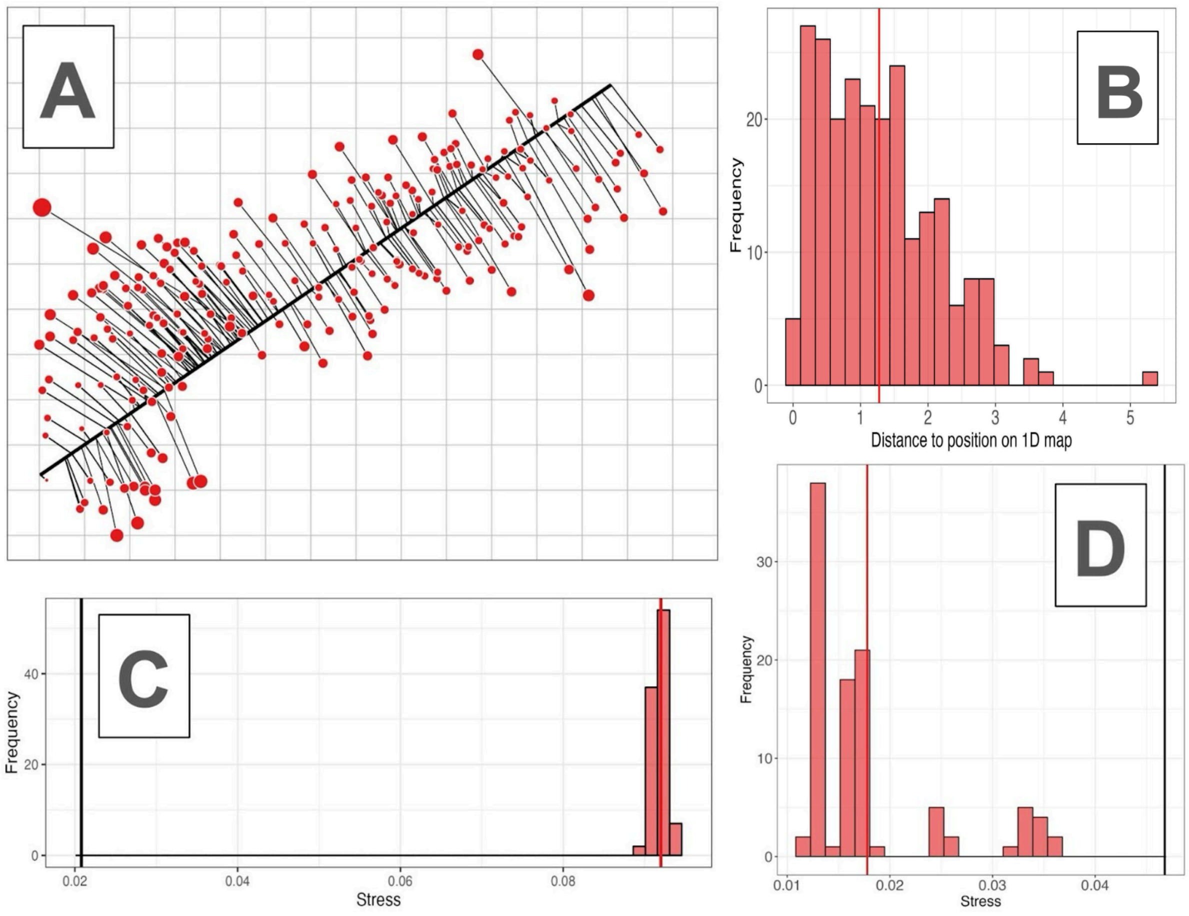


**Supplementary Figure S7.** Permutation analyses. Panel A shows the 1-dimensional map (thick black line) superimposed over the 2-dimensional map using Procrustes rotation. Here, isolate positions on the 2-dimensional map are connected to their position on the 1-dimensional map. The size of the points corresponds to their stress in the 1-dimensional map, with larger points representing those with a larger relative contribution to overall stress. Notably, the isolates further from the line tend to have higher stress. Panel B shows the Euclidean distance between each unique phenotype on the 2-dimensional map and their position on the 1-dimensional map, with the mean value represented as a solid red line. Panel C shows the stress values for the permutation analysis, in which each drug was permuted in the original MIC dataset and the map was remade (n = 100). Here each stress value (mean = solid red line) is above the value of the map made with the observed data (thick black line). Panel D shows stress values of the one-dimensional permutation, in which simulated 1-dimensional datasets were generated (n = 100), with the mean as a thick red line and the observed 1-dimensional map stress in black.

**Supplementary Table S8.** Error estimates in dimensionality analysis.

| | 1D map | Permuting all columns | | One-Dimensional Dataset | |
|---|---|---|---|---|---|
| | Mean Distance (MIC units) | Mean Stress | Mean Percentage Difference (%) | Mean Stress | Mean Percentage Difference (%) |
| **Phenotype map** | 1.28 | 0.092 | +341.26 | 0.018 | -61.952 |

### *Supplementary Section 2 - Genetic analyses in detail*

1. *PBP-types in S. pneumoniae*

In *S. pneumoniae*, three key penicillin binding proteins (*pbp1a, pbp2b, and pbp2x*) have been identified as the major determinants of phenotypic variation in MIC to beta-lactams. We therefore focused on the amino acid sequences of the transpeptidase regions in these proteins for all genotype-phenotype analyses (Supplementary Table S9).

**Supplementary Table S9.** Description of PBP transpeptidase regions. Active site motifs were defined according to (13). Throughout this work, PBP1A, PBP2B, and PBP2X are capitalised to refer specifically to their transpeptidase regions only.

| PBP | Classifier | Metric |
|---|---|---|
| *PBP1A* | Total no. amino acid positions | 277 |
| | Transpeptidase location | 371-647 |
| | Variant/invariant positions | 94/183 |
| | Active sites | S370TMK, S466SN, K557TG |
| *PBP2B* | Total no. amino acid positions | 278 |
| | Transpeptidase location | 379-656 |
| | Variant/invariant positions | 80/198 |
| | Active sites | S386VVK, S443SN, K615TG |
| *PBP2X* | Total no. amino acid positions | 359 |
| | Transpeptidase location | 229-587 |
| | Variant/invariant positions | 111/248 |
| | Active sites | S337TMK, S395SN, K547SG |

2. *Genetic map and clustering*

To relate the genetic maps to the phenotype maps and make further genotype-phenotype comparisons (Supplementary Section 2.7), we classified PBP genotypes using a hierarchical clustering algorithm. Determining the optimal number of clusters in an MDS plot is subjective and depends on the data and the rationale of the study. Here, the aim was to choose a high enough number of clusters to partition PBP-types into groups with relevant differences between them, but not so high that the resulting groups no longer represent useful classifications. We therefore based the number of clusters on the following criteria:

- Interpretability of the clusters - Ultimately, the number of clusters is decided on whether they capture biologically useful information. The groups should have a high enough number of PBP-types, but they should be genotypically distinct enough to make useful comparisons. This was therefore the primary criterion in deciding on the number of clusters.
- Choosing a partition on the dendrogram which splits the centroids into roughly equal sized groups, with large distances between them (Supplementary Figure S9).
- Choosing the 'elbow' on a scree plot - After a certain point, adding additional clusters add little additional information to the clustering. This can be visualised by looking at the decrease in within-group variance between different numbers of clusters.

We emphasised the first of these criteria in deciding on the number of clusters, as the goal of the clustering was to make genotype-phenotype comparisons in Supplementary Section 2.7. Based on this, the optimal number of clusters was therefore between 10-14. The two remaining criteria were then used to refine this selection, for which we chose 12 clusters. This choice of clustering separated the PBP-types into groups with relatively small within-cluster phenotypic distances (Mean = 0.444 MIC units). Although there was overlap between genetic clusters on the phenotype map (Supplementary Figure S8), within-cluster pairwise distances were smaller than between-cluster pairwise distances (Mean = 4.748 MIC units). On a 3D genetic map, the clustering and results of downstream analysis were essentially the same as for the 2D map.

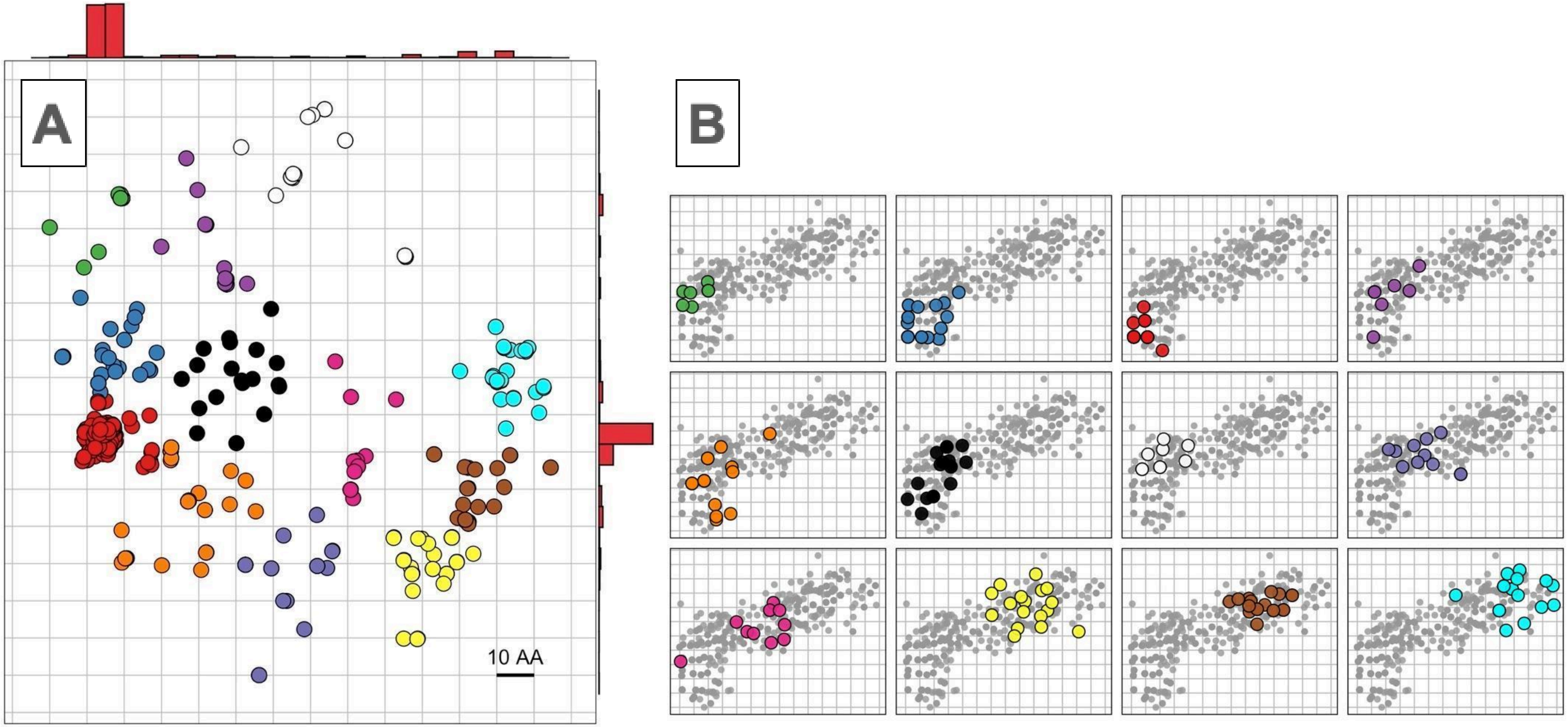


**Supplementary Figure S8.** Panel A shows the genetic map for *S. pneumoniae*. As with the phenotype map, the points represent isolates but the vertical and horizontal axes represent genetic distance, where the gridlines represent 10 amino acids difference in any direction. Notably, many of the most susceptible PBP-types fall on the left of the genotype map (red), while there is a diversity of phenotypes among the other isolates. The hierarchical clustering algorithm split the isolates into 12 groups, based on their position on the genotype map. The positions of these clusters are shown in panel B, where large, coloured points represent the median centroid position of each PBP-type on the phenotype map. Grey points represent the isolates of the other clusters as a reference.

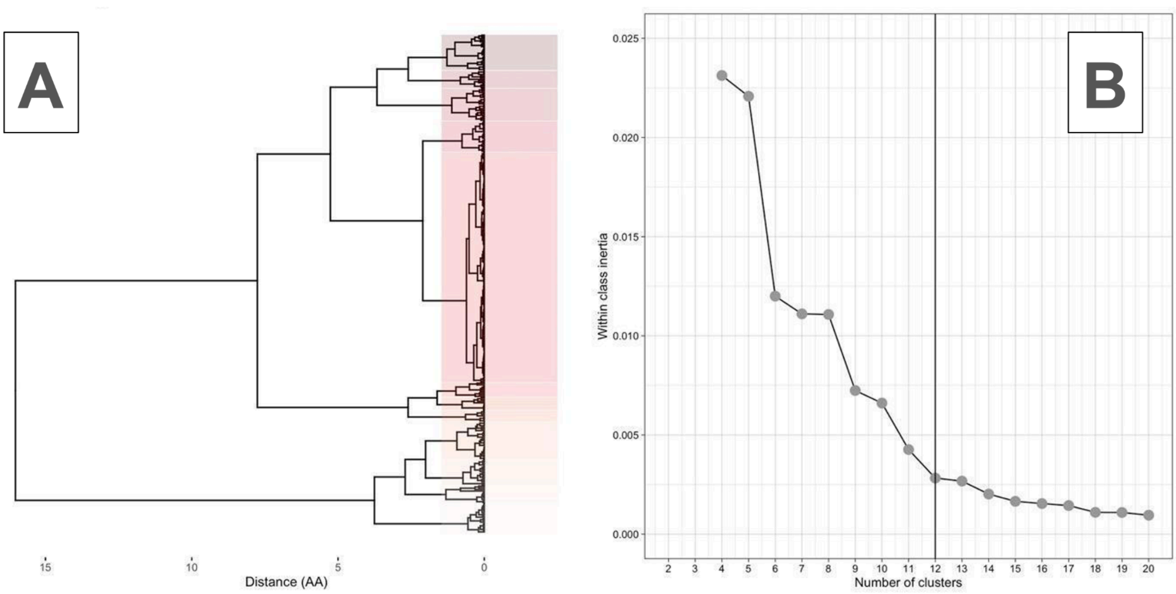


**Supplementary Figure S9.** Choice of clustering for genotype map. A) shows a dendrogram of the pairwise distances between isolates on the genotype map. Boxes indicate the clusters, splitting the dendrogram into 12 clusters. B) shows the within cluster variance for increasing numbers of clusters. The vertical lines show the choice of cluster used for each map, at the inflection point (elbow), beyond which adding additional clusters does not substantially decrease within-cluster variation. Note that although much of the variation could be captured in a relatively small number of clusters (~3), such a small number of clusters was not useful for making genotype-phenotype comparisons between isolates. A larger number of clusters (n = 12) was therefore used.

#### 3. Heritability and per-PBP variance decomposition

To justify focusing solely on these PBP proteins, we quantified the proportion of phenotypic variance in beta-lactam MIC values explained by amino acid variation in the transpeptidase regions. Heritability is the proportion of phenotypic variance in a trait which can be explained by the genetic variation among individuals within a given population, as opposed to environmental factors or experimental error. For all estimates of heritability, we used isolate position on the phenotypic map axes as proxy phenotypes, and amino acid sequence variation rather than genetic sequences. Estimates of heritability can be subdivided into 'broad' and 'narrow' sense heritability ($H^2$ and $h^2$ respectively):

##### 3.1. 'Broad-sense' heritability (H2)

'Broad-sense' heritability ($H^2$) is the proportion of phenotypic variance which is explained by all genetic or amino acid sequence variation in a population, including epistatic, dominance, or interaction effects. Under certain assumptions, $H^2$ can be estimated by calculating the intraclass correlation (repeatability) of a given trait (14). When a phenotype is measured for multiple isolates with the same PBP sequence (PBP-type), repeatability can be measured as:

$$Repeatability = \frac{\sigma_g^2}{(\sigma_g^2 + \sigma_e^2)} \tag{3.1}$$

where:

$$\sigma_g^2 = \frac{(MS(G) - MS(E))}{r} \tag{3.2}$$

and:

$$\sigma_e^2 = MS(E) \tag{3.3}$$

Here, r represents the number of replicates per PBP-type. $MS(G)$ is the mean sum of squares for each PBP-type, while $MS(E)$ is their residual error. $MS(G)$ and $MS(E)$ were both estimated using an analysis of variance. For our calculations, since each PBP-type had a different number of isolates, $r$ was replaced with $\underline{r}$:

$$\underline{r} = (n - 1)^{-1}\left(R_1 - \frac{R_2}{R_1}\right)$$

(3.4)

where:

$$R_1 = \sum r_i \text{ and } R_2 = \sum r_i^2$$

(3.5)

Since isolate MIC values were measured under the same conditions, we would expect differences between them to be due to differences in their PBP amino acid sequences, rather than environmental differences. Where this assumption is met, repeatability is equal to $H^2$. However, given there may be subtle differences in measurements, for example between labs, estimates of $H^2$ can only be taken as an upper limit. $H^2$ estimates were calculated using the 'heritability' package in R v 4.0.4 (15).

*3.2. 'Narrow-sense' heritability (h2)*

In contrast, 'narrow-sense' heritability ($h^2$) is the proportion of phenotypic variance explained by additive genetic variation only. $h^2$ is estimated as the ratio of additive genetic variance to total phenotypic variance and can be estimated using linear mixed models (LMMs). LMMs decompose variation in a phenotype into the variance explained by individual genetic markers, genetic relatedness, and error (see Supplementary Section 2.8). Here, $h^2$ is calculated without the fixed effect of genetic markers and only estimating the other parameters. Variance in a phenotype can be decomposed as:

$$y \sim N\left(0,\ \sigma_g^2 R + \sigma_e^2 I_N\right)$$

(3.6)

where y is the phenotypic vector, $\sigma_g^2$ is the proportion of variance explained by variation in amino acid sequences, and $R$ is the Realised Relatedness Matrix (see below). $\sigma_e^2$ is the proportion of variance explained by residual error, where $I$ is a N x N identity matrix. Using this model, narrow-sense heritability can be estimated as:

$$h^2 = \frac{\hat{\sigma}_g^2}{\left(\hat{\sigma}_g^2 + \hat{\sigma}_e^2\right)}$$

(3.7)

where $\hat{\sigma}_g^2$ and $\hat{\sigma}_e^2$ are the maximum likelihood estimates of $\sigma_g^2$ and $\sigma_e^2$ respectively. Notably, an advantage of LMMs is they can allow for complex genetic covariance structures based on similarity between SNP-based relatedness matrices (or amino acid-based relatedness in this case). Here, the Realised Relatedness Matrix (RRM) is used to estimate relatedness between individuals:

$$R = \frac{1}{S}XX^T$$

(3.8)

S is the number of amino acids used to calculate relatedness, and X is the N x S matrix of standardised sequences, where N is the number of isolates. Framing relatedness in this way allows the similarity between isolate PBP-types to be captured in an N x N matrix, where the similarity between isolates *i* and *j* is contained within the *i*th row of the *j*th column. The Realised Relatedness Matrix was used for all subsequent estimations of relatedness between PBP-types (see Supplementary Section 2.8).

#### 3.3. Per-PBP variance decomposition

The equations above can be used to separate phenotypic variance into that which is explained by particular genes of interest (16). We used this extension to quantify the relative contribution of each PBP protein on position on the phenotypic map axes. Here, $M$ is the multiple sets of amino acids (PBPs in this case) - $\{G_1, ..., G_M\}$, into which to decompose variation. $S_M$ is the number of amino acids in set $m$, and $R_m = \frac{1}{S_m}G_m G_m^T$ is the Realised Relatedness Matrix for a given set of amino acids ($m$). Phenotypic variance can be then decomposed into each set using the following:

$$y \sim N\left(0, \left(\sum_{m=1}^{M} \sigma_m^2 R_m + \sigma_e^2 I_N\right)\right)$$

(3.9)

as

$$h_m^2 = \frac{\hat{\sigma}_m^2}{\sum_{m=1}^{M} \hat{\sigma}_m^2 + \hat{\sigma}_e^2}$$

(3.10)

Where the maximum likelihood estimates of $\sigma_m^2$ and $\sigma_e^2$ are provided by $\hat{\sigma}_m^2$ and $\hat{\sigma}_e^2$ respectively. All variance decomposition and narrow-sense heritability estimates were calculated using the FastLMM/LIMIX framework for LMMs in Python (17,18).

#### 3.4. Heritability results

Estimates of broad-sense heritability were high across both dimensions (95.8-97.6%) (Supplementary Table S10). Narrow-sense heritability estimates were lower than broad-sense heritability, with between 57.8-71.4% of phenotypic variation explained by additive genetic variation for each dimension. Much of the phenotypic variation that was explained by additive genetic variation could be explained by modifications of the PBP2X protein, as indicated by the high variance decomposition values. Relatively little phenotypic variation could be explained by residual noise (< 4% for each map axis).

These estimates suggest much of the variation in beta-lactam resistance phenotype can be explained by amino acid sequence variation in the three PBP proteins. Consequently, we would expect isolates with the same PBP sequence to have the same coordinates on the phenotype map, albeit with a degree of error due to differences in testing. We therefore quantified the phenotypic variance present within PBP-types. There were 310 unique PBP sequences for the *S. pneumoniae data*, of which 138 had more than a single isolate. Median within PBP-type distances were low–typically below than one $\log_2$ MIC unit (0.643 units), suggesting isolates within PBP-types were phenotypically very similar (Supplementary Figure S10). This was the case for 96 of 138 PBP-types in *S. pneumoniae* (69.5%), though one caveat is that many PBP-types had lower censored MIC values for all drugs tested, and therefore showed little to no phenotypic variation, as their MIC values were set at the lower dilution threshold. Some PBP-types showed substantial variation in phenotypes (median phenotypic distance of >2 MIC units). This could be due to experimental error in measurements, incorrect labelling of isolates, or indicate other areas of the genome may play a role in determining beta-lactam MIC.

**Supplementary Table S10.** Heritability and per PBP additive variance decomposition estimate for each map dimension.

| Map dimension | Heritability estimate (%) | | Per PBP additive variance decomposition (%) | | | |
|---|---|---|---|---|---|---|
| | $H^2$ (CI) | $h^2$ | PBP1A | PBP2B | PBP2X | Residual |
| 1 | 97.6 (97.2-97.9) | 71.4 | 11.9 | 11 | 76 | 1.1 |
| 2 | 95.8 (95.2-96.4) | 57.8 | 18.2 | 17.7 | 61.3 | 2.8 |

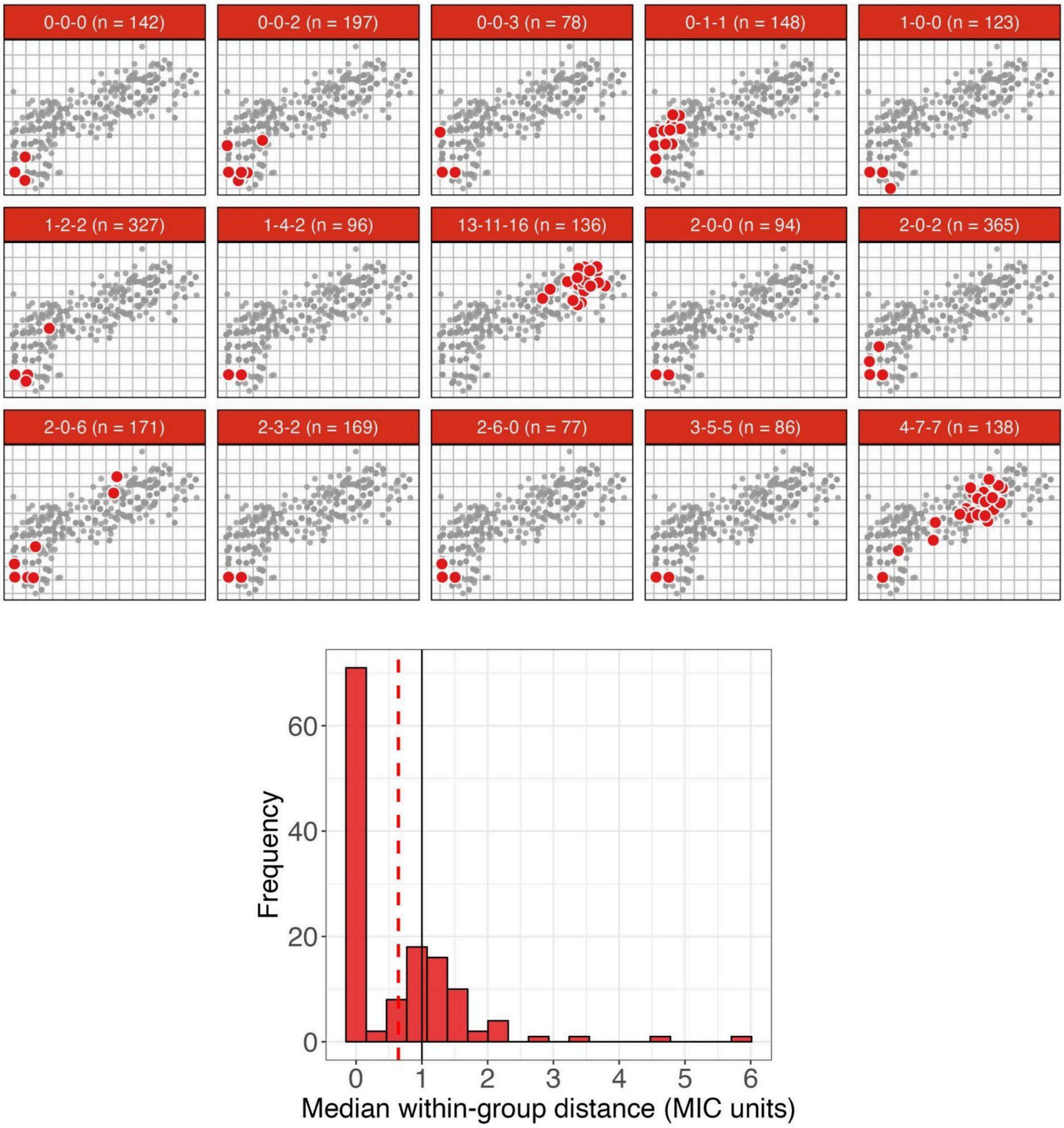


**Supplementary Figure S10.** Top panels show the 15 most common PBP-types are represented visually on the map as large red points, with the distribution of the other isolates marked in grey. Within PBP-type variation is low, however, for many PBP-types, there are several outlier points which are very different in their beta-lactam phenotypes (e. g. PBP-type 2-0-6 and 4-7-7). Bottom panel shows the distribution of median within PBP-type distances (MIC units). Here, the median within PBP-type distance was calculated for each PBP-type with more than one isolate and represented on the histogram. The red dashed and black solid lines represent the mean within-group distance and a reference of one $log_2$ MIC unit respectively.

4. *Strong relationship between number of amino acid changes from susceptible PBP-type and beta-lactam MIC*

Supplementary Figure S11 demonstrates a strong relationship between increased genetic distance from the susceptible representative PBP-type (2-0-2) and increasing MIC (Spearman's rank, rho = 0.791, $p < 0.001$). The cyan, brown, yellow, and pink isolates (clusters 10, 11, 12, and 9) are positioned on the right of both genotype and phenotype maps, highlighting that isolates with the most extensive PBP modifications tend to have the highest MIC values. In contrast, intermediate phenotypes can be caused by a range of different combinations of PBP substitutions (orange - 5, black - 6, blue - 2, green - 1, purple - 4 and white - 7 clusters). Notably, cluster 7 (white) is unique, in that although it is highly divergent from the susceptible PBP-type, it only exhibits a low-level or intermediate increase in MIC (below the penicillin breakpoint). Isolates of this cluster have not increased in MIC to the same level as the clusters 12 (yellow) and 11 (brown), despite a similar number of PBP substitutions. A similar effect is also observed among the green and purple clusters (cluster 1 and 4), which are not as strongly divergent in phenotype as those of the violet/pink clusters (8 and 9), despite similar numbers of amino acids difference.

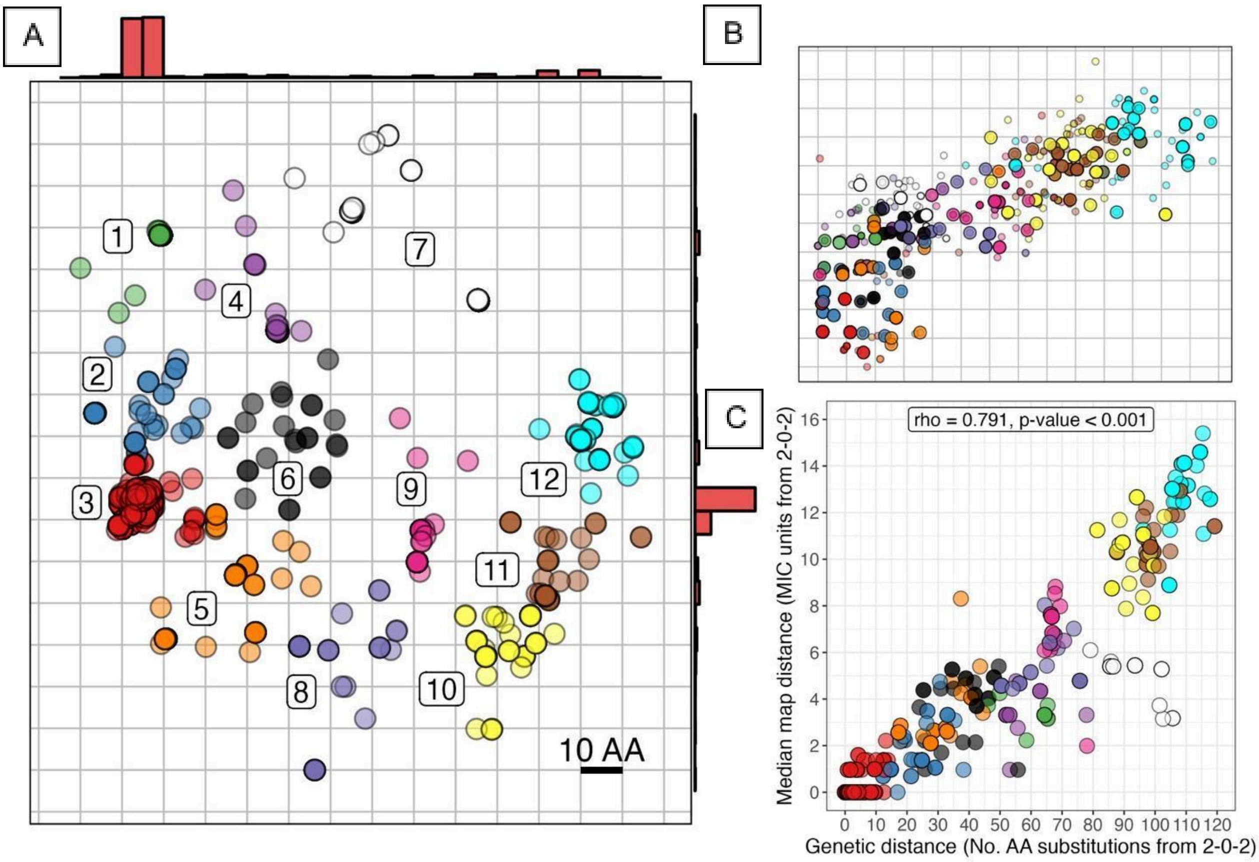


**Supplementary Figure S11.** Comparison of genotype (A) and phenotype (B) maps coloured by the hierarchical clustering algorithm. On the phenotype map, smaller points represent the individual isolates on the phenotype map, while large points represent the median centroid position for each PBP-type. Panel C) shows the relationship between genetic distance (amino acid substitutions) and median phenotypic distance of each PBP-type (MIC units) from the most common susceptible PBP-type. For example, cluster 7 (white) is unique, in that although it is highly divergent from in PBP-type, it only exhibits a low-level or intermediate increase in MIC (below the penicillin breakpoint). Importantly, isolates of this cluster have not increased in MIC to the same level as clusters 10 and 11 (yellow and brown respectively), despite a comparable total number of PBP substitutions. A similar effect is also observed among clusters 1 and 4 (green and purple), which are not as strongly divergent in phenotype as those of the clusters 8 and 9 (violet and pink), despite similar numbers of amino acids difference.

*4.2 Phenotypic and genetic diversity within MLSTs*

Several MLSTs displayed similar phenotypes (see Figure 2 - Main text), indicated by the low pairwise phenotypic distances between their PBP-types (Supplementary Table S11 and Figure S12). For example, PBP-types within MLST 1840 (n=26) exhibited a mean phenotypic distance of 1.02 MIC units, indicating limited phenotypic variation. Similarly, PBP-types within MLST 695 (n=60) were positioned closely together on the map (mean pairwise distance = 1.68 MIC units), suggesting relatively uniform resistance profiles. By contrast, isolates of MLSTs 63, 156, and 1092 had larger mean within-lineage phenotypic distances, typically above 2 MIC units, with isolates extending across clinically relevant breakpoints. Moreover, some MLSTs harbor numerous unique PBP-types (e.g., MLST 63 with 21), potentially explaining their broader phenotypic range.

Genotypic divergence from the most common sensitive PBP-type (2-0-2) varied widely between MLSTs. For example, isolates of MLSTs 156, 1451 and 320 had diverged extensively in both phenotype and genotype from sensitive PBP-types. In contrast, MLST 338 and 3280 showed substantial amino acid divergence in PBPs (71.5 and 93.5, respectively) yet only moderate increases in MIC - 4.52 and 5.50 MIC units relative to 2-0-2. This was reflected in a high phenotypic/genetic distance ratio for these MLST types, of 15.83 and 16.99 respectively. MLST 63 and 199 showed the highest genetic variation among PBP-types, with a mean pairwise genetic distance of 46.67 and 30.99 respectively.

**Supplementary Table S11.** Phenotypic and genetic diversity for the 12 most common MLST types. Here, we report the average distance of each MLST from the most common sensitive PBP-type (2-0-2) in both phenotype (MIC units) and genotype (amino acid changes), as well as the within-MLST mean and standard deviation (SD) of pairwise distances.

| MLST (no. isolates) | Unique PBP-types | Phenotypic Distance (MIC Units) | | | | | Genetic Distance (AA Changes) | | | Phenotypic Distance /Genetic Distance |
|---|---|---|---|---|---|---|---|---|---|---|
| | | Distance from 2-0-2 | | | Pairwise within MLST | | Distance from 2-0-2 | Pairwise within MLST | | |
| | | Euclidean | X Axis | Y Axis | Mean | SD | Mean | Mean | SD | |
| **156 (n = 22)** | 5 | 10.23 | 8.32 | 5.75 | 2.74 | 1.41 | 82.43 | 27.1 | 13.64 | 8.05 |
| **1451 (n = 20)** | 2 | 13.21 | 10.96 | 7.36 | 0.86 | NA | 108.04 | 9.23 | NA | 8.18 |
| **320 (n = 132)** | 9 | 12.81 | 10.93 | 6.56 | 2.62 | 1.65 | 108.87 | 6.72 | 3.86 | 8.5 |
| **1092 (n = 15)** | 5 | 8.75 | 7.11 | 5.09 | 2.33 | 1.14 | 77.75 | 20.42 | 13.42 | 8.88 |
| **558 (n = 131)** | 8 | 9.43 | 7.69 | 5.24 | 3.21 | 3.81 | 95.73 | 11.14 | 15.26 | 10.15 |
| **199 (n = 114)** | 15 | 2.05 | 1.37 | 1.34 | 2.82 | 3.11 | 20.87 | 30.99 | 33.53 | 10.2 |
| **63 (n = 77)** | 21 | 4.45 | 3.03 | 2.98 | 2.97 | 1.69 | 53.53 | 46.67 | 23.66 | 12.04 |
| **1840 (n = 26)** | 5 | 0.68 | 0.3 | 0.57 | 1.02 | 0.77 | 10.42 | 18.21 | 10.31 | 15.37 |
| **338 (n = 118)** | 7 | 4.52 | 2.25 | 3.85 | 1.8 | 0.98 | 71.53 | 25.35 | 16.42 | 15.83 |
| **3280 (n = 28)** | 1 | 5.5 | 3.45 | 4.26 | NA | NA | 93.52 | NA | NA | 16.99 |
| **695 (n = 60)** | 2 | 0.86 | 0.31 | 0.78 | 1.68 | NA | 14.83 | 28.63 | NA | 17.22 |
| **1373 (n = 50)** | 2 | 3.16 | 0.5 | 3.08 | 0.85 | NA | 64.85 | 1.14 | NA | 20.5 |

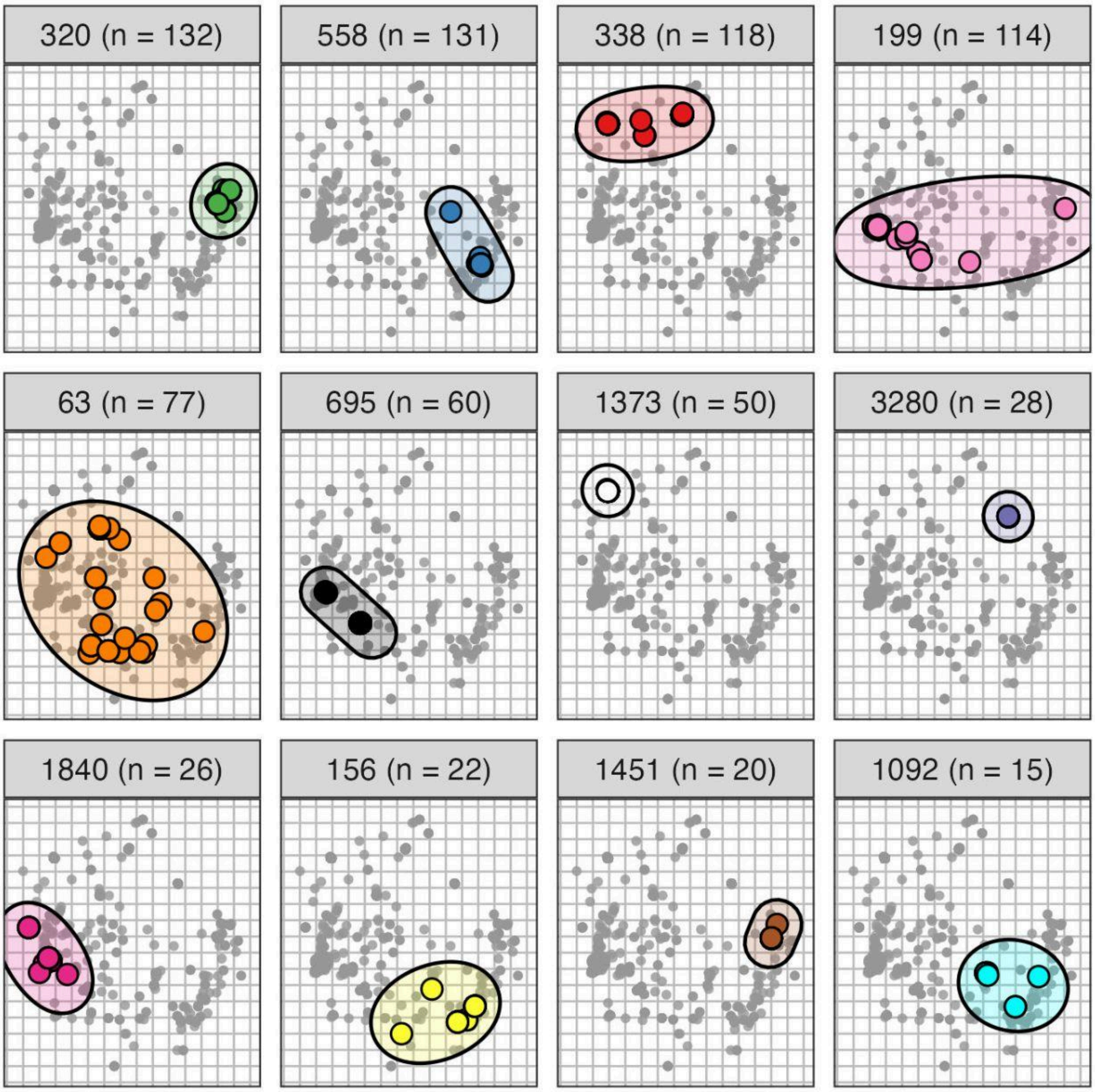


**Supplementary Figure S12.** Coloured points highlight the 12 most common MLSTs on the genetic map, while smaller grey points indicate isolates of the remaining MLST types. Each MLST type is shown as a separate panel, where coloured regions highlight the range of locations for each unique PBP sequence. Within MLSTs, most isolates tend to have similar PBP-types, however MLST 63 and 199 are outliers, as both displayed extensive diversity in PBP sequence. This was also reflected in their phenotypes, as isolates of these MLSTs displayed a range of susceptible and intermediate MIC values. Notably, three MLST types had extensive modification of their PBP-types but only exhibited low-to-intermediate MIC (338, 1373 and 3280). PBP-types within these MLSTs were divergent not only from the most common sensitive PBP-type (2-0-2), but also differed extensively in PBP-type from isolates of other MLST types (e.g. 320, 1451 and 1092).

#### 5. Identification of causal amino acid substitutions

After quantifying the phenotypic variation explained by variation in PBP amino acid sequences, we used a series of methods to identify which substitutions are associated with a change in phenotypic position on the beta-lactam maps. All of these methods were originally developed to identify causal substitutions underlying antigenic change in H3N2 influenza and modified for use in this context (4,12).

#### 6. Identifying pairs of isolates that differ by a single PBP substitution

The first method involves identifying pairs of isolates which vary at a single amino acid position within one of the three PBP proteins. For example, isolate A (XY<u>Z</u>) and B (XY<u>A</u>) have the same genetic background (XY#) but differ by a single substitution, in this case Z/A (Supplementary Figure S13). The Euclidean distance between the two isolates on the phenotype map is then calculated and credited to the variation at this position. In many cases, there may be several isolates of each comparison within a collection. Where there was more than one isolate of a given genotype, we calculated the pairwise phenotypic distance between each isolate in the comparison and took the mean value. In some cases, the same substitution can occur on multiple genetic backgrounds, which allows comparison of the effects of a substitution on different genetic backgrounds in the PBP proteins.

In the *S. pneumoniae* dataset, there were 189 single substitution comparisons, at 88 of the 914 PBP amino acid positions. 15 out of 189 (~8%) of these showed a phenotypic effect above one $\log_2$ MIC unit, occurring at 14 positions in the PBP proteins (1 in 1A, 5 in 2B and 8 in 2X). However, in several cases, isolates in the comparison overlapped in their distribution of phenotypes or had low sample sizes (Supplementary Figure S14). Notably, there were 23 examples of the same amino acid changes occurring on different genetic backgrounds. For example, a substitution of V591I in the 2B protein caused different phenotypic effects on different genetic backgrounds, causing a distance of 0.965 $\log_2$ MIC units on one background, but 0.011 on the other (Supplementary Figure S14 and S15).

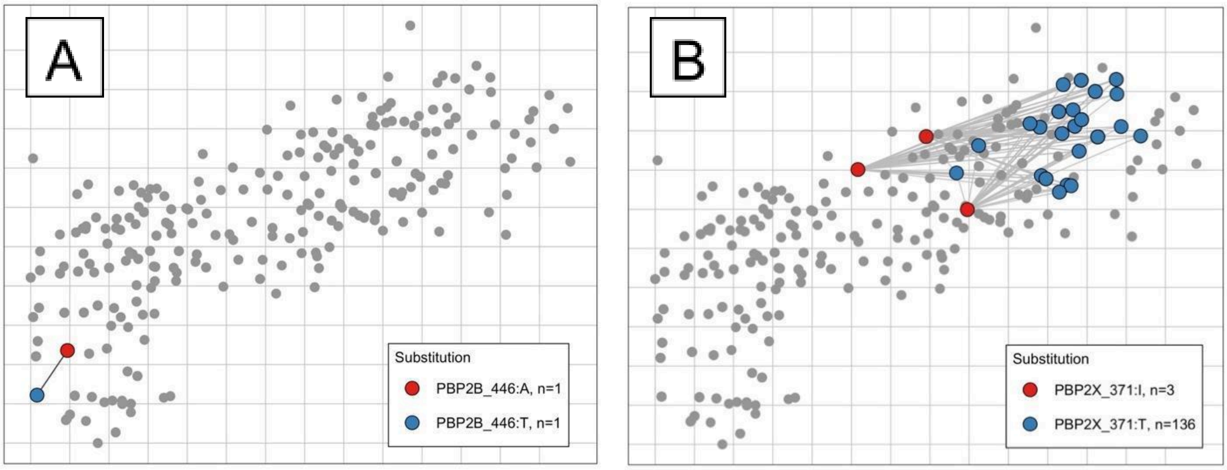


**Supplementary Figure S13.** Example of identifying isolates that vary by a single PBP substitution. A) Here, two isolates have exactly the same PBP genetic background, but differ by a single substitution in the 2B protein, in this case at location T446A. B) Where there were several isolates of each comparison within a collection, the pairwise distances between all isolates in the comparison are calculated and the mean taken.

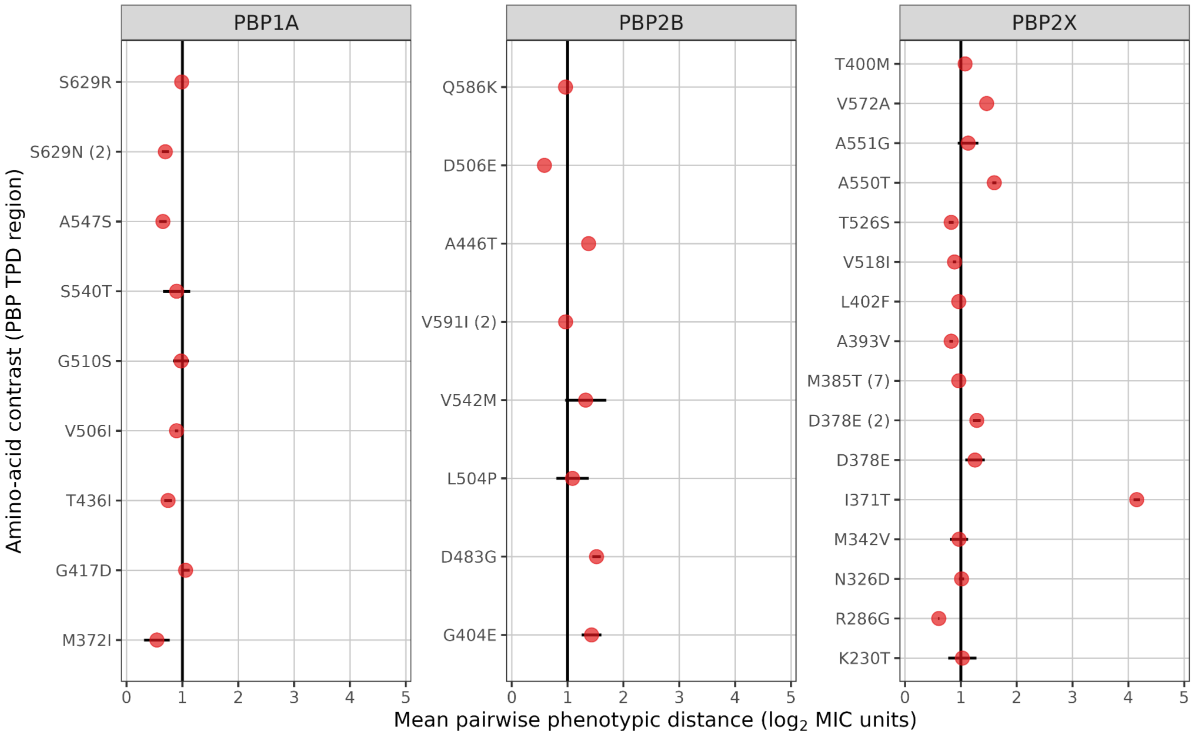


**Supplementary Figure S14.** Mean pairwise phenotypic distances for single-amino-acid comparisons in the PBP transpeptidase regions. Points show the mean phenotypic-map distance for each comparison. Where the same amino-acid contrast occurred on more than one PBP genetic background, entries are numbered (e.g. PBP2B V591I (2) denotes the second such background). Horizontal black bars show the mean ± standard error where estimable; bars are omitted for four comparisons based on a single pairwise distance. The vertical black line marks one log2 MIC unit. Only comparisons with a mean phenotypic distance >0.5 log2 MIC units are shown.

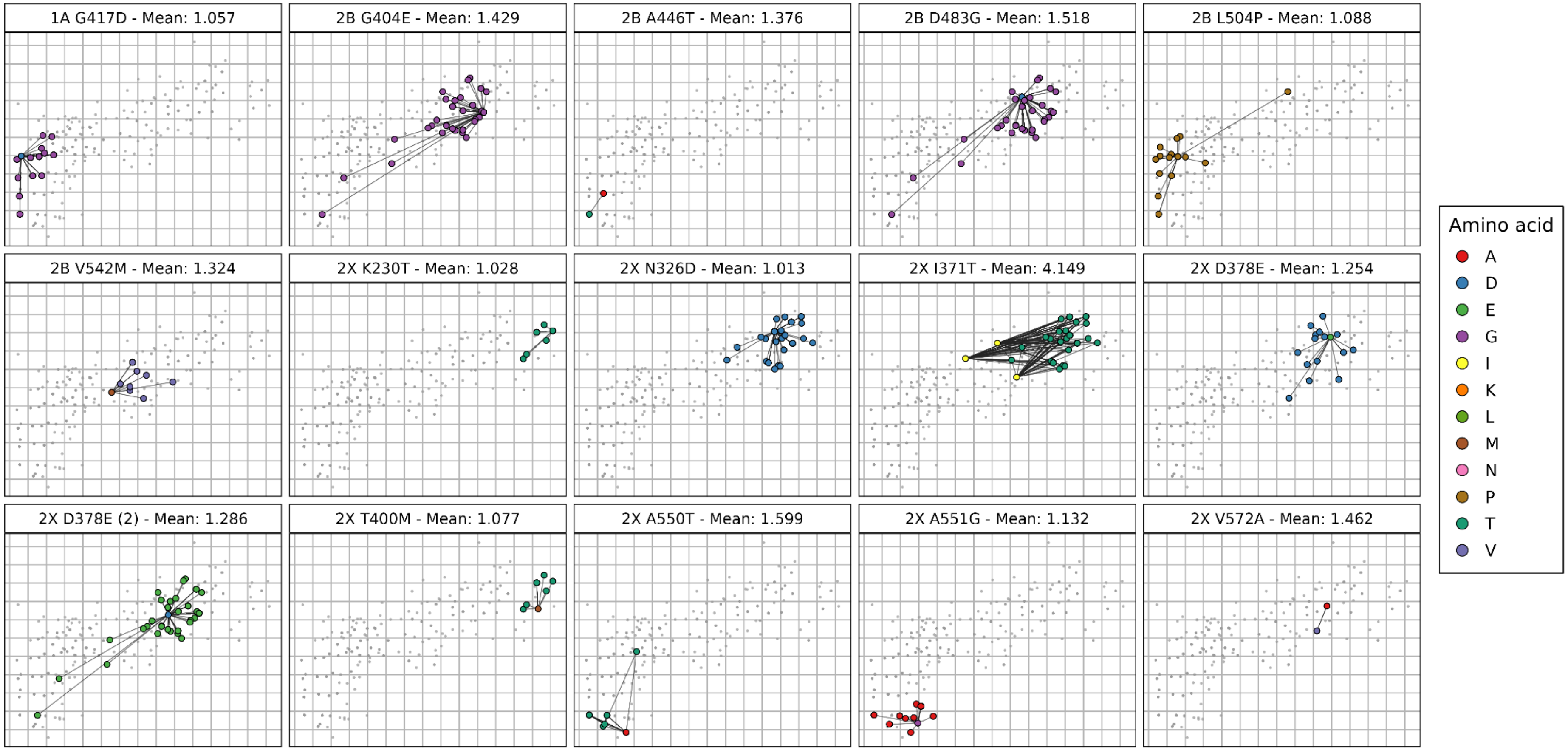


**Supplementary Figure S15. Single-amino-acid PBP comparisons visualised on the phenotype map.** Each panel shows a comparison with a mean pairwise phenotypic distance of at least one log2 MIC unit. Large coloured points represent isolates belonging to the two PBP groups, which differ at the indicated amino-acid position; grey points represent the remaining isolates. Black lines connect isolates from the opposing comparison groups to illustrate their pairwise phenotypic distances. Panel labels report the mean pairwise phenotypic-distance estimate. Where the same amino-acid contrast occurred on more than one PBP genetic background, the additional background is numbered, as for PBP2X D378E (2). Several comparisons show overlapping phenotype distributions or are based on small groups, notably PBP2B A446T and PBP2X V572A.

### 7. *Identifying 'cluster-difference' substitutions*

A second method to identify causal substitutions involves clustering of isolates based on their phenotype and genotype and identifying 'cluster-difference' substitutions (4,12). While this method was originally developed in the context of antigenic cartography, it needed to be modified for use here. In the original H3N2 antigenic map, strains of influenza were clearly clustered into groups of phenotypically distinct strains. The K-means clustering algorithm was then used to pick out groups, and used to identify amino acid substitutions which underlie phenotypic change between clusters. In the case of the phenotype maps for *S. pneumoniae*, isolates do not obviously fall into discrete clusters, and show considerably higher genetic, yet lower phenotypic diversity than the original antigenic maps. For this reason, it was necessary to split PBP-types into less diverse subsets.

We first subdivided isolates into groups with relatively low PBP genetic diversity using the hierarchical clustering algorithm (as outlined in Supplementary Section 2.2). Within each genetic cluster, the following method was used to identify causal substitutions among several varying positions. Firstly, median centroid positions were calculated for each PBP-type on the phenotype map. Clusters of these centroid positions were delineated using the hierarchical clustering algorithm. The number of clusters for each subset was chosen based on the same criteria used for the full genotype map (Supplementary Section 2.2). 'Cluster-difference' variants were identified under the following definition: they were a variant at a given PBP location, where all PBP-types of cluster A have amino acid X at that position, and all PBP-types of cluster B have a different amino acid at that position. A substitution was marked as 'strong-evidence' of generating an effect if the above was true. In contrast, a variant was considered as having 'weak evidence' if the above was true, but where this was based on a low sample size (5 or fewer PBP-types/less than 3 isolates for each type), or in some cases where there was not full segregation of a given substitution between phenotypic clusters. In this case, 'Strong' evidence indicates an amino acid change likely has an effect, whereas 'Weak' evidence only indicates a change potentially having an effect–and therefore requires further investigation.

For *S. pneumoniae*, this method picked out 103 out of 914 positions as having phenotypic effects. Of these, 20 showed 'strong evidence' of being involved in phenotypic change, with 3 substitutions showing evidence of generating phenotypic effects across several genetic backgrounds (Supplementary Figure S16 and S17). A further 83 positions showed some limited evidence of generating a phenotypic effect ('Weak') based on low sample sizes or incomplete segregation of substitutions between phenotypic groups.

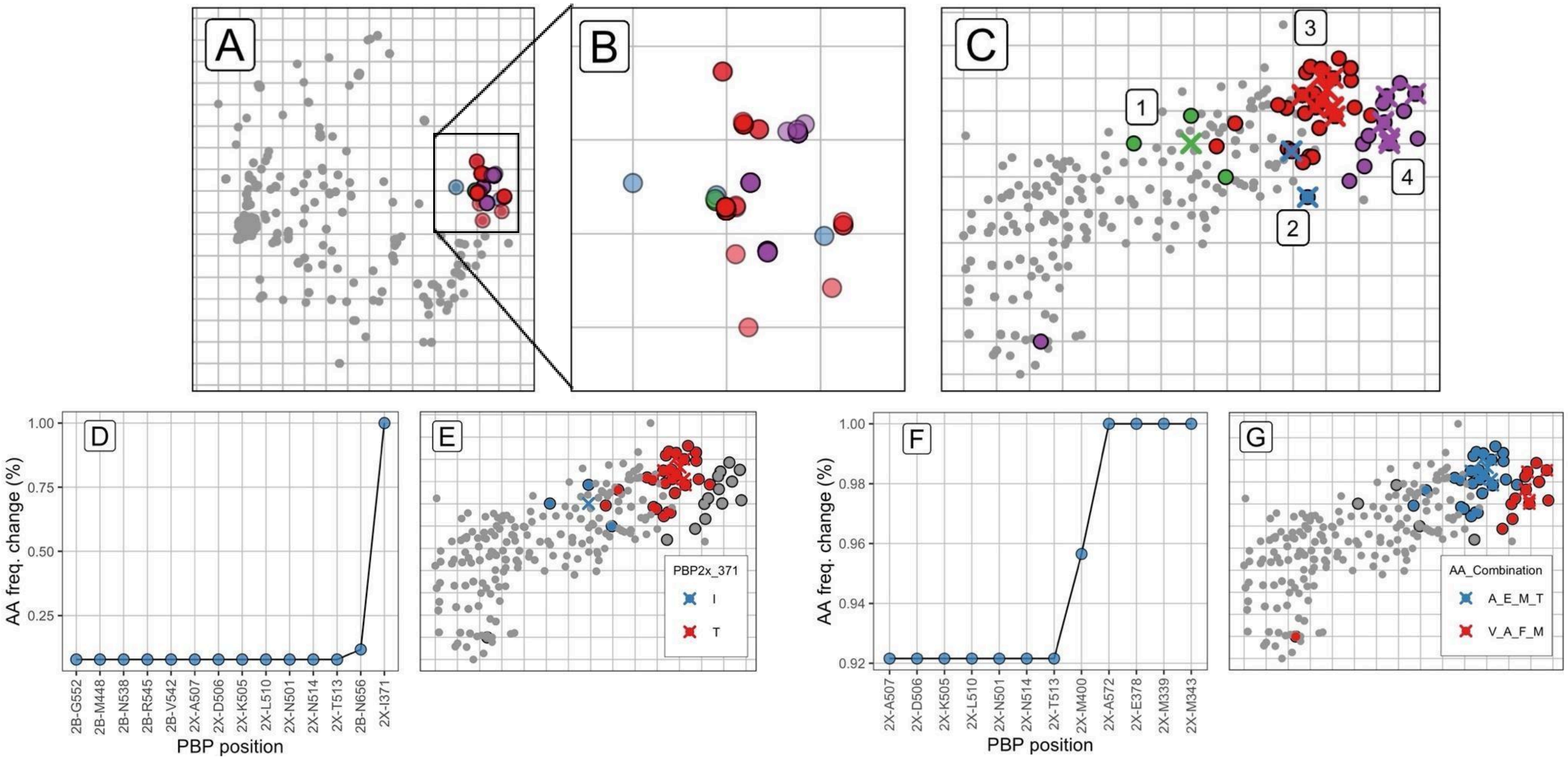


**Supplementary Figure S16.** Example of clustering method for cluster 12. Panels A, B and C show isolates of cluster 12 on the genetic map (A and B), and phenotype map (C), coloured by clustering. The PBP-type centroids (crosses) were split into four phenotypic clusters using the hierarchical clustering algorithm (C). Panels D-G show two examples of the method. On plots D and F, each PBP position is plotted on the X axis. The difference in relative frequency of each amino acid between clusters was plotted on the y axis. Plots E and G show the 1 and 4 PBP positions that were identified in comparisons (clusters 1-3, D/E and 3-4, F/G) plotted on the phenotype map.

| PBP | Loci | No. genetic clusters |
|---|---|---|
| 2X | N514 | 3 |
| 2X | A572 | 2 |
| 2X | D506 | 2 |
| 2X | A369 | 1 |
| 2X | A491 | 1 |
| 2X | A507 | 1 |
| 2X | E320 | 1 |
| 2X | E378 | 1 |
| 2X | I371 | 1 |
| 2X | I462 | 1 |
| 2X | K505 | 1 |
| 2X | L510 | 1 |
| 2X | L565 | 1 |
| 2X | M339 | 1 |
| 2X | M343 | 1 |
| 2X | N501 | 1 |
| 2X | T490 | 1 |
| 2X | T513 | 1 |
| 2X | V423 | 1 |
| 1A | V386 | 1 |

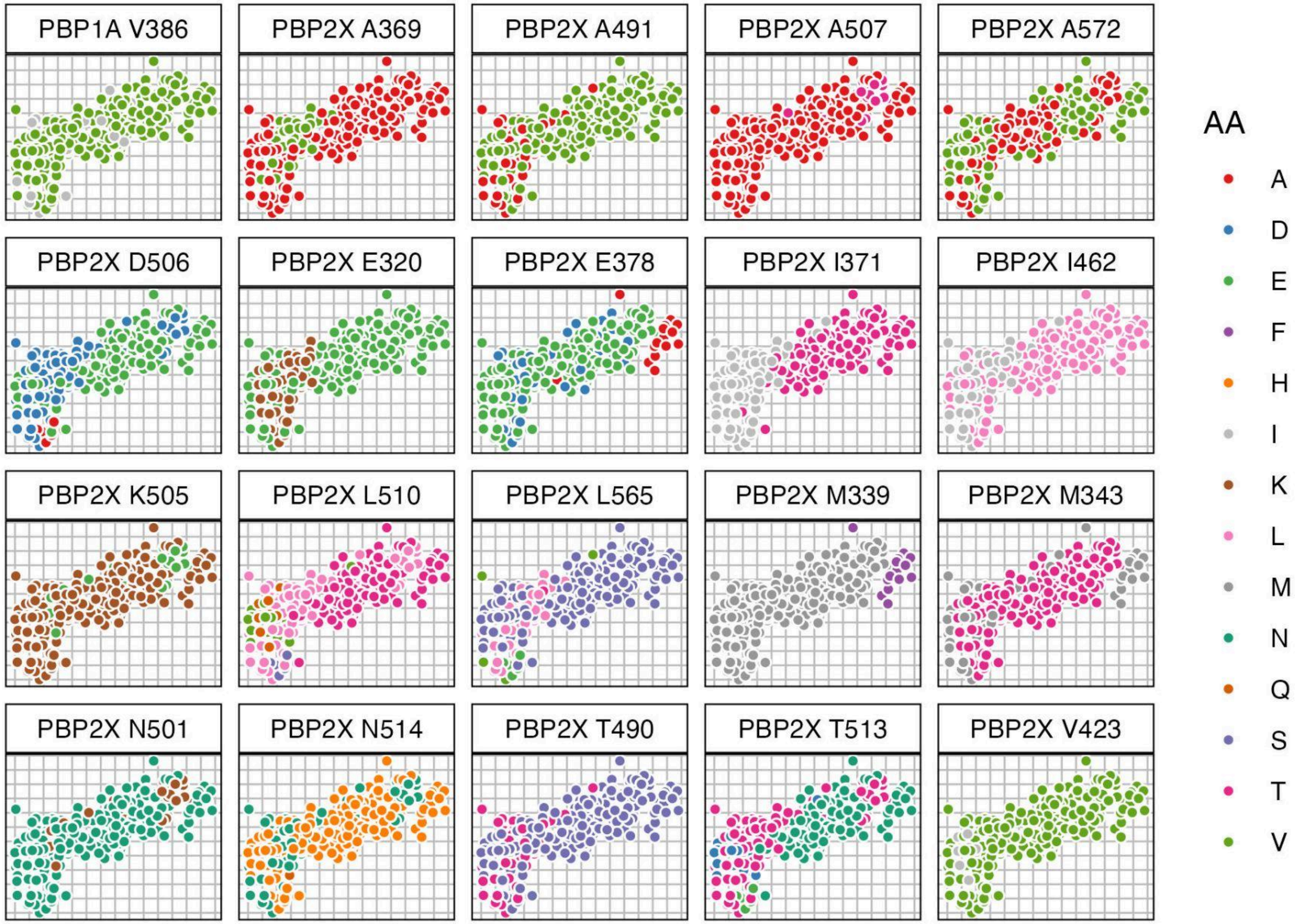


**Supplementary Figure S17.** 20 loci identified by clustering as having strong evidence of generating phenotypic change, coloured by amino acid. Three loci were associated with phenotypic change in several genetic clusters (PBP2X N514, A572, and D506).

*8. Multivariate linear mixed models (LMMs)*

The third method involves applying linear mixed models to the map axes (4,12). Linear mixed models are a well-established method of identifying causal genetic variants responsible for phenotypic variation. Importantly, these methods provide a framework to statistically test the effect of a particular substitution, while accounting for the similarity we would expect between isolates simply because they have similar PBP sequences. LMMs decompose variation in a phenotype (*y*) into several components, modelled as both fixed and random effects:

$$y = \beta x + g + \psi \tag{3.11}$$

Firstly, this includes the effects of a single amino acid or genetic marker of interest ($\beta x$), modelled as a fixed effect. Here, $\beta$ is the effect size, and $x$ is the presence/absence of the genetic marker being tested (0 or 1). Secondly, the relatedness between isolates, modelled as a random effect ($g$). This captures the phenotypic similarity we would expect between isolates based on how similar their PBP proteins are. $g$ is modelled as a normal distribution, with mean 0 and covariance $\sigma_g^2$:

$$g \sim N\left(0,\ \sigma_g^2 R\right) \tag{3.12}$$

This represents the similarity between isolates in their PBP-type, where *R* represents the relatedness RRM (see Equation (3.8)). Lastly, experimental noise is also included as a random effect ($\psi$), and is modelled as a normal distribution with mean 0, and variance $\sigma_e^2$:

$$\psi \sim N\left(0,\ \sigma_e^2 I_N\right) \tag{3.13}$$

Here, *I* represents the N x N identity matrix. Statistical testing is then made possible by comparing the null model $\mathcal{H}_0$ ($\beta$ = 0), the marker having no effect, against the full model $\mathcal{H}_1$, that the genetic marker has an effect ($\beta \neq 0$):

$$\mathcal{H}1 : y \sim N(0 + x\beta + g + \psi)$$
$$\mathcal{H}0 : y \sim N(0 + g + \psi)$$

(3.14)

The likelihood ratio ($D$) is used to compare the likelihood of the two models. Here, $D$ is defined as:

$$D = L\left(\beta, \hat{g}, \hat{\sigma}_e\right) / L\left(\underline{g}, \underline{\sigma}_e\right)$$

(3.15)

where $L\left(\beta, \hat{g}, \hat{\sigma}_e\right)$ is the likelihood of the parameters of the full model $\mathcal{H}_1$, while $L\left(\underline{g}, \underline{\sigma}_e\right)$ is the likelihood given the parameters of the null model, $\mathcal{H}_0$. Since $2D$ is equivalent to a $\chi^2$ distribution, it can be used to calculate a p-value. The univariate LMM can then be extended to include multiple phenotypes ($Y$), where $Y$ is a matrix of $(N, P)$ phenotypes, and β is an effect size vector of length $P$ :

$$Y = x\beta^T + g + \psi$$
$$g \sim MN_{N,P}\left(0, \sigma_g^2 R, C_g\right)$$
$$\psi \sim MN_{N,P}\left(0, \sigma_e^2 I_N, C_N\right)$$

(3.16)

$g$ is now modelled as a multivariate matrix distribution, where $R$ and $C_g$ are the genetic individual-to-individual and genetic trait-to-trait covariance matrices respectively. ψ captures residual error as a multivariate matrix-distribution $I_N$, and $C_N$ is the environmental trait-to-trait covariance matrix. All LMMs were implemented using the FastLMM/LIMIX package in Python (17,18).

9. *Epistatic interactions between PBP loci*

While several methods have been developed to investigate epistatic interactions in GWAS, including in the context of variation in beta-lactam MIC in *S. pneumoniae* (19–21), these methods are often 'phenotype-blind' and do not estimate the effect sizes of significant interactions. Since this analysis focuses on only a small portion of the *S. pneumoniae* genome (<1000 amino acids across the PBP transpeptidase regions), it is possible to test all pairwise interactions between loci and estimate their effect sizes using mvLMMs. A modification of the mvLMM framework was originally proposed to test for epistatic interactions in the context of antigenic cartography for H3N2 influenza (12). We used this modification to test for second-order epistatic interactions between PBP loci. In each case, an interaction variable is coded as the logical 'and' between two dummy variables (Supplementary Table S12).

**Supplementary Table S12.** Coding a pairwise interaction between two variables (adapted from 12)

| Isolate | Locus A | Locus B | Loci A&B |
|---|---|---|---|
| a | 1 | 1 | 1 |
| b | 1 | 0 | 0 |
| c | 0 | 1 | 0 |
| d | 0 | 0 | 0 |

The LMM is then run on each interaction variable to test whether it is associated with an effect. Importantly, this modification tests for the presence of an interaction effect independently of the additive effects of each substitution, by including each original marker as covariates:

$$y = x_A \beta_A^T + x_B \beta_B^T + x_{AB} \beta_{AB}^T + g + \psi \tag{3.18}$$

Here, locus A, locus B, and loci A & B are each included as a fixed effect. For each test, both markers were removed from the data and the relatedness matrix was recalculated. Low frequency interactions (i. e. those occurring in less than 1% of the dataset) were excluded from the analysis. A mvLMM was then used to test each interaction effect, recomputing the random genetic and error components for each test, as described in Supplementary Section 2.8. As before, the null hypothesis is competed against the alternative model, by comparing the likelihood ratio of the two models to derive a p-value.

*10. Increasing the statistical power of mvLMMs*

Several computational methods have been applied to identify causal substitutions in the PBPs underlying beta-lactam resistance in streptococci. However, the low overlap in causal substitutions identified between methods may be partly due to low statistical power. We therefore used a series of modifications to maximise the statistical power of mvLMMs to detect effects, based on similar implementations developed for antigenic cartography (12).

*10.1. Focus on the transpeptidase regions of the PBP proteins only*

As the transpeptidase regions of the PBP proteins have been repeatedly identified as responsible for the majority of variation in beta-lactam phenotypes, we apply mvLMMs only to these regions. These regions (<1000 amino acid positions) are several orders of magnitude smaller than the entire streptococci genome, where genome size can be up to 2.5 Mb. By focusing on these small regions, this massively decreases the number of statistical tests which need to be conducted and means less strict correction for multiple comparisons can be applied.

*10.2. Applying mvLMMs to phenotype map axes*

We used MDS axes as proxy phenotypes rather than raw MIC values individually, as this increases statistical power in several ways. Firstly, by using fewer statistical tests in total, this allows less strict correction for multiple comparisons, increasing statistical power. Secondly, effect sizes are larger in joint dimensions than for either dimension separately, providing additional power to detect effects (12). Thirdly, combining multiple correlated phenotypes controls for these correlations when identifying significant substitutions and allows for comparison of joint and specific effects. Lastly, the dimensionality reduction step reduces experimental noise and allows post-hoc visualisation of mvLMM results.

*10.3. Amino acid sequences rather than genetic sequences*

A third way of increasing statistical power to detect effects is to use amino acid sequences rather than genetic sequences (SNPs). Amino acid sequences are shorter than the genetic sequences, and therefore require fewer statistical tests and less severe controlling for multiple comparisons. One caveat is that since an amino acid at a given locus can be any of 20 amino acids, this may actually increase the total number of statistical tests, as a test of association would be required for each amino acid at a given position. However, such a high degree of variation at a single position is very uncommon, meaning practically, fewer tests in total are required compared to using the genetic sequences. In all cases where the RRM was calculated (as in Equation 3.8), the PBP amino acid sequences were used rather than a standard SNP matrix.

#### *10.4. Dummy variables*

A fourth method to increase statistical power involves recoding the amino acid at each locus as a 'dummy' variable, in the form of a presence/absence matrix of amino acids at each position. Recoding protein sequences as dummy variables in this way can be used to substantially reduce the total number of tests (12). For example, effect sizes induced by a given substitution are mirrored by a change back to the original amino acid. Therefore, only changes away from a representative 'susceptible' PBP genotype need to be included. The most common susceptible PBP-type was defined as the reference with which to compare. For the *S. pneumoniae* dataset, this was the representative ‘2-0-2’. This PBP-type was the most common PBP in the dataset with the lowest MIC values to all six drugs. In each case, the amino acid belonging to the ‘2-0-2’ PBP-type was removed from each locus. This generated a set of dummy variables reflecting changes away from this susceptible genotype. Furthermore, variables which are identical or ‘precisely inverse’ can be combined together, as these positions do not code additional information relative to their mirrored position (12). By removing these variables, far fewer tests are required, requiring less strict correction for multiple comparisons, and increasing statistical power. Supplementary Table S13 describes the conversion of a PBP molecular alignment to a set of dummy variables.

#### *10.5. Per-test variance decomposition*

LMMs decompose variance in a phenotypic trait into genetic and error components (Equations 3.12 and 3.13), to separate these from the fixed effects associated with a genetic marker. Typically, these components are reused for each association test to reduce computational burden (12). However, where tests are only completed on a small number of positions, it is possible to recompute these parameters for each individual test, providing additional power to detect effects. This is useful here, as some PBP substitutions are known to generate larger phenotypic effects relative to others (13). For each test, the given substitution was removed from the dummy variable matrix and the RRM was recalculated without this substitution. A mvLMM was then used to test each substitution individually, recomputing the random genetic and error components for each test.

#### *10.6. Correction of p-values for multiple comparisons*

Correction of p-values for multiple comparisons is important in GWAS due to the high number of tests on correlated genotypes. One commonly used method of p-value correction in GWAS is the ‘Bonferroni’ correction. This correction scales the derived p-values by the total number of tests conducted (i. e. each marker tested), under the assumption that each test is independent under the null hypothesis. However, when conducting tests on correlated genotypes, this assumption is often not valid in practice, and the correction risks missing causal variants. This is particularly important in the context of recombination in the PBP proteins, as there are strong patterns of linkage disequilibrium between loci. One way of avoiding this overcorrection is by correcting for the 'effective' number of tests, given the correlation between genetic markers, rather than the total number of tests. One proposed method is the Galwey method (12,23). This method uses the eigenvalues of the marker correlation matrix to estimate the effective number of tests ($M_{eff}$):

$$M_{eff} = \frac{\left(\sum_{i=1}^{M} \sqrt{\lambda_i}\right)^2}{\sum_{i=1}^{M} \lambda_i} \tag{3.17}$$

where $\lambda$ are the eigenvalues of the amino acid marker correlation matrix. The p-values are then scaled by the effective number of tests, with p-values greater than one set to one.

#### *10.7. Setting conservative p-value thresholds*

As with all statistical tests, conducting multiple mvLMMs increases the probability of false positives. To help set a conservative p-value threshold, we repeated the mvLMM analysis 100 times, each time permuting phenotypes (map dimensions), while keeping the structure of the PBP sequences the same. This has the effect of breaking the relationships between genotype and phenotype, meaning any significant associations found using the random data are invalid. The lowest p-value found across the 100 random permutations was used as a threshold to help interpret the real data.

**Supplementary Table S13.** Recoding a molecular alignment of amino acids (AA) to dummy variables. A) Original molecular alignment. B) Molecular alignment from (A) re-coded as a presence-absence matrix of dummy variables, resulting in twelve statistical tests. C) Amino acids belonging to the most common susceptible representative PBP-type (ref) were removed from (B). Invariant positions such as AA_6H are therefore also removed (resulting in six statistical tests). D) Identical or 'precisely inverse' positions were then merged, resulting in fewer statistical tests (four independent tests). Table adapted from (12).

A

| Isolate | AA_1 | AA_2 | AA_3 | AA_4 | AA_5 | AA_6 |
|---|---|---|---|---|---|---|
| a (ref) | D | T | K | A | S | H |
| b | D | T | T | E | F | H |
| c | E | I | T | E | G | H |
| d | E | I | K | A | S | H |

B

| Isolate | AA_1D | AA_1E | AA_2T | AA_2I | AA_3K | AA_3T | AA_4A | AA_4E | AA_5S | AA_5F | AA_5G | AA_6H |
|---|---|---|---|---|---|---|---|---|---|---|---|---|
| a (ref) | 1 | 0 | 1 | 0 | 1 | 0 | 1 | 0 | 1 | 0 | 0 | 1 |
| b | 1 | 0 | 1 | 0 | 0 | 1 | 0 | 1 | 0 | 1 | 0 | 1 |
| c | 0 | 1 | 0 | 1 | 0 | 1 | 0 | 1 | 0 | 0 | 1 | 1 |
| d | 0 | 1 | 0 | 1 | 1 | 0 | 1 | 0 | 1 | 0 | 0 | 1 |

C

| Isolate | AA_1E | AA_2I | AA_3T | AA_4E | AA_5F | AA_5G |
|---|---|---|---|---|---|---|
| a (ref) | 0 | 0 | 0 | 0 | 0 | 0 |
| b | 0 | 0 | 1 | 1 | 1 | 0 |
| c | 1 | 1 | 1 | 1 | 0 | 1 |
| d | 1 | 1 | 0 | 0 | 0 | 0 |

D

| Isolate | AA_1E\|AA_2I | AA_3T\|AA_4E | AA_5F | AA_5G |
|---|---|---|---|---|
| a (ref) | 0 | 0 | 0 | 0 |
| b | 0 | 1 | 1 | 0 |
| c | 1 | 1 | 0 | 1 |
| d | 1 | 0 | 0 | 0 |

*11. mvLMMs identify many PBP substitutions with additive and epistatic effects*

There were 914 positions across the three PBP transpeptidase regions, 285 of which varied. After removing low frequency amino acid variants (those occurring in less than 1% of all samples - 36.2 isolates), coding the remaining positions as dummy variables resulted in 170 independent tests. 95 of these were associated with phenotypic change exceeding one $\log_2$ MIC unit, after correction for multiple comparisons ($p < 0.05$). After randomly permuting phenotypes 100 times, and rerunning the mvLMM on each permuted dataset, the lowest p-value found was $p = 1.42 \times 10^{-4}$. 92 of the 95 identified changes were significant below this threshold (Supplementary Table S14 and Supplementary File 1). 27 were located in PBP1A, 26 in PBP2B, and 39 in PBP2X. 18 changes had very large overall joint effect sizes (over two MIC units). Notably, many changes with the lowest p-values had positive effects on the y map axis (associated with penicillin), but negative effects on the x-axis (associated with cephalosporins), in particular those located in PBP2B (e. g. G467L/S469N/G483A/A490S, L609S, P568S, and V503I) (see Supplementary Section 3.1).

We then coded interaction terms between these 170 independent changes. After excluding interactions which had a frequency of less than 1% of the dataset (< 36.2 isolates), 4052 interactions remained. 3147 of these interactions had a significant phenotypic effect after correction for multiple comparisons ($p < 1.68 \times 10^{-3}$, Supplementary Figure S18). Although significant, some identified interactions had large standard error estimates. After excluding interactions which had a lower bound standard error estimate below one $\log_2$ MIC unit, 1926 interactions remained. These interactions involved changes at a total 159 of the 285 variant PBP locations (55.8%). Notably, 39 substitutions were found to have significant interactions with more than 40 other positions, with many of the most densely connected changes located within the 1A (35.9%) and 2X (48.7%) proteins.

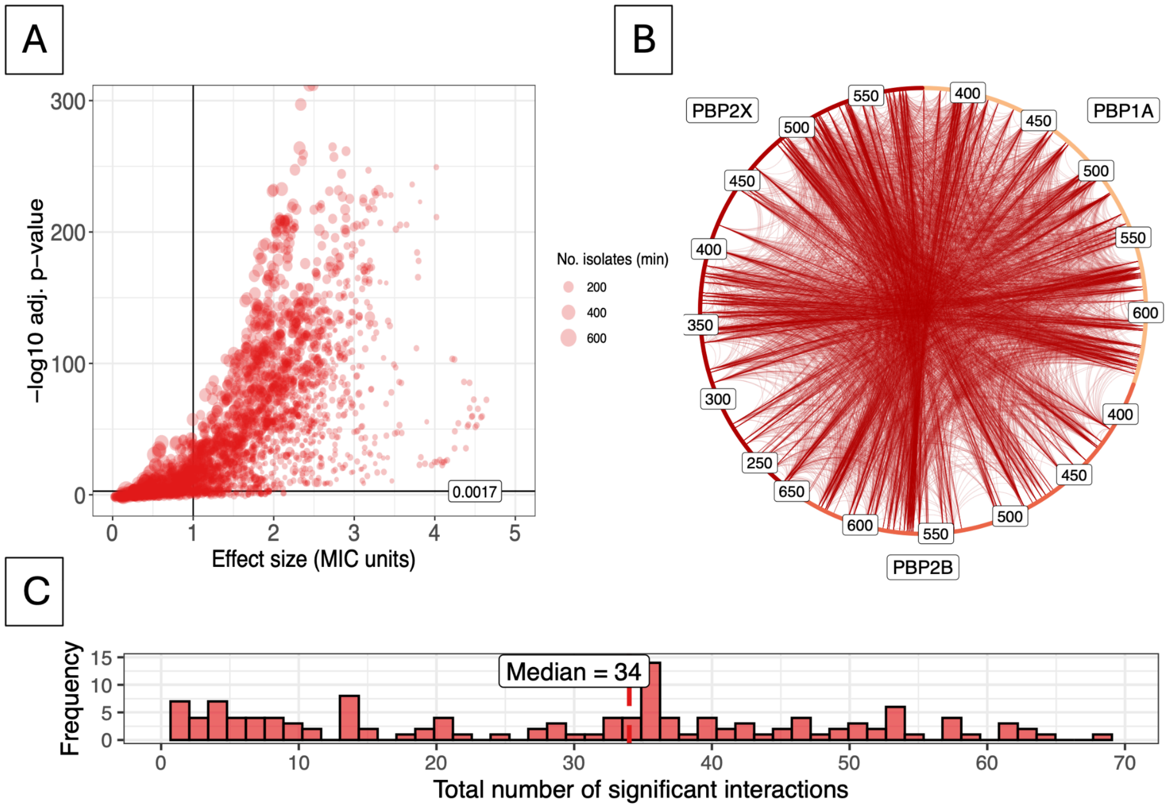


**Supplementary Figure S18.** Epistatic interaction LMM for *S. pneumoniae*. A) Effect size (MIC units) against -log10 adj. p-value. Red points represent a given interaction between two substitutions. Black vertical and horizontal lines represent a reference of one log2 MIC unit and p-value cut-offs respectively. Size of points represents the minimum number of isolates present for that interaction of two substitutions, with larger points representing larger numbers of isolates. B) Network of significant interactions between PBP positions. Each node represents a given substitution, with connected nodes representing a significant interaction between two substitutions. Nodes are coloured by PBP. The size of each node represents the number of significant interactions that substitution was found to have. All significant interactions are shown in the network ($p < 1.68 \times 10^{-3}$ level), with edge thickness representing -log10 adj. p-value. C) Histogram of significant interactions for each substitution. Here, the x-axis represents the total number of significant interactions found for each substitution. The y-axis represents how many substitutions had each number of significant interactions.

*12. Ranking PBP substitutions based on strength of evidence for phenotypic effect*

We summarised the strength of evidence for each substitution by counting the number of methods which indicated a phenotypic effect. For example, if all four methods confirmed an effect for a given substitution, it was marked as 4, for 'Very Strong' evidence. Ratings were then categorised as follows: 3 – 'Strong', 2- 'Moderate', 1 – 'Weak', and 0 – 'No Evidence'. There was very strong to weak evidence for 215 amino acid substitutions having phenotypic effects on beta-lactam MIC, at 179 locations (62.8% of variant positions) (Supplementary Table S14). 117 of the 215 amino acid substitutions had evidence from multiple methods, with one substitution (2X-I371T) identified in all four analyses. These 117 substitutions occurred at 108 unique locations in the PBP proteins (PBP1A - 25, PBP2B - 42, PBP2X - 41). Of the 117 component-expanded substitutions supported by two or more methods, 108 (92.3%) had both a significant additive mvLMM association (Galwey-adjusted $p \leq 0.00397$, without applying the ≥1 effect-size evidence gate) and support from at least one epistatic interaction.

**Supplementary Table S14.** PBP substitutions associated with changes in beta-lactam MIC (only the first 20 substitutions with the largest effect sizes are shown). The strength of evidence for each substitution is categorised by the number of methods identifying a phenotypic effect: 4 methods - ‘Very Strong’ evidence, 3 – ‘Strong’, 2 - ‘Moderate’, 1 – ‘Weak’, and 0 – ‘No Evidence’. '-' indicates an absence of evidence for a given method, such as positions without single substitution comparisons (full table in Supplementary File 1). Sig. mvLMM/uvLMM denotes whether a substitution was significant in the mvLMM on map axes, and uvLMM tests on individual MIC values respectively, with the number of drugs which were significantly associated in brackets.

| Evidence | PBP | Amino acid | Single Subs. Comparison | | | Clustering | mvLMM | Adj. p-value | Effect Size Axis 1 | Effect Size Axis 2 | Epistatic LMM | Sig. mvLMM/uvLMM (No. drugs) |
|---|---|---|---|---|---|---|---|---|---|---|---|---|
| | | | Phenotypic Distance | SE | No. isolates | Confidence | Joint Effect size | | | | No. Sig. interactions | |
| Very Strong | 2X | I371T | 4.149 | 0.058 | 3/136 | Strong | 3.187 | <1e-16 | 2.85 | 1.425 | 40 | Yes/Yes (6) |
| Strong | 2X | E320K | - | - | -/- | Strong | 1.368 | <1e-16 | -0.854 | 1.069 | 18 | Yes/No (0) |
| | 2X | A491V | - | - | -/- | Strong | 1.354 | <1e-16 | 0.713 | 1.151 | 21 | Yes/Yes (1) |
| | 2X | T490S | - | - | -/- | Strong | 1.341 | <1e-16 | 0.707 | 1.139 | 21 | Yes/Yes (2) |
| | 2X | A369V | - | - | -/- | Strong | 1.313 | <1e-16 | -0.251 | 1.288 | 9 | Yes/Yes (1) |
| | 2X | M343T | - | - | -/- | Strong | 1.246 | <1e-16 | 0.53 | 1.128 | 21 | Yes/Yes (1) |
| Moderate | 2X | N444S | - | - | -/- | - | 2.741 | <1e-16 | 2.468 | 1.192 | 36 | Yes/Yes (1) |
| | 2X | S531Y | - | - | -/- | - | 2.445 | <1e-16 | 2.215 | 1.036 | 35 | Yes/No (0) |
| | 1A | T371S | - | - | -/- | Weak | 2.362 | <1e-16 | 2.198 | 0.867 | 8 | Yes/Yes (2) |
| | 2X | L364F | - | - | -/- | Weak | 2.221 | <1e-16 | 2.036 | 0.888 | 34 | Yes/Yes (2) |
| | 1A | N443D | - | - | -/- | Weak | 2.214 | 1.055e-06 | 1.764 | 1.338 | 64 | Yes/Yes (1) |
| | 1A | L611F | - | - | -/- | Weak | 2.189 | 3.829e-04 | 1.881 | 1.12 | 39 | Yes/Yes (1) |
| | 2B | T489A | - | - | -/- | Weak | 2.161 | <1e-16 | 1.414 | 1.633 | 51 | Yes/Yes (3) |
| | 1A | L606I | - | - | -/- | Weak | 2.16 | 7.770e-04 | 1.85 | 1.115 | 39 | Yes/Yes (1) |
| | 2X | V523L | - | - | -/- | Weak | 2.094 | 1.980e-09 | 1.844 | 0.992 | 45 | Yes/No (0) |
| | 1A | G414V | - | - | -/- | Weak | 2.09 | 6.235e-07 | 1.864 | 0.944 | 7 | Yes/Yes (1) |
| | 2B | L609S | - | - | -/- | - | 2.053 | <1e-16 | -1.681 | 1.178 | 5 | Yes/No (0) |
| | 2B | A533G | - | - | -/- | - | 2.052 | <1e-16 | -1.687 | 1.169 | 4 | Yes/No (0) |
| | 2B | D555E | - | - | -/- | - | 2.052 | <1e-16 | -1.687 | 1.169 | 4 | Yes/No (0) |
| | 2B | G597A | - | - | -/- | - | 2.052 | <1e-16 | -1.687 | 1.169 | 4 | Yes/No (0) |

### *Supplementary Section 3 - Additional Analyses*

1. *Many identified substitutions have contrasting effects on different subclasses of beta-lactams*

We assessed whether substitutions were associated with common or specific effects across different subclasses of drugs. A total of 134 substitutions and 3147 epistatic interactions were significantly associated with phenotypic change (any effect size) (Supplementary Figure S19). 262 (7.99%) had positive effects on penicillin (≥1 $\log_2$ MIC unit), while 866 (26.39%) had positive effects on cephalosporins. 35 substitutions/interactions (1.07%) had positive effects on both map dimensions, such as PBP2X-I371T, which showed evidence of generating a phenotypic effect in all four analyses. 1603 substitutions/interactions (48.86%) generated negative effects on penicillin MIC, almost all of which were epistatic interactions, while only 336 (10.24%) generated negative effects on cephalosporin MIC. Notably, several substitutions in PBP2B generated positive effects on penicillin MIC, but a negative effect on cephalosporin MIC (e.g. P568S, L609S and G467L/S469N/G483A/A490S) (Supplementary File 1).

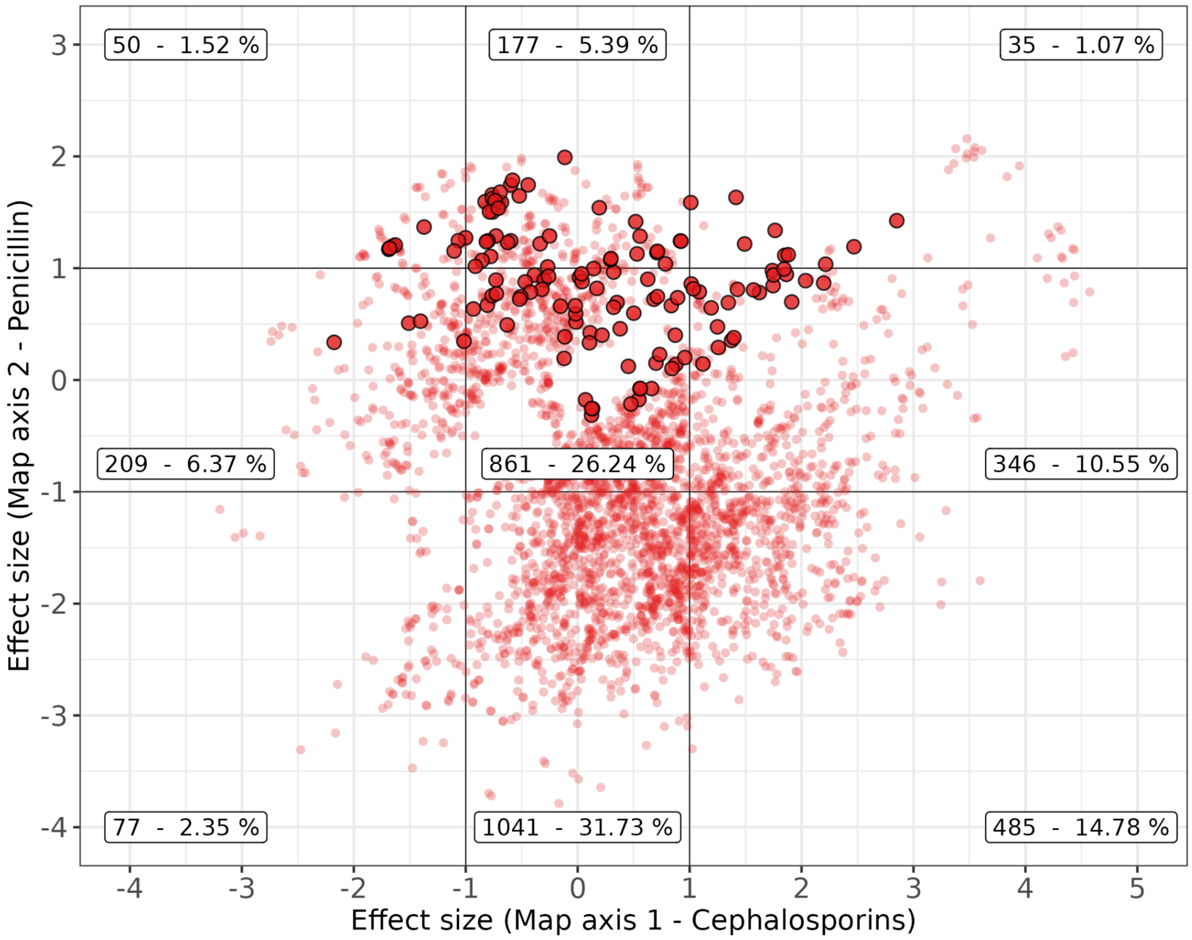


**Supplementary Figure S19.** Significant effect size estimates on map axis one and two as estimated by mvLMMs. Solid points represent amino acid substitutions, while smaller transparent points represent an epistatic interaction between two substitutions. Black lines represent reference lines of one $\log_2$ MIC unit, and show the categories of negative, neutral and positive effects for each map axis. Labels indicate the number and percentage of significant effects in each category.

2. *PBP substitutions identified using multivariate methods show strong overlap with previously published work*

The PBP substitutions identified here show strong overlap with the 72 positions previously identified by GWAS, random forest/cut-offs and laboratory methods (Supplementary Figure S20). Notably, 12 positions found to cause phenotypic effects in laboratory methods were identified using cartography, but not using previous GWAS or random forest/threshold methods. In contrast, a further 21 positions were found to cause phenotypic effects by GWAS (n = 10), and *in vitro* work (n = 11) but were not identified here. However, many of these positions were either invariant, or the relevant substitutions had very low frequency (<1%) within this dataset, making it not possible to test them here. The cartography methods also identified 128 positions not highlighted by previous methods, with many of those identified having contrasting effects on subclasses of drugs, or epistatic interactions with other positions.

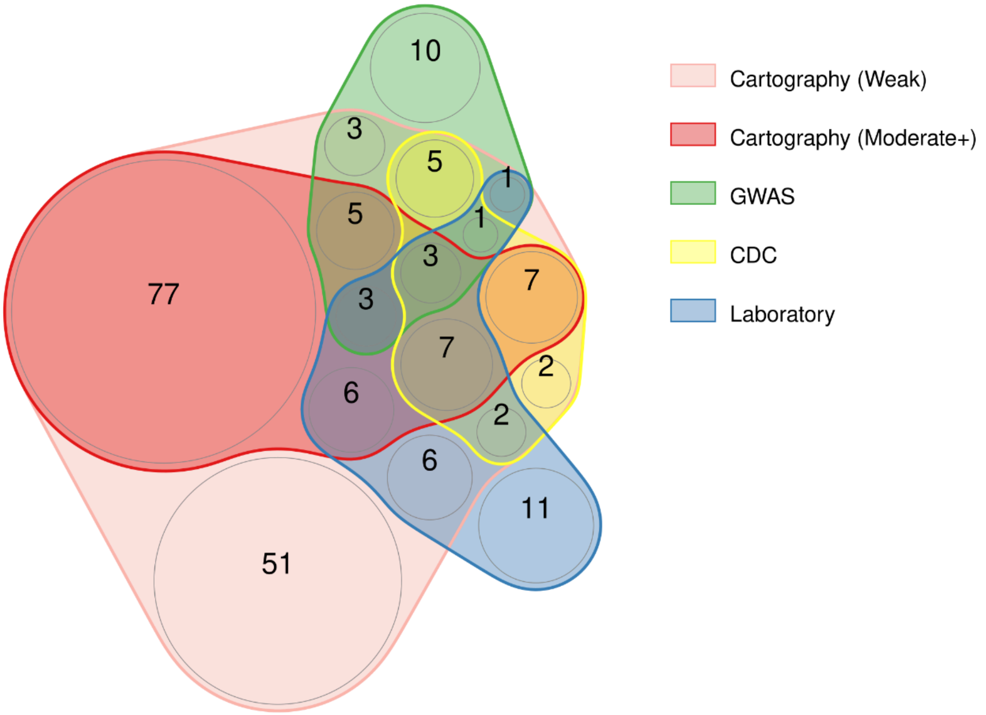


**Supplementary Figure S20.** Overlap between key PBP positions identified using cartography methods and those identified previously**.** Previous methods used GWAS, random forest or *in vitro* experiments to identify PBP substitutions associated with change in beta-lactam MIC (2,3,13,24). "CDC" refers to PBP positions reported using random forest and threshold cutoff methods on the same CDC Active Bacterial Core surveillance pneumococcal dataset presented here (2,3). Here, all positions identified by the cartography method are shown, where those with weak evidence are highlighted in pink, and those with moderate evidence or above are highlighted in red. References for *in vitro* work were taken from a review of laboratory work on PBP proteins in *S. pneumoniae* (reviewed in (13), references: (25–41)).

3. *Multivariate methods identify additional associations compared to univariate counterparts*

We tested whether the mvLMM on MDS coordinates offered increased statistical power compared with univariate LMMs conducted on each drug individually (Supplementary File 1). After compound markers were split into their individual amino acid changes, 178 candidate substitutions were identified by either analysis at an adjusted p-value threshold of 0.001: 53 by both methods, 123 by the multivariate analysis only, and 2 by the univariate analysis only. The 123 substitutions identified only by the multivariate analysis occurred at 107 PBP positions. Eleven of these substitutions were also supported by the clustering analysis, although none met the single-substitution effect threshold. Notably, 81 of the 123 multivariate-only substitutions (66%) had effects of opposing sign on the two map axes, compared with 5 of the 53 substitutions (9%) detected by both methods. One example was PBP2B L609S, which had negative estimated effects on cephalosporin and meropenem MIC but a positive estimated effect on penicillin MIC. Some substitutions identified by the mvLMM also showed nominal evidence in the uvLMM ($p < 0.05$) but did not pass the adjusted threshold of $p \leq 0.001$. Two marker variables were detected only by the uvLMM. PBP2B N656X represents a dataset-specific ambiguous amino-acid call rather than a genuine substitution and had no support from the other analyses. PBP2X T338A, in contrast, is a previously characterised resistance-associated substitution within the S337TMK active-site motif. It did not pass the mvLMM comparison threshold (adjusted p = 0.031; joint effect size = 0.956) but was involved in 52 supported epistatic interactions.

**References:**


1. Schuchat A, Hilger T, Zell E, Farley MM, Reingold A, Harrison L, et al. Active bacterial core surveillance of the emerging infections program network. Emerg Infect Dis. 2001 Jan;7(1):92–9. doi:10.3201/eid0701.010114 PubMed PMID: 11266299; PubMed Central PMCID: PMC2631675.

2. Li Y, Metcalf BJ, Chochua S, Li Z, Gertz RE, Walker H, et al. Validation of β-lactam minimum inhibitory concentration predictions for pneumococcal isolates with newly encountered penicillin binding protein (PBP) sequences. BMC Genomics. 2017 Aug 15;18(1):621. doi:10.1186/s12864-017-4017-7 PubMed PMID: 28810827.

3. Li Y, Metcalf BJ, Chochua S, Li Z, Gertz RE, Walker H, et al. Penicillin-Binding Protein Transpeptidase Signatures for Tracking and Predicting β-Lactam Resistance Levels in Streptococcus pneumoniae. mBio. 2016 Jun 14;7(3):10.1128/mbio.00756-16. doi:10.1128/mbio.00756-16

4. Smith DJ, Lapedes AS, de Jong JC, Bestebroer TM, Rimmelzwaan GF, Osterhaus ADME, et al. Mapping the antigenic and genetic evolution of influenza virus. Science. 2004 Jul 16;305(5682):371–6. doi:10.1126/science.1097211 PubMed PMID: 15218094.

5. Katzelnick LC, Fonville JM, Gromowski GD, Bustos Arriaga J, Green A, James SL, et al. Dengue viruses cluster antigenically but not as discrete serotypes. Science. 2015 Sep 18;349(6254):1338–43. doi:10.1126/science.aac5017 PubMed PMID: 26383952; PubMed Central PMCID: PMC4876809.

6. Mair P, Groenen PJF, De Leeuw J. More on Multidimensional Scaling and Unfolding in R: smacof Version 2. J Stat Softw. 2022 May 13;102:1–47. doi:10.18637/jss.v102.i10

7. Leeuw JD, Mair P, Org E, De Leeuw J. Multidimensional scaling using majorization: SMACOF in R. J Stat Softw. 2011 Oct 25;31:1–30. doi:10.18637/jss.v031.i03

8. Mair P, Borg I, Rusch T. Goodness-of-Fit Assessment in Multidimensional Scaling and Unfolding. Multivar Behav Res. 2016;51(6):772–89. doi:10.1080/00273171.2016.1235966 PubMed PMID: 27802073.

9. Graffelman J. A Guide to Scatterplot and Biplot Calibration.

10. Michael A, Kelman T, Pitesky M. Overview of Quantitative Methodologies to Understand Antimicrobial Resistance via Minimum Inhibitory Concentration. Anim Open Access J MDPI. 2020 Aug 12;10(8):1405. doi:10.3390/ani10081405 PubMed PMID: 32806615; PubMed Central PMCID: PMC7459578.

11. Andrews JM. Determination of minimum inhibitory concentrations. J Antimicrob Chemother. 2001 Jul 1;48(suppl_1):5–16. doi:10.1093/jac/48.suppl_1.5

12. Pattinson DJ. Predicting the Antigenic Evolution of Influenza Viruses with Application to Vaccination Strategy [Internet]. [Cambridge, UK]: University of Cambridge; 2020 [cited 2025 Aug 15]. Available from: https://www.repository.cam.ac.uk/items/0fd9bca6-c64b-47cd-92b4-34302f5e6bd6

13. Hakenbeck R, Brückner R, Denapaite D, Maurer P. Molecular mechanisms of β-lactam resistance in Streptococcus pneumoniae. Future Microbiol. 2012 Mar 6;7(3):395–410. doi:10.2217/fmb.12.2 PubMed PMID: 22393892.

14. Lynch M, Walsh B. Genetics and analysis of quantitative traits [Internet]. Vol. 1. Sinauer Sunderland, MA; 1998 [cited 2025 Sep 8]. Available from: https://www.invemar.org.co/redcostera1/invemar/docs/RinconLiterario/2011/febrero/AG_8.pdf

15. Kruijer W, Boer MP, Malosetti M, Flood PJ, Engel B, Kooke R, et al. Marker-Based Estimation of Heritability in Immortal Populations. Genetics. 2015 Feb;199(2):379–98. doi:10.1534/genetics.114.167916 PubMed PMID: 25527288; PubMed Central PMCID: PMC4317649.

16. Yang J, Manolio TA, Pasquale LR, Boerwinkle E, Caporaso N, Cunningham JM, et al. Genome partitioning of genetic variation for complex traits using common SNPs. Nat Genet. 2011;43(6):519–25. doi:10.1038/NG.823 PubMed PMID: 21552263.

17. Lippert C, Casale FP, Rakitsch B, Stegle O. LIMIX: genetic analysis of multiple traits. bioRxiv. 2014 May 22;003905. doi:10.1101/003905

18. Lippert C, Listgarten J, Liu Y, Kadie CM, Davidson RI, Heckerman D. FaST linear mixed models for genome-wide association studies. Nat Methods. 2011 Oct;8(10):833–5. doi:10.1038/nmeth.1681

19. Pensar J, Puranen S, Arnold B, MacAlasdair N, Kuronen J, Tonkin-Hill G, et al. Genome-wide epistasis and co-selection study using mutual information. Nucleic Acids Res. 2019 Oct 10;47(18):e112–e112. doi:10.1093/NAR/GKZ656 PubMed PMID: 31361894.

20. Skwark MJ, Croucher NJ, Puranen S, Chewapreecha C, Pesonen M, Xu YY, et al. Interacting networks of resistance, virulence and core machinery genes identified by genome-wide epistasis analysis. PLoS Genet. 2017;13(2):e1006508. doi:10.1371/journal.pgen.1006508

21. Puranen S, Pesonen M, Pensar J, Xu YY, Lees JA, Bentley SD, et al. SuperDCA for genome-wide epistasis analysis. Microb Genomics. 2018 Jun 1;4(6). doi:10.1099/MGEN.0.000184 PubMed PMID: 29813016.

23. Galwey NW. A new measure of the effective number of tests, a practical tool for comparing families of non-independent significance tests. Genet Epidemiol. 2009 Nov;33(7):559–68. doi:10.1002/GEPI.20408 PubMed PMID: 19217024.

24. Chewapreecha C, Marttinen P, Croucher NJ, Salter SJ, Harris SR, Mather AE, et al. Comprehensive identification of single nucleotide polymorphisms associated with beta-lactam resistance within pneumococcal mosaic genes. PLoS Genet. 2014;10(8):e1004547. doi:10.1371/journal.pgen.1004547

25. Kocaoglu O, Tsui HCT, Winkler ME, Carlson EE. Profiling of β-lactam selectivity for penicillin-binding proteins in Streptococcus pneumoniae D39. Antimicrob Agents Chemother. 2015 Jun 1;59(6):3548–55. doi:10.1128/AAC.05142-14

26. Coffey TJ, Daniels M, McDougal LK, Dowson CG, Tenover FC, Spratt BG. Genetic analysis of clinical isolates of Streptococcus pneumoniae with high-level resistance to expanded-spectrum cephalosporins. Antimicrob Agents Chemother. 1995;39(6):1306–13. doi:10.1128/AAC.39.6.1306 PubMed PMID: 7574521.

27. Hakenbeck R, Grebe T, Zähner D, Stock JB. β-lactam resistance in Streptococcus pneumoniae: Penicillin-binding proteins and non-penicillin-binding proteins. Mol

Microbiol. 1999 Aug 1;33(4):673–8. doi:10.1046/j.1365-2958.1999.01521.x PubMed PMID: 10447877.

28. Hakenbeck R. Discovery of β-lactam-resistant variants in diverse pneumococcal populations. Genome Med. 2014 Sep 25;6(9):72. doi:10.1186/s13073-014-0072-8

29. Hsieh YC, Su LH, Hsu MH, Chiu CH, Cheng-Hsun Chiu C. Alterations of penicillin-binding proteins in pneumococci with stepwise increase in beta-lactam resistance. Pathog Dis. 2013;67(1):84–8. doi:10.1111/2049-632x.12018

30. Dowson CG, Coffey TJ, Kell C, Whiley RA. Evolution of penicillin resistance in Streptococcus pneumoniae; the role of Streptococcus mitis in the formation of a low affinity PBP2B in S. pneumoniae. Mol Microbiol. 1993;9(3):635–43. doi:10.1111/J.1365-2958.1993.TB01723.X PubMed PMID: 8412708.

31. Nagai K, Davies TA, Jacobs MR, Appelbaum PC. Effects of amino acid alterations in penicillin-binding proteins (PBPs) 1a, 2b, and 2x on PBP affinities of penicillin, ampicillin, amoxicillin, cefditoren, cefuroxime, cefprozil, and cefaclor in 18 clinical isolates of penicillin-susceptible, -intermedia. Antimicrob Agents Chemother. 2002;46(5):1273–80. doi:10.1128/AAC.46.5.1273-1280.2002 PubMed PMID: 11959556.

32. Laible G, Spratt BG, Hakenbeck R. Interspecies recombinational events during the evolution of altered PBP 2x genes in penicillin-resistant clinical isolates of Streptococcus pneumoniae. Mol Microbiol. 1991;5(8):1993–2002. doi:10.1111/J.1365-2958.1991.TB00821.X PubMed PMID: 1766375.

33. Pernot L, Chesnel L, Gouellec AL, Croizé J, Vernet T, Dideberg O, et al. A PBP2x from a clinical isolate of Streptococcus pneumoniae exhibits an alternative mechanism for reduction of susceptibility to β-lactam antibiotics. J Biol Chem. 2004 Apr 16;279(16):16463–70. doi:10.1074/JBC.M313492200

34. Lalble G, Hakenbeck R. Penicillin-binding proteins in beta-lactam-resistant laboratory mutants of Streptococcus pneumoniae. Mol Microbiol. 1987;1(3):355–63. doi:10.1111/J.1365-2958.1987.TB01942.X PubMed PMID: 3448464.

35. Sauerbier J, Maurer P, Rieger M, Hakenbeck R. Streptococcus pneumoniae R6 interspecies transformation: genetic analysis of penicillin resistance determinants and genome-wide recombination events. Mol Microbiol. 2012 Nov 1;86(3):692–706. doi:10.1111/MMI.12009

36. Carapito R, Chesnel L, Vernet T, Zapun A. Pneumococcal beta-lactam resistance due to a conformational change in penicillin-binding protein 2x. J Biol Chem. 2006 Jan 20;281(3):1771–7. doi:10.1074/JBC.M511506200 PubMed PMID: 16303769.

37. Job V, Carapito R, Vernet T, Dessen A, Zapun A. Common alterations in PBP1a from resistant Streptococcus pneumoniae decrease Its reactivity toward β-lactams: structural insights. J Biol Chem. 2008 Feb 22;283(8):4886–94. doi:10.1074/JBC.M706181200 PubMed PMID: 18055459.

38. Mouz N, Di Guilmi AM, Gordon E, Hakenbeck R, Dideberg O, Vernet T. Mutations in the active site of penicillin-binding protein PBP2x from Streptococcus pneumoniae. Role in the specificity for beta-lactam antibiotics. J Biol Chem. 1999 Jul 2;274(27):19175–80. doi:10.1074/JBC.274.27.19175 PubMed PMID: 10383423.

39. Zerfaß I, Hakenbeck R, Denapaite D. An important site in PBP2x of penicillin-resistant

clinical isolates of Streptococcus pneumoniae: Mutational analysis of Thr338. Antimicrob Agents Chemother. 2009 Mar;53(3):1107–15. doi:10.1128/AAC.01107-08/ PubMed PMID: 19075056.

40. Grebe T, Hakenbeck R. Penicillin-binding proteins 2b and 2x of Streptococcus pneumoniae are primary resistance determinants for difierent classes of β-lactam antibiotics. Antimicrob Agents Chemother. 1996;40(4):829–34. doi:10.1128/AAC.40.4.829

41. Philippe J, Gallet B, Morlot C, Denapaite D, Hakenbeck R, Chen Y, et al. Mechanism of β-lactam action in Streptococcus pneumoniae: The piperacillin paradox. Antimicrob Agents Chemother. 2015 Jan 1;59(1):609–21. doi:10.1128/AAC.04283-14/ PubMed PMID: 25385114.
